# Natural variation in *C. elegans* arsenic toxicity is explained by differences in branched chain amino acid metabolism

**DOI:** 10.1101/373787

**Authors:** Stefan Zdraljevic, Bennett W. Fox, Christine Strand, Oishika Panda, Francisco J. Tenjo, Shannon C. Brady, Tim A. Crombie, John G. Doench, Frank C. Schroeder, Erik C. Andersen

## Abstract

We find that variation in the *dbt-1* gene underlies natural differences in *Caenorhabditis elegans* responses to the toxin arsenic. This gene encodes the E2 subunit of the branched-chain α-keto acid dehydrogenase (BCKDH) complex, a core component of branched-chain amino acid (BCAA) metabolism. We causally linked a non-synonymous variant in the conserved lipoyl domain of DBT-1 to differential arsenic responses. Using targeted metabolomics and chemical supplementation, we demonstrate that differences in responses to arsenic are caused by variation in iso-branched chain fatty acids. Additionally, we show that levels of branched chain fatty acids in human cells are perturbed by arsenic treatment. This finding has broad implications for arsenic toxicity and for arsenic-focused chemotherapeutics across human populations. Our study implicates the BCKDH complex and BCAA metabolism in arsenic responses, demonstrating the power of *C. elegans* natural genetic diversity to identify novel mechanisms by which environmental toxins affect organismal physiology.

## Introduction

An estimated 100 million people are currently at risk of chronic exposure to arsenic, a toxic metalloid that can be found in the environment [1]. The high prevalence of environmental arsenic and the severe toxicity associated with exposure has made it the number one priority for the United States Agency for Toxic Substances and Disease Registry (https://www.atsdr.cdc.gov/SPL/). Inorganic trivalent arsenic As(III) compounds, which include arsenic trioxide (As_2_O_3_), are the most toxic forms of environmental arsenic [2,3]. In humans, As(III) is detoxified by consecutive methylation events, forming dimethylarsenite (DMA) [4,5]. However, this methylation process also creates the highly toxic monomethylarsenite (MMA) intermediate, so ratios of DMA to MMA determine levels of arsenic toxicity. Both MMA and DMA are produced from As(III) via the arsenic methyltransferase (AS3MT) [6]. Interestingly, individuals from human subpopulations that inhabit high arsenic environments have higher DMA/MMA ratios than individuals from low-arsenic environments. The elevated DMA/MMA ratio in these individuals is associated with natural differences in the *AS3MT* gene [7–9], which shows signs of strong positive selection. These results suggest that a more active AS3MT enzyme in these human subpopulations makes more DMA and enables adaptation to elevated environmental arsenic levels [6]. Importantly, population-wide differences in responses to environmental arsenic cannot be explained solely by variation in *AS3MT*, indicating that other genes must impact arsenic toxicity.

Despite its toxicity, arsenic trioxide has been used as a therapeutic agent for hundreds of years. Most recently, it was introduced as a highly effective cancer chemotherapeutic for the treatment of acute promyelocytic leukemia (APL) [10–13]. Hematopoietic differentiation and apoptosis in APL patients is blocked at the level of promyelocytes by the Promyelocytic Leukemia/Retinoic Acid Receptor alpha fusion protein caused by a t(15;17) chromosomal translocation [14,15]. Arsenic trioxide has been shown to directly bind a cysteine-rich region of the RING-B box coiled-coil domain of PML-RARα, which causes the degradation of the oncogenic fusion protein [16,17]. The success of arsenic trioxide (Trisenox®) has spurred its use in over a hundred clinical trials in the past decade [18]. Despite these successes, individual differences in the responses to arsenic-based treatments, including patient-specific dosing regimens and side effects, limit the full therapeutic benefit of this compound [19]. Medical practitioners require knowledge of the molecular mechanisms for how arsenic causes toxicity to provide the best individual-based therapeutic benefits.

Studies of the free-living roundworm *Caenorhabditis elegans* have greatly facilitated our understanding of basic cellular processes [20–23], including a number of studies that show that the effects of arsenic are similar to what is observed in mammalian model systems and humans. These effects include mitochondrial toxicity [24,25], the generation of reactive oxygen species (ROS) [26], genotoxicity [27], genome-wide shifts in chromatin structure [28], reduced lifespan [26], and the induction of the heat-shock response [29]. However, these studies were all performed in the genetic background of the standard *C. elegans* laboratory strain (N2). To date, 152 *C. elegans* strains have been isolated from various locations around the world [30–32], which contain a largely unexplored pool of genetic diversity much of which could underlie adaptive responses to environmental perturbations [33].

We used two quantitative genetic mapping approaches to show that a major source of variation in *C. elegans* responses to arsenic trioxide is caused by natural variation in the *dbt-1* gene, which encodes an essential component of the highly conserved branched-chain α-keto acid dehydrogenase (BCKDH) complex [34]. The BCKDH complex is a core component of branched-chain amino acid (BCAA) catabolism, which has not been previously implicated in arsenic responses. Furthermore, we show that a single missense variant in DBT-1(S78C), located in the highly conserved lipoyl-binding domain, underlies phenotypic variation in response to arsenic. Using targeted and untargeted metabolomics and chemical rescue experiments, we show that differences in wild isolate responses to arsenic trioxide are caused by differential synthesis of mono-methyl branched chain fatty acids (mmBCFA), metabolites with a central role in development [20]. These results demonstrate the power of using the natural genetic diversity across the *C. elegans* species to identify mechanisms by which environmental toxins affect physiology.

## Results

### Natural variation on chromosome II underlies differences in arsenic responses

We quantified arsenic trioxide sensitivity in *C. elegans* using a high-throughput fitness assay that utilizes the COPAS BIOSORT [35,36]. In this assay, three L4 larvae from each strain were sorted into arsenic trioxide or control conditions. After four days of growth, we quantified various attributes of populations that relate to the ability of *C. elegans* to grow in the presence of arsenic trioxide or control conditions (see Materials and Methods). To determine an appropriate concentration of arsenic trioxide for mapping experiments, we performed dose-response experiments on four genetically diverged isolates of *C. elegans*: N2, CB4856, JU775, and DL238 (Figure 1-source data 1). We focused on brood size and progeny length traits because they are the most direct measures of animal responses to environmental perturbations. We defined brood size as the total number of objects detected by the BIOSORT and progeny length by the median of populations of specific *C. elegans* strains. When compared to control conditions, all four strains produced fewer progeny at all arsenic trioxide concentrations, and the lowest concentration at which we observed a significant reduction in brood size for all strains was 1 mM (Figure 1-figure supplement 1A). We estimated broad-sense heritability (*H*^2^) for the brood size trait in 1 mM arsenic trioxide to be (0.65) (Figure 1-source data 2) and the strain effect to be 0.48 (partial omega squared, 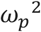, Figure 1-source data 3), indicating that this trait has a large genetic component and a large strain effect. In addition to reduced brood sizes, we observed that the progeny of animals exposed to arsenic trioxide were shorter in length than the progeny of animals grown in control conditions (Figure 1-figure supplement 1B), which indicates an arsenic-induced developmental delay (*H*^2^ = 0.15 - 0.49; Figure 1-source data 2 and 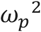 = 0.02 - 0.60; Figure 1-source data 3, depending on the animal length summary statistic used). We found that CB4856 animals produced approximately 16% more offspring that were on average 20% larger than the other three strains when treated with 1 mM arsenic trioxide, suggesting that the CB4856 strain was more resistant to arsenic trioxide than the other three strains. To reduce the number of traits for subsequent analyses, we performed principal component analysis on all measured traits, which transforms the data to orthogonal axes that capture the most phenotypic variance (Figure 1-source data 1 and 4-6; See Materials and Methods). For 1 mM arsenic trioxide, we estimated the broad-sense heritability (*H*^2^) of the first principal component to be 0.12 (Figure 1-source data 2) with an effect size of 0.16 (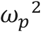) (Figure 1-source data 3). It is not surprising that the first principal component explained a large percentage (70%) of the total phenotypic variance because many of the summary statistic traits are highly correlated (Figure 1-figure supplement 2A-D; Figure 1-source data 7). We noted that the first principal component (PC1) was most strongly influenced by traits related to animal length, as indicated by the loadings (Figure 1-figure supplement 2A-D; Figure 1-source data 3 and 6), suggesting that PC1 is a biologically relevant trait (Figure 1-figure supplement 1C). Furthermore, because we observe a large range of effect sizes and broad-sense heritability estimates across measured traits (Figure 1-figure supplement 3), we focused our analyses on the PC1 trait for subsequent experiments.

**Figure 1:**
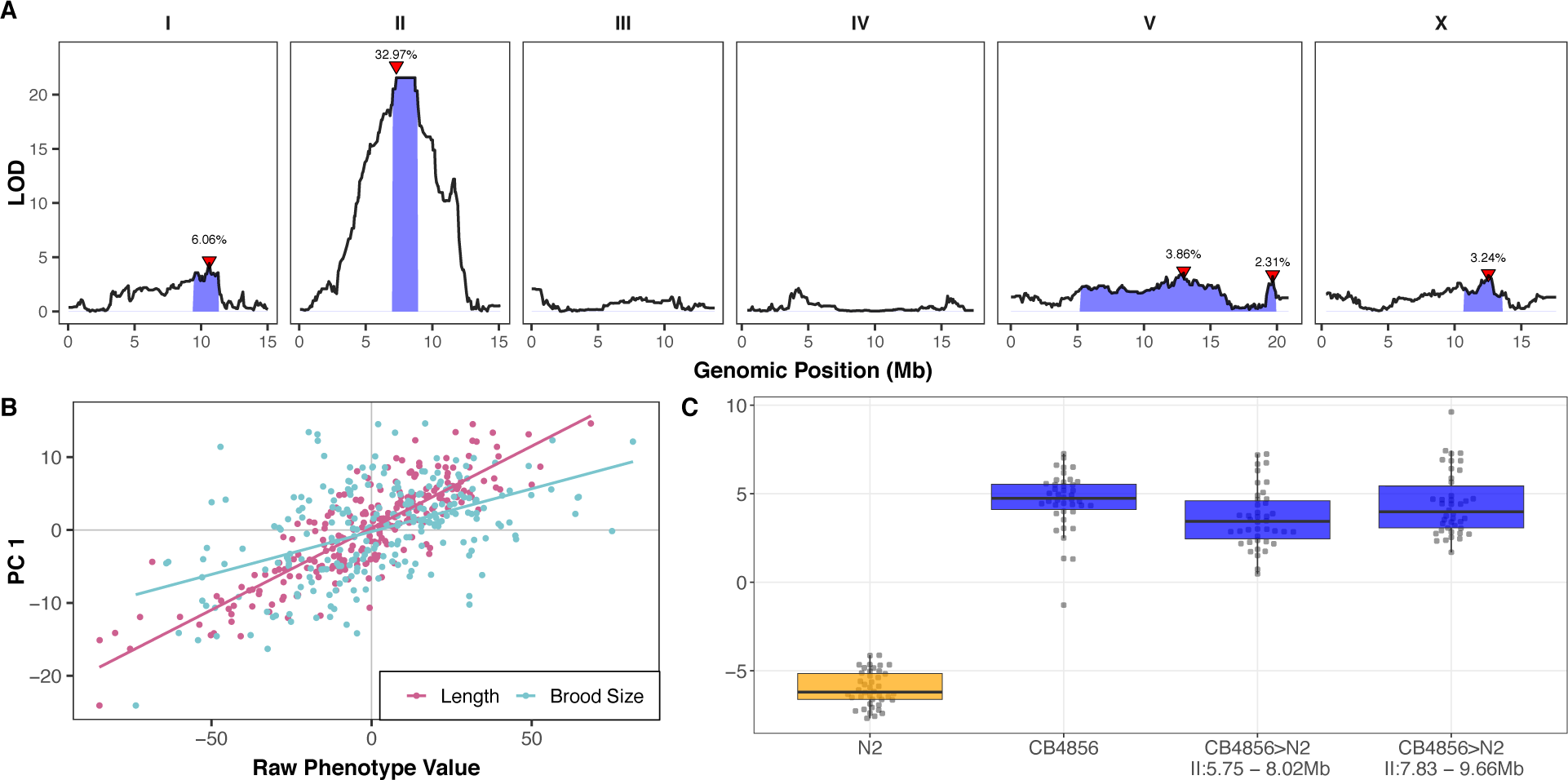
A large-effect QTL on the center of chromosome II explains differences in arsenic trioxide response between N2 and CB4856. (A) Linkage mapping plots for the first principal component trait in the presence of 1000 µM arsenic trioxide is shown. The significance values (logarithm of odds, LOD, ratio) for 1454 markers between the N2 and CB4856 strains are on the y-axis, and the genomic position (Mb) separated by chromosome is plotted on the x-axis. The associated 1.5 LOD-drop confidence intervals are represented by blue boxes. (B) The correlation between brood size (blue; *r*^2^ = 0.23, *p-*value = 2.4E-15) or animal length (pink; *r*^2^ = 0.72, *p-*value = 7E-70) with the first principal component trait. Each dot represents an individual RIAIL’s phenotype, with the animal length and brood size phenotype values on the x-axis and the first principal component phenotype on the y-axis. (C) Tukey box plots of near-isogenic line (NIL) phenotype values for the first principal component trait in the presence of 1000 µM arsenic trioxide is shown. NIL genotypes are indicated below the plot as genomic ranges. The N2 trait is significantly different than the CB4856 and NIL traits (Tukey HSD *p*-value < 1E-5).

The increased arsenic trioxide resistance of CB4856 compared to N2 motivated us to perform linkage mapping experiments with a panel of recombinant inbred advanced intercross lines (RIAILs) that were previously constructed through ten generations of intercrossing between an N2 derivative (QX1430) and CB4856 [35]. To capture arsenic trioxide-induced phenotypic differences, we exposed a panel of 252 RIAILs to 1 mM arsenic trioxide and corrected for growth differences among RIAILs in control conditions and assay-to-assay variability using linear regression (Figure 1-source data 8; see Materials and Methods). We performed linkage mapping on processed traits and the eigenvector-transformed traits (principal components or PCs) obtained from PCA that explained 90% of the variance in the processed trait set (Materials and Methods). The rationale of this approach was to minimize trait fluctuations that could be caused by only measuring the phenotypes of one replicate per RIAIL strain. In agreement with our observations from the dose-response experiment, we found that PC1 captures 70.9% of the total measured trait variance and is strongly influenced by correlated animal length traits (Figure 1-figure supplement 4; Figure 1-source data 9-10). Linkage mapping analysis of the PC1 trait revealed that arsenic trioxide-induced phenotypic variation is significantly associated with genetic variation on the center of chromosome II (Figure 1A-figure supplement 5; Figure 1-source data 11). An additional four quantitative trait loci (QTL) were significantly associated with variation in arsenic responses on chromosomes I, V, and X (Figure 1A-figure supplement 5). Consistent with the loadings of PC1, we determined that PC1 is highly correlated with the both brood size and animal length traits (Figure 1B), suggesting that PC1 captures RIAIL variation in these traits. To further support this relationship to interpretable biological significance, we found that the animal length and brood size traits map to the same region on the center of chromosome II (Figure 1-figure supplement 6; Figure 1-source data 11). The QTL on the center of chromosome II explains 32.97% of the total RIAIL phenotypic variation for the PC1 trait, which accounts for 61.7% of the total phenotypic variation that can be explained by genetic factors (*H*^2^ = 0.53) (Figure 1-figure supplement 7; Figure 1-source data 12). Taken together, the five QTL identified by mapping the PC1 trait account for 48.4% of the total RIAIL variation, corresponding to 91.3% of the total phenotypic variation that can be explained by genetic factors. However, we did not account for errors in genomic heritability estimates. In addition to the PC1, brood size, and animal length traits, 69 additional measured traits map to the same genomic region (Figure 1-figure supplement 8; Figure 1-source data 11). This result was expected given correlation structure of the measured traits (Figure 1-figure supplement 4). The PC1 QTL confidence interval spans from 7.04 to 8.87 Mb on chromosome II. This QTL is completely encompassed by the brood size (6.06-9.30 Mb) and animal length (6.82-8.93 Mb) QTL confidence intervals (Figure 1-figure supplement 6). However, each of these QTL confidence intervals span genomic regions greater than 1.5 megabases and contain hundreds of genes that vary between the N2 and CB4856 strains. Therefore, we constructed near-isogenic lines (NILs) to isolate and narrow each QTL in a controlled genetic background.

To split the large QTL confidence intervals in half, we introgressed genomic regions from the CB4856 strain on the left and right halves of the confidence intervals into the N2 genetic background. In the presence of arsenic trioxide, both of these NILs recapitulated the parental CB4856 PC1 phenotype (Figure 1C; Figure 1-source data 13), and had similar brood sizes and progeny of similar length as the CB4856 parental strain (Figure 1-figure supplement 9; Figure 1-source data 13). Furthermore, we show that similar to the RIAIL phenotypes, many of the measured traits were highly correlated (Figure 1-figure supplement 10A; Figure 1-source data 14) and contributed similarly to the PC1 trait (Figure 1-figure supplement 10B; Figure 1-source data 15). Importantly, the PC1 trait was highly correlated with the brood size and animal length phenotypes (Figure 1-figure supplement 11). The phenotypic similarity of these NILs to the CB4856 parental strain suggested that the two NILs might share an introgressed region of the CB4856 genome. To identify this shared introgressed region, we performed low-coverage whole-genome sequencing of the NIL strains and defined the left and right bounds of the CB4856 genomic introgression to be from 5.75 to 8.02 Mb and 7.83 to 9.66 Mb in the left and right NILs, respectively (Figure 1-source data 16). The left and right NILs recapitulate 88.6% and 91.6% of the effect size difference between N2 and CB4856 as measured by Cohen’s F, respectively [37], which exceeds our observations the linkage mapping results where the QTL on chromosome II explained 61.7% of the total phenotypic variation in the RIAIL population. This discrepancy was observed likely because the NILs are a more homogenous genetic background and the experiment was performed at higher replication than the linkage mapping. Taken together, these results suggested that genetic differences between N2 and CB4856 within 7.83 to 8.02 Mb on chromosome II conferred resistance to arsenic trioxide.

In parallel to the linkage-mapping approach described above, we performed a genome-wide association (GWA) mapping experiment by quantifying the responses to arsenic trioxide for 86 wild *C. elegans* strains (Figure 2-source data 1) [30]. Consistent with previous experiments, the measured traits that most strongly influence PC1 are representative of animal length, as indicated by the loadings (Figure 2-figure supplement 1; Figure 2-source data 2–3). In agreement with the results from the linkage mapping approach, PC1 differences among the wild isolates mapped to a QTL on the center of chromosome II that spans from 7.71 Mb to 8.18 Mb (Figure 2A; Figure 2-source data 4-5). We noted that the brood size trait did not map to a significant QTL with the GWA mapping approach. To address this discrepancy, we estimated genomic broad- and narrow-sense heritability (*H*^2^; *h*^2^) for all of the wild isolate measured and principal component traits (Figure 2-figure supplement 2; Figure 2-source data 6). Broad-sense heritability for the brood size trait was approximately 0.05 for the wild isolates tested, which explains why we were unable to detect a significant QTL associated with this trait. By contrast, the PC1 trait *H*^2^ was 0.16 when estimated using an additive-only model and 0.25 (*h*^2^ = 0.11) when we incorporated the possibility for epistatisis (See Materials and Methods; Figure 2-source data 6). The marker found to be most correlated with the PC1 trait from GWA mapping (II:7,931,252), explains 84.6% (epistasis model) of the total heritable phenotypic variation. Of the 64 measured and principal component traits we mapped, 26 were significantly correlated with genetic variation on the center of chromosome II (Figure 2-figure supplement 3; Figure 2-source data 7). Notably, the CB4856 strain, which was one of the parents used to construct the RIAIL panel used for linkage mapping, had the non-reference genotype at the marker most correlated with PC1 (Figure 2B), suggesting that the same genetic variant(s) might be contributing to differential arsenic trioxide response among the RIAIL and wild isolate populations.

**Figure 2:**
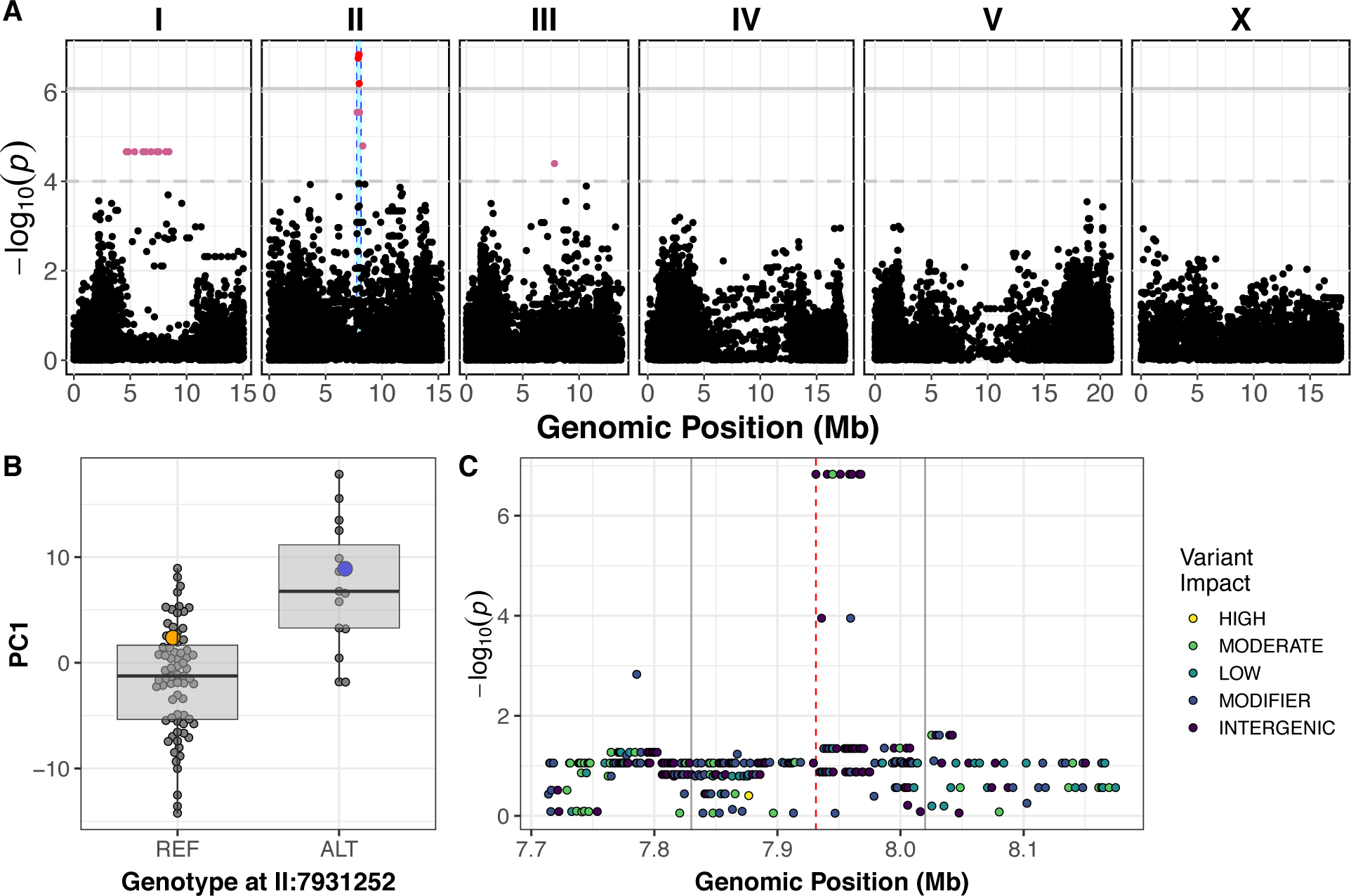
Variation in *C. elegans* wild isolates responses to arsenic trioxide maps to the center of chromosome II. (A) A manhattan plot for the first principal component in the presence of 1000 µM arsenic trioxide is shown. Each dot represents an SNV that is present in at least 5% of the assayed wild population. The genomic position in Mb, separated by chromosome, is plotted on the x-axis and the −*log*_10_(*p*) for each SNV is plotted on the y-axis. SNVs are colored red if they pass the genome-wide Bonferroni-corrected significance threshold, which is denoted by the gray horizontal line. SNVs are colored pink if they pass the genome-wide eigen-decomposition significance threshold, which is denoted by the dotted gray horizontal line. The genomic region of interest surrounding the QTL is represented by a cyan rectangle. (B) Tukey box plots of phenotypes used for association mapping in (A) are shown. Each dot corresponds to the phenotype of an individual strain, which is plotted on the y-axis. Strains are grouped by their genotype at the peak QTL position (red SNV from panel A, ChrII:7,931,252), where REF corresponds to the allele from the reference N2 strain. The N2 (orange) and CB4856 (blue) strains are highlighted. (C) Fine mapping of the chromosome II region of interest (cyan region from panel A, 7.71 - 8.18 Mb) is shown. Each dot represents an SNV present in the CB4856 strain. The association between the SNV and first principal component is shown on the y-axis and the genomic position of the SNV is shown on the x-axis. Dots are colored by their SnpEff predicted effect.

To fine map the PC1 QTL, we focused on variants from the *C. elegans* whole-genome variation dataset [31] that are shared among at least five percent of the 86 wild isolates exposed to arsenic trioxide. Under the assumption that the linkage and GWA mapping QTL are caused by the same genetic variation, we only considered variants present in the CB4856 strain. Eight markers within the QTL region are in complete linkage disequilibrium with each other and are most correlated with the first principal component trait (Figure 2C-figure supplement 4; Figure 2-source data 8). Only one of these markers is located within an annotated gene (*dbt-1*) and is predicted to encode a cysteine-to-serine variant at position 78 (C78S). Although it is possible that the causal variant underlying differential arsenic trioxide response in the *C. elegans* population is an intergenic variant, we focused on the DBT-1(C78S) variant as a candidate to test for an effect on arsenic response.

### A cysteine-to-serine variant in DBT-1 contributes to arsenic response variation

The *C. elegans dbt-1* gene encodes the E2 component of the branched-chain α-keto acid dehydrogenase complex (BCKDH) [34]. The BCKDH complex is a core component of branched-chain amino acid (BCAA) catabolism and catalyzes the irreversible oxidative decarboxylation of amino acid-derived branched-chain α-ketoacids [38]. The BCKDH complex belongs to a family of α-ketoacid dehydrogenases that include pyruvate dehydrogenase (PDH) and α-ketoglutarate dehydrogenase (KGDH) [39]. All three of these large enzymatic complexes include a central E2 component that is lipoylated at one critical lysine residue (two residues in PDH). The function of these enzymatic complexes depends on the lipoylation of these lysine residues [39,40]. In *C. elegans*, the putative lipoylated lysine residue is located at amino acid position 71 of DBT-1, which is in close proximity to the C78S residue that we found to be highly correlated with arsenic trioxide resistance.

To confirm that the C78S variant in DBT-1 contributes to differential arsenic trioxide responses, we used CRISPR/Cas9-mediated genome editing to generate allele-replacement strains by changing the C78 residue in the N2 strain to a serine and the S78 residue in the CB4856 strain to a cysteine. When treated with arsenic trioxide, the N2 DBT-1(S78) allele-replacement strain recapitulated 75.5% of the phenotypic difference between the CB4856 and N2 strains as measured with the first principal component, as measured by Cohen’s F [37] (Figure 3; Figure 1-source data 13). Similarly, the CB4856 DBT-1(C78) allele-replacement strain recapitulated 63.9% of the total phenotypic difference between the two parental CB4856. The degree to which the allele-replacement strains recapitulated the difference in the first principal component between the N2 and CB4856 strains matched our observations from the linkage mapping experiment, where the chromosome II QTL explained 61.7% of the total phenotypic variation in the RIAIL population. This result suggested that the majority of heritable variation in arsenic trioxide response was explained by the DBT-1(C78S) allele. We obtained similar results for the progeny length trait (Figure 3-figure supplement 1; Figure 1-source data 13), which is likely because of the high level of correlation between the animal length and first principal component trait (Figure 1-figure supplement 10; Figure 3-figure supplement 2; Figure 1-source data 10–11). However, when considering brood size, the N2 DBT-1(C78S) allele-replacement strain produced an intermediate number of progeny in the presence of arsenic trioxide relative to the parental N2 and CB4856 strains. And the CB4856 DBT-1(S78C) allele-replacement strain produced fewer offspring than both parental strains (Figure 3-figure supplement 1; Figure 1-source data 13). These results suggested that additional genetic variants between the N2 and CB4856 strains might interact with the DBT-1(C78S) allele to affect different aspects of physiology. Nevertheless, these results functionally validated that the DBT-1 C78S variant underlies differences in physiological responses to arsenic trioxide.

**Figure 3:**
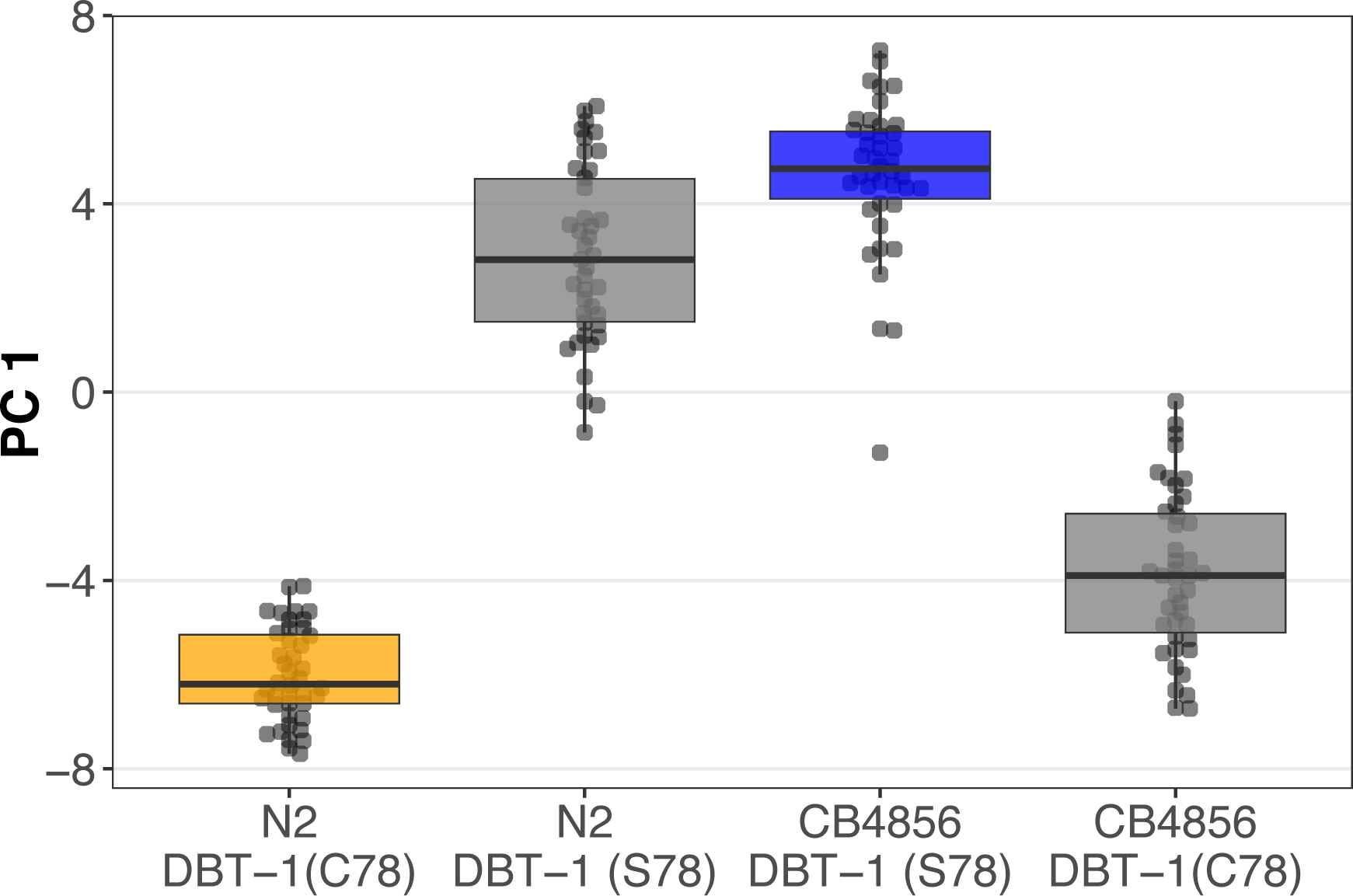
The DBT-1(C78S) variant contributes to arsenic trioxide responses. Tukey box plots of the first principal component generated by PCA on allele-replacement strain phenotypes measured by the COPAS BIOSORT 1000 µM arsenic trioxide exposure are shown (N2, orange; CB4856, blue; allele replacement strains, gray). Labels correspond to the genetic background and the corresponding residue at position 78 of DBT-1 (C for cysteine, S for serine). All pair-wise comparisons are significantly different (Tukey HSD, *p-*value < 1.42E-5).

### Arsenic trioxide inhibits the DBT-1 C78 allele

Mono-methyl branched chain fatty acids (mmBCFA) are an important class of molecules that are produced via BCAA catabolism [20,34,41,42]. The production of mmBCFA requires the BCKDH, fatty acid synthase (FASN-1), acetyl-CoA carboxylase (POD-2), fatty acyl elongases (ELO-5/6), β-ketoacyl dehydratase (LET-767), and acyl CoA synthetase (ACS-1) [20,34,41,43–45]. Strains that lack functional *elo-5*, *elo-6*, or *dbt-1* produce less C15ISO and C17ISO mmBCFAs, arrest at the L1 larval stage, and can be rescued by supplementing the growth media with C15ISO or C17ISO [20,34,41] (Figure 4A).

**Figure 4:**
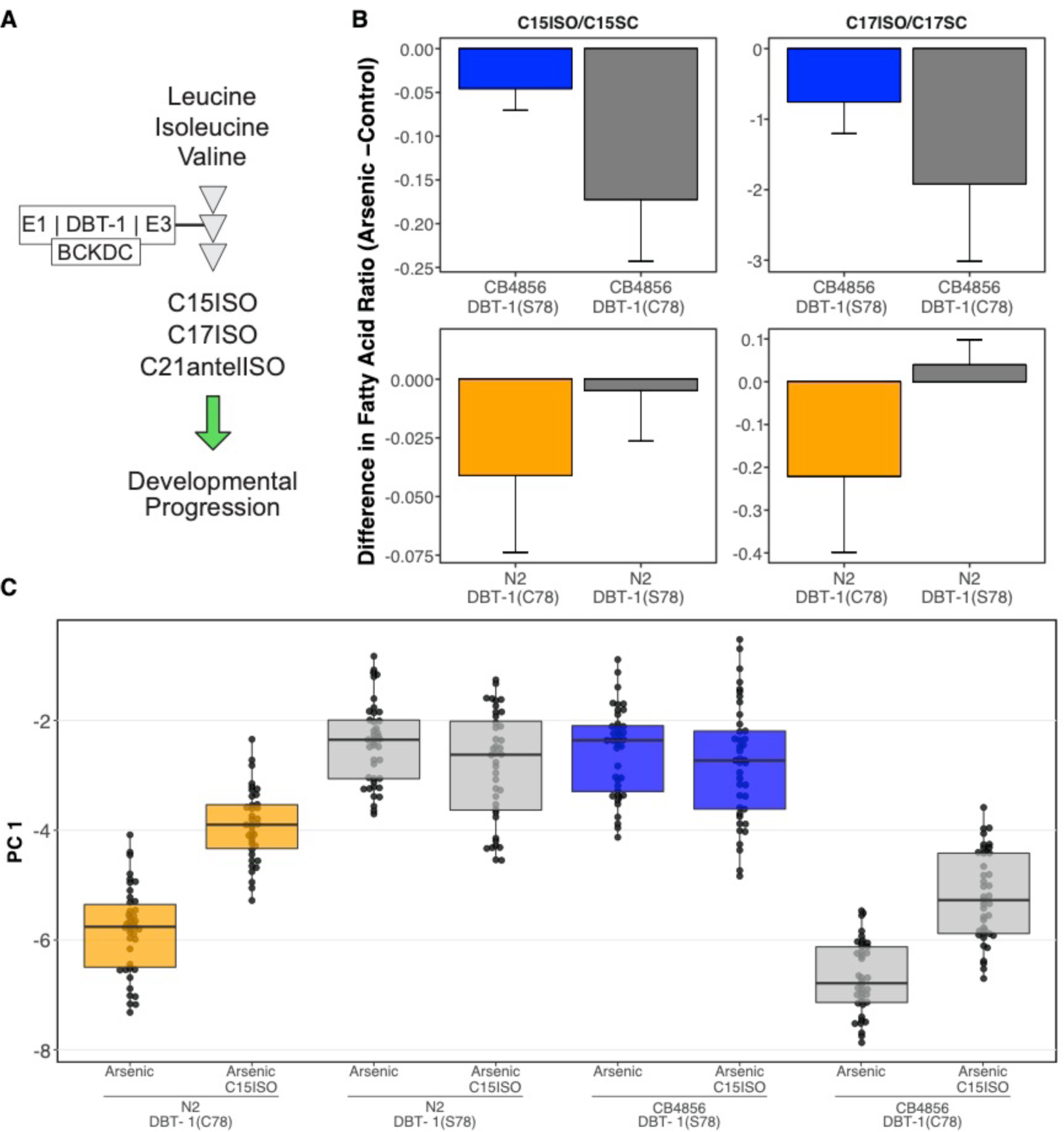
Differential production of mmBCFA underlies DBT-1(C78)-mediated sensitivity to arsenic trioxide. (A) A simplified model of BCAA catabolism in *C. elegans*. The BCKDH complex, which consists of DBT-1, catalyzes the irreversible oxidative decarboxylation of branched-chain ketoacids. The products of these breakdown can then serve as building blocks for the mmBCFA that are required for developmental progression. (B) The difference in the C15ISO/C15SC (left panel) or C17ISO/C17SC (right panel) ratios between 100 µM arsenic trioxide and control conditions is plotted on the y-axis for three independent replicates of the CB4856 and CB4856 allele swap strains and six independent replicates of the N2 and N2 allele swap strains. The difference between the C15 ratio for the CB4856-CB4856 allele swap comparison is significant (Tukey HSD *p-*value = 0.0427733), but the difference between the C17 ratios for these two strains is not (Tukey HSD *p-*value = 0.164721). The difference between the C15 and C17 ratios for the N2-N2 allele swap comparisons are both significant (C15: Tukey HSD *p-*value = 0.0358; C17: Tukey HSD *p-* value = 0.003747). (C) Tukey box plots median animal length after arsenic trioxide or arsenic trioxide and 0.64 µM C15ISO exposure are shown (N2, orange; CB4856, blue; allele replacement strains, gray). Labels correspond to the genetic background and the corresponding residue at position 78 of DBT-1 (C for cysteine, S for serine). Every pair-wise strain comparison is significant except for the N2 DBT-1(S78) - CB4856 comparisons (Tukey’s HSD *p*-value < 1.43E-6).

Because DBT-1 is involved in BCAA catabolism, we hypothesized that the DBT-1(C78S)-dependent difference in progeny length between the N2 and CB4856 strains after arsenic trioxide treatment might be caused by differential larval (L1) arrest through depletion of downstream mmBCFAs. To test this hypothesis, we quantified the abundance of the monomethyl-branched (ISO) and straight-chain (SC) forms of C15 and C17 in the N2, CB4856, and allele-swap genetic backgrounds. We measured the metabolite levels in staged L1 animals and normalized the detected amounts of C15ISO and C17ISO relative to the abundances of C15SC and C17SC, respectively, to mitigate confounding effects of differences in developmental rates that could result from genetic background differences or arsenic trioxide exposure. Generally, the ratios of C15ISO/C15SC and C17ISO/C17SC were reduced in arsenic-treated animals relative to controls (Figure 4B; Figure 4-source data 1–3). However, arsenic trioxide treatment had a 7.6-fold stronger effect on the C15ISO/C15SC ratio in N2, which naturally has the C78 allele, than on the N2 DBT-1(S78) allele swap strain. This difference suggests that the DBT-1(C78) allele is more strongly inhibited by arsenic trioxide (0.04 to 0.004, Tukey HSD *p-*value = 0.0358, n = 6). Similarly, we observed a 6.6-fold arsenic-induced reduction in the C17ISO/C17SC ratio when comparing the N2 DBT-1(C78) and N2 DBT-1(S78) strains (Tukey HSD *p-*value = 0.003747, n = 6). When comparing the CB4856 DBT-1(S78) and CB4856 DBT-1(C78) strains, we observed a 2.8-fold lower C15ISO/C15SC ratio (Tukey HSD *p-*value = 0.0427733, n = 3) and 1.5-fold lower C17ISO/C17SC ratio (Tukey HSD *p-*value = 0.164721, n = 3) in the in the CB4856 DBT-1(C78) strain. We noted that the C17ISO/straight-chain ratio difference was not significantly different between the two CB4856 genetic background strains. However, we observed a significant arsenic-induced decrease in raw C17ISO production in the CB4856 DBT-1(C78) strain (Tukey HSD *p*-value = 0.029) and no significant difference in the CB4856 DBT-1(S78) strain (Tukey HSD *p*-value = 0.1) (Figure 4-figure supplement 1). Importantly, these DBT-1(C78S) allele-specific reductions in ISO/straight-chain ratios were not driven by arsenic-induced differences in straight-chain fatty acids (Figure 4-figure supplement 2). These results explained the majority of the physiological differences between the N2 and CB4856 strains in the presence of arsenic trioxide (Figure 3) and suggested that the DBT-1(C78) allele was inhibited by arsenic trioxide more strongly than DBT-1(S78). Taken together, the differential reduction in branched-chain fatty acids likely underlies the majority of physiological differences between the sensitive and resistant *C. elegans* strains.

In addition to arsenic-induced differences in branched chain fatty acid production, we observed significant differences in branched/straight-chain ratios between the parental and allele swap strains when L1 larval animals were grown in control conditions (Figure 4-figure supplement 3; Figure 4-source data 1-3). Strains with the DBT-1(C78) had higher ISO/SC ratios relative to strains with the DBT-1(S78) for the C17 (CB4856 DBT-1(C78): Tukey HSD *p-*value = 0.0342525, n = 3; N2 DBT-1(C78): Tukey HSD *p-*value = 0.0342525, n = 6) and C15 ratios (CB4856: Tukey HSD *p-*value = 0.0168749, n = 3; N2: Tukey HSD *p-*value = 0.1239674, n = 6) (Figure 4-figure supplement 3; Figure 4-source data 1-3). We noted that the C15ISO/straight-chain ratio was not significantly different when comparing the N2 and N2 allele swap strain, but the direction of effect matched our other observations and we saw significant differences in C15ISO levels (N2-C15ISO DBT-1(C78): Tukey HSD *p-*value = 0.0265059, n = 6, Figure 4-figure supplement 4; Figure 4-source data 1-3). Importantly, the DBT-1 allele-specific differences in the fatty acid ratio and ISO measurements were not driven by differences in straight-chain fatty acids (Figure 4-figure supplement 4). However, we did not observe the same effect of the DBT-1(C78S) allele at the young adult life stage (Figure 4-figure supplement 5; Figure 4-source data 4). Taken together, these results suggest that the DBT-1(C78) allele produces more branched chain fatty acids than the DBT-1(S78) allele, but this effect was dependent on the developmental stage of the animals.

To test the hypothesis that differential arsenic-induced depletion of branched-chain fatty acids in strains with the DBT-1(C78S) causes physiological differences in growth, we tested if mmBCFA supplementation could rescue the effects of arsenic trioxide-induced toxicity. We exposed the parental and the DBT-1 allele-replacement strains to media containing arsenic trioxide alone, C15ISO alone, or a combination of arsenic trioxide and C15ISO. In agreement with previous experiments, the PC1 trait was highly correlated with animal length traits (Supplementary Figure 4-figure supplement 6–7; Figure 4-source data 5-7). When grown in arsenic-trioxide alone the average phenotypic difference between N2 DBT-1(C78) and N2 DBT-1(S78) allele was 3.39 units for the PC1 trait. C15ISO supplementation of the arsenic growth media caused a 56.3% rescue of the allele-specific effect in the N2 genetic background (Figure 4C-figure supplement 8). Similarly, when CB4856 DBT-1(C78) animals were supplemented with C15ISO, the allele-specific PC1 phenotypic difference was reduced by 35.9% when compared to the difference between the CB4856 DBT-1(C78) and CB4856 DBT-1(S78) strains in arsenic trioxide alone (Figure 4C-figure supplement 8). By contrast, CB4856 DBT-1(S78) and N2 DBT-1(S78) phenotypes were unaffected by C15ISO supplementation in arsenic trioxide media. Collectively, these data support the hypothesis that the cysteine/serine variant in DBT-1 contributes to arsenic sensitivity in *C. elegans* by reducing ISO fatty acid biosynthesis, and the DBT-1(C78) variant can be partially rescued by supplementation with mmBCFAs.

### Arsenic exposure increases mmBCFA production and favors a cysteine allele in human DBT1

To test whether our results from *C. elegans* translate to human variation in arsenic sensitivity, we tested the role of DBT1 variation on arsenic trioxide responses and mmBCFA biosynthesis in human cells. The human DBT1 enzyme contains a serine at position 112 that corresponds to the C78 residue in *C. elegans* DBT-1 (Figure 5A-figure supplement 1). Using CRISPR/Cas9, we edited batch cultures of 293 T cells to generate a subset of cells with DBT1(S112C). These cells were exposed to arsenic trioxide or control conditions and we monitored changes in the fraction of cells carrying the DBT1(C112) allele. We found that arsenic exposure caused a 4% increase in the fraction of cells that contained the DBT1(C112) allele (Figure 5B, Fisher’s exact test, *p-* value < 1.9E-16; Figure 5-source data 1-2). Though the human DBT1 does not vary within the human population at S112, two residues in close spatial proximity to S112 do vary among individuals in the population (R113C and W84C) (Figure 5A) [46]. To test the effects of these residues on arsenic trioxide sensitivity, we performed the same editing and arsenic selection procedure described above. Over the course of the selection experiment, cells with DBT1(W84C) and DBT1(R113C) increased by 2% and 1%, respectively (Figure 5B). Therefore, it appears that all three missense variants caused a slight increase in fitness in batch-edited human cell cultures exposed to arsenic. To determine if branched-chain fatty acid production was altered by arsenic exposure, as we found in *C. elegans*, we measured mmBCFA production in unedited 293 T cells in arsenic and mock-treated cultures. We found that overall fatty acid production was markedly reduced in arsenic-treated cultures. In contrast to our observations in *C. elegans*, straight-chain fatty acids were more drastically reduced than ISO fatty acids (Figure 5-source data 3). These results are consistent with our observations that arsenic does not have as strong of an effect on *C. elegans* strains with the DBT-1 S78 allele. Additionally, the DBT1(C112) allele, similar to the *C. elegans* DBT-1(C78) allele, was associated with higher production of mmBCFAs.

**Figure 5:**
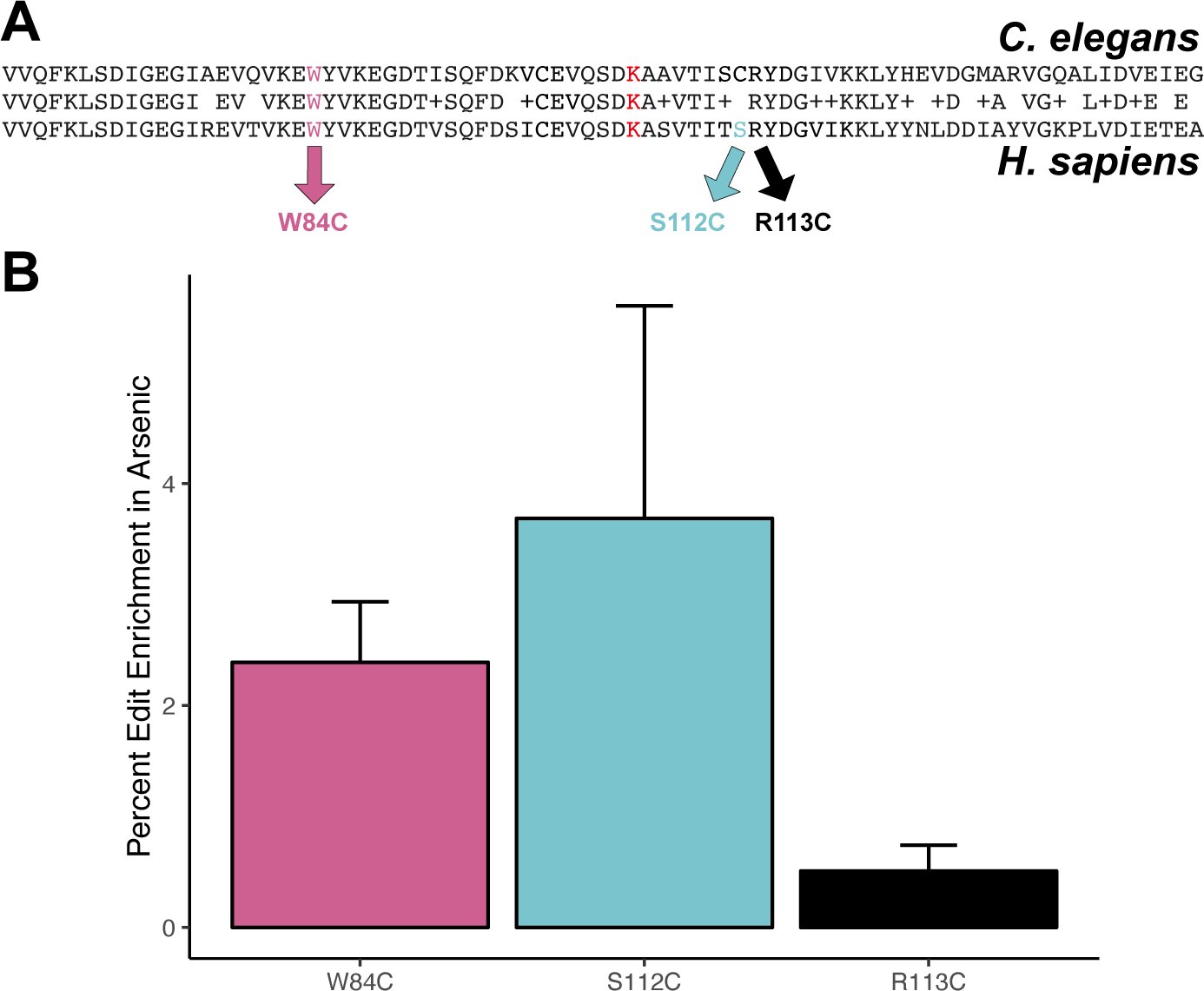
Protective effect of cysteine residues in human DBT1. (A) Alignment of *C. elegans* DBT-1 and *H. sapiens* DBT1. The residues tested for an arsenic-specific effect are indicated with arrows - W84C (pink), S112C (blue), and R113C (black). The lysine that is post-translationally modified with a lipoid acid is highlighted in red. (B) The percent increase of edited human cells that contain the W84C, S112C, or R113C amino acid change in DBT1 in the presence 5 µM arsenic trioxide relative to control conditions are shown. The number of reads in 5 µM arsenic trioxide for all replicates are significantly different from control conditions (Fisher’s exact test, *p-*value < 0.011).

## Discussion

In this study, we characterized the effects of *C. elegans* natural genetic variation on physiological responses to the pervasive environmental toxin arsenic trioxide. Though the effects of this toxin have been extensively studied in a variety of systems [2,3,39,47,48], recent evidence from human population studies have revealed local adaptations within region-specific subpopulations [6,8,9,49]. Our investigation into the natural variation in *C. elegans* responses to arsenic trioxide led to the discovery of a novel mechanism by which this compound could elicit toxicity. We show that arsenic trioxide differentially inhibits two natural alleles of the E2 domain of the BCKDH complex, which is encoded by the *dbt-1* gene. Specifically, strains with the DBT-1(C78) allele are more sensitive to arsenic trioxide than strains carrying the DBT-1(S78). Furthermore, we show that the increased sensitivity of the DBT-1(C78) allele is largely explained by differences in the production of mmBCFAs (Figure 4B-C), which are critical molecules for developmental progression beyond the first larval stage. Arsenic is thought to inhibit the activity of both the pyruvate dehydrogenase (PDH) and the α-ketoglutarate (KGDH) dehydrogenase complexes through interactions with the reduced form of lipoate [39], which is the cofactor for the E2 domain of these complexes. Like the PDH and KGDH complexes, the E2 domain of BCKDH complex requires the cofactor lipoate to perform its enzymatic function [50–52]. The inhibition of DBT-1 by arsenic trioxide could involve three-point coordination of arsenic by the C78 thiol and the reduced thiol groups of the nearby lipoate. However, based on the crystal structure (PDB:1Y8N), the atomic distance between the C78 thiol group and the thiol groups from the lipoylated lysine is ∼32 Å, which might be too large a distance for coordinating arsenic (Figure 5A-figure supplement 1) [53]. Alternatively, arsenic trioxide could inhibit DBT-1(C78) through coordination between the thiol groups of C78 and C65 (∼8.5 Å) (Figure 5A-figure supplement 1). Analogous thiol-dependent mechanisms have been proposed for the inhibition of other enzymes by arsenic [48]. Despite structural similarities and a shared cofactor, no evidence in the literature indicates that BCKDH is inhibited by arsenic trioxide, so these results demonstrate the first connection of arsenic toxicity to BCKDH E2 subunit inhibition.

Multiple sequence alignments show that cysteine residues C65 and C78 of *C. elegans* DBT-1 correspond to residues S112 and C99 of human DBT1 (Figure 5A). Though DBT1 does not vary at position 112 within the human population, two residues (R113C and W84C) in close spatial proximity do [46]. We hypothesized that cysteine variants in DBT1 would sensitize human cells to arsenic trioxide. However, we found that the cysteine variants (W84C, S112C, and R113C) proliferated slightly more rapidly than the parental cells in the presence of arsenic. Notably, a growing body of evidence suggests that certain cancer cells upregulate components involved in BCAA metabolism, and this upregulation promotes tumor growth [54,55]. Perhaps the increased proliferation of human cell lines that contain the DBT1 C112 allele (Figure 5) is caused by increased activity of the BCKDH complex. It is worth noting that human cell lines grown in culture do not have the same strict requirements for mmBCFA, and the requirements for different fatty acids are variable among diverse immortalized cell lines [56,57]. Furthermore, in *C. elegans*, the developmental defects associated with *dbt-1* loss-of-function can be rescued by neuronal-specific expression of *dbt-1* [34], suggesting that the physiological requirements of mmBCFA in *C. elegans* depend on the coordination of multiple tissues that cannot be recapitulated with cell-culture experiments. These results further highlight the complexity of arsenic toxicity, as well as the physiological requirements for BCAA within and across species and could explain the discrepancy between the physiological effects we observed in *C. elegans* and human cell-line experiments. Given that arsenic trioxide has become the standard-of-care for treating APL [58] and is gaining traction in treating other leukemias, it will be important to further explore the effects of arsenic on BCAA metabolism and cancer growth.

The C78 allele of DBT-1 is likely the derived allele in the *C. elegans* population because all other organisms have a serine at the corresponding position. The loss of the serine allele in the *C. elegans* population might have been caused by relaxed selection at this locus, though this hypothesis is difficult to test because of the effects of linked selection and decreased recombination in chromosome centers. It is hypothesized that the *C. elegans* species originated from the Pacific Rim and that the ancestral state more closely resembles the CB5846 strain than the N2 strain [30,59]. The CB4856 strain was isolated from the Hawaiian island of O’ahu [60], where environments could have elevated levels of arsenic in the soil from volcanic activity, the farming of cane sugar, former construction material (canec) production facilities, or wood treatment plants (Hawaii.gov). It is possible that as the *C. elegans* species spread across the globe into areas with lower levels of arsenic in the soil and water, the selective pressure to maintain high arsenic tolerance was relaxed and the cysteine allele appeared. Alternatively, higher mmBCFA levels at the L1 larval stage in strains with the DBT-1(C78) allele (Figure 4-figure supplement 3-4) might cause faster development in certain conditions, although we did not observe allele-specific growth differences in laboratory conditions. Despite these clues suggesting selection in local environments, the genomic region surrounding the *dbt-1* locus does not show a signature of selection as measured by Tajima’s D [61] (Figure 2-figure supplement 5; Figure 2-source data 9), and the strains with the DBT-1 S78 allele show no signs of geographic clustering (Figure 2-figure supplement 6; Figure 2-source data 10). Our study suggests that *C. elegans* is a powerful model to investigate the molecular mechanisms for how populations respond to environmental toxins.

## Materials and methods

### Key Resources Table

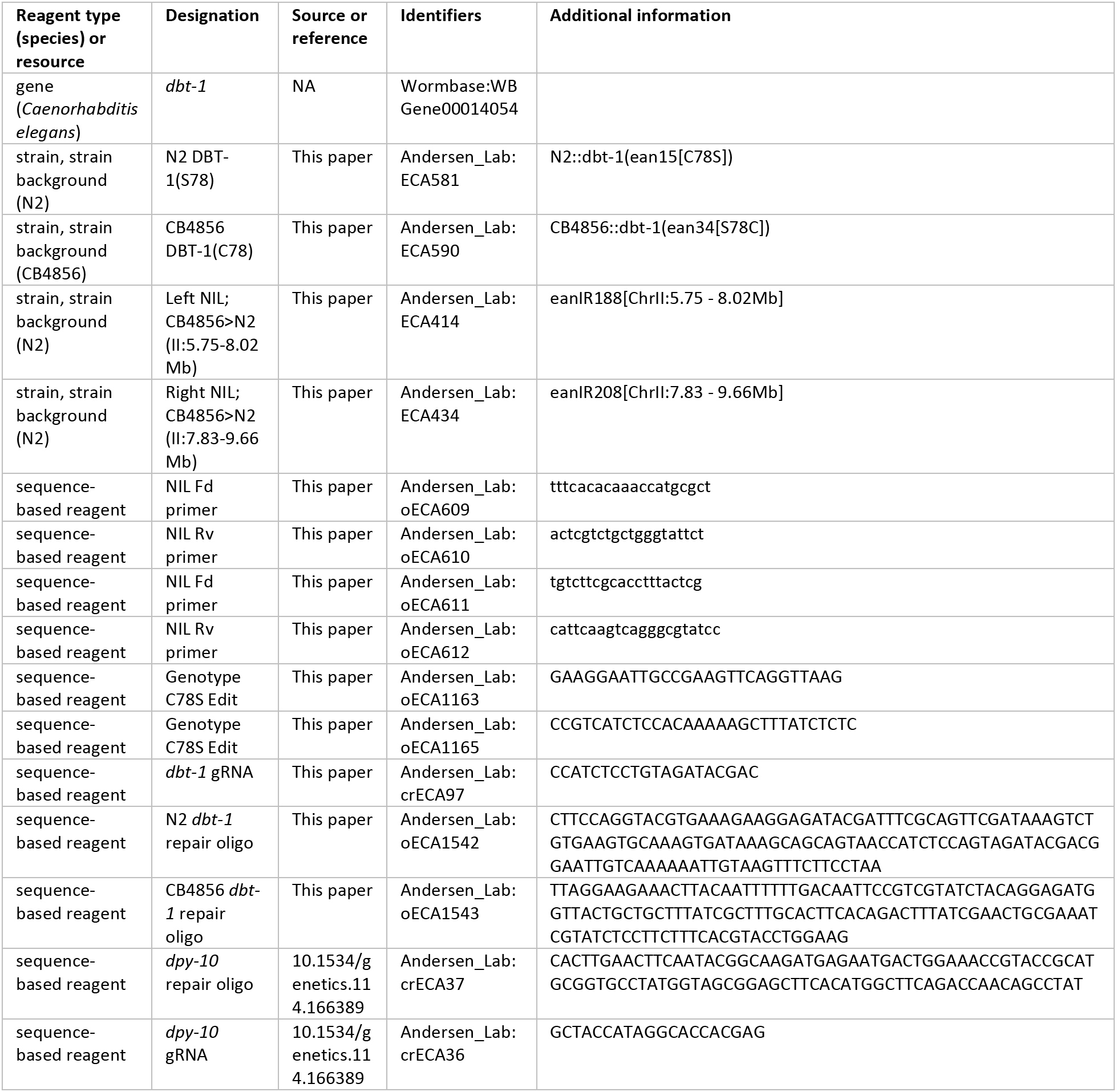
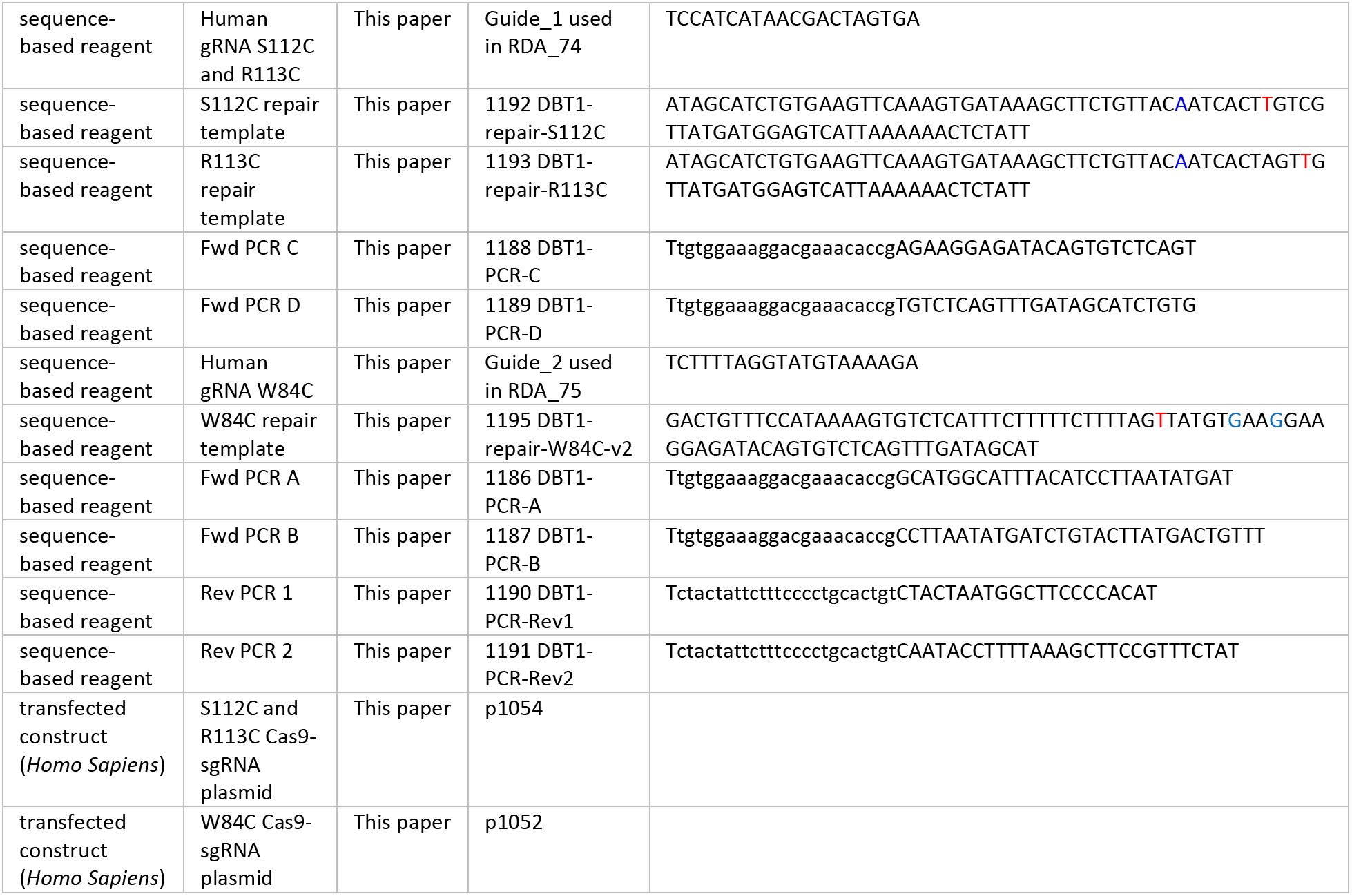

### Strains

Animals were fed the bacterial strain OP50 and grown at 20°C on modified nematode growth medium (NGM), containing 1% agar and 0.7% agarose to prevent burrowing of the wild isolates [62]. For each assay, strains were grown for five generations with no strain entering starvation or encountering dauer-inducing conditions [63]. Wild *C. elegans* isolates used for genome-wide association and recombinant inbred advanced intercross lines (RIAILs) used for linkage mapping have been described previously [31,32,35]. Strains constructed for this manuscript are listed above in the Key Resources Table.

### High-throughput fitness assay

We used the high-throughput fitness assay (HTA) described previously [35]. In short, strains are passaged for four generations before bleach-synchronization and aliquoted to 96-well microtiter plates at approximately one embryo per microliter in K medium [62]. The final concentration of NaCl in the K medium for the genome-wide association (GWA) and linkage mapping assays was 51 mM. For all subsequent experiments the final NaCl concentration was 10.2 mM. The following day, hatched and synchronized L1 animals were fed HB101 bacterial lysate (Pennsylvania State University Shared Fermentation Facility, State College, PA, [64]) at a final concentration of 5 mg/ml and grown for two days at 20°C. Next, three L4 larvae were sorted using a large-particle flow cytometer (COPAS BIOSORT, Union Biometrica, Holliston, MA) into microtiter plates that contain HB101 lysate at 10 mg/ml, K medium, 31.25 µM kanamycin, and either arsenic trioxide dissolved in 1% water or 1% water alone. The animals were then grown for four days at 20°C. The resulting populations were treated with sodium azide (50 mM) prior to being measured with the BIOSORT. All raw experimental data can be found on FigShare (https://figshare.com/articles/Raw_experimental_files_used_for_manuscript/7458980).

### Calculation of fitness traits for genetic mappings

Phenotype data generated using the BIOSORT were processed using the R package *easysorter*, which was specifically developed for processing this type of data [65]. Briefly, the function *read_data*, reads in raw phenotype data and runs a support vector machine to identify and eliminate bubbles. Next, the *remove_contamination* function eliminates any wells that were identified as contaminated prior to scoring population parameters. This analysis results in processed BIOSORT data where each observation is for a given strain corresponds to the measurements for an individual animal. However, the phenotypes we used for mapping and follow-up experiments are summarized statistics of populations of animals in each well of a 96-well plate. The *sumplate* function was used to generate summary statistics of the measured parameters for each animal in each well. These summary statistics include the 10th, 25th, 50th, 75th, and 90th quantiles for time of flight (TOF), animal extinction (EXT), and three fluorescence channels (Green, Yellow, and Red), which correspond to animal length, optical density, and ability to pump fluorescent beads, respectively. Measured brood sizes (n) are normalized by the number of animals that were originally sorted into each well (norm.n). For mapping experiments, a single well replicate for each strain is summarized using the *sumplate* function. For follow-up experiments, multiple replicates for each strain indicated by a unique plate, well, and column were used. After summary statistics for each well are calculated, we accounted for differences between assays using the *regress(assay = TRUE)* function in the *easysorter* package. Outliers in the GWA and linkage mapping experiments were identified and eliminated using the *bamf_prune* function in *easysorter*. For follow-up experiments that contained multiple replicates for each strain, we eliminated strain replicates that were more than two standard deviations from the strain mean for each condition tested. Finally, arsenic-specific effects were calculated using the *regress(assay = FALSE)* function from *easysorter*, which accounts for strain-specific differences in growth parameters present in control conditions.

### Principal component analysis of processed BIOSORT measured traits

The COPAS BIOSORT measures individual animal length (TOF), optical density (EXT), fluorescence (green, yellow, and red). We use these data to calculate the total number of animals in a well and then normalize by the number of animals initially sorted into the well (brood size). All of these measurements were then summarized using the *easysorter* package to generate various summary statistics of each measured parameter, including five distribution quantiles and measures of dispersion [35]. Because fluorescent beads were only used for linkage and GWA mapping experiments, we eliminated all of these traits prior to PCA analysis for all non-mapping experiments. Additionally, we removed all summary statistic traits related to data dispersion (IQR, variance, and coefficient of variation) for all experiments. Prior to principal component analysis (PCA), HTA phenotypes were scaled to have a mean of zero and a standard deviation of one using the *scale* function in R. PCA was performed using the *prcomp* function in R [66]. Eigenvectors were subsequently extracted from the object returned by the *prcomp* function.

### Arsenic trioxide dose-response assays

All dose-response experiments were performed on four genetically diverged strains (N2, CB4856, DL238, and JU775) in technical quadruplicates prior to performing GWA and linkage mapping experiments (Figure 1-source data 1). Animals were assayed using the HTA, and phenotypic analyses were performed as described above. The arsenic trioxide concentration for GWA and linkage mapping experiments was chosen based on an observable effect for animal length and brood size phenotypes in the presence of arsenic.

### Heritability calculations

For dose response experiments, broad-sense heritability (*H*^2^) estimates were calculated using the *lmer* function in the lme4 package with the following linear mixed model (phenotype ∼1 + (1|strain)) [67]. *H*^2^ was then calculated as the fraction of the total variance that can be explained by the random component (strain) of the mixed model. Prior to estimating *H*^2^, we removed outlier replicates that we defined as replicates with values greater than two standard deviations away from the mean phenotype. Outliers were defined on a per-trait and per-strain basis. Heritability estimates for dose response experiments are in Figure 1-source data 2.

Heritability estimates for the linkage mapping experiment were calculated using two approaches. In both approaches, we used the previously described RIAIL genotype matrix to compute relatedness matrices [35]. In the first approach, a variance component model using the R package regress was used to estimate the fraction of phenotypic variation explained by additive and epistatic genetic factors, *H*^2^, or just additive genetic factors, *h*^2^ [68,69], using the formula (y ∼ 1, ∼ZA + ZAA), where y is a vector of RIAIL phenotypes, ZA is the additive relatedness matrix, and ZAA is the pairwise-interaction relatedness matrix. The additive relatedness matrix was calculated as the correlation of marker genotypes between each pair of strains. In addition, a two-component variance model was calculated with both an additive and pairwise-interaction effect. The pairwise-interaction relatedness matrix was calculated as the Hadamard product of the additive relatedness matrix.

The second approach utilized a linear mixed model and the realized additive and epistatic relatedness matrices [70–73]. We used the *mmer* function in the sommer package with the formula (y ∼ A + E) to estimate variance components, where y is a vector of RIAIL phenotypes, A is the realized additive relatedness matrix, and E is the epistatic relatedness matrix. This same approach was used to estimate heritability for the GWA mapping phenotype data, with the only difference being that we used the wild isolate genotype matrix described below. Heritability estimates for RIAIL and wild isolate data are in Figure 1-source data 12 and Figure 2-source data 6, respectively.

### Effect size calculations for dose response assay

We first fit a linear model with the formula (phenotype ∼ strain) for all measured and principal component traits for each concentration of arsenic trioxide using the *lm* R function. Next, we extracted effect sizes using the *anova_stats* function from the sjstats R package [74]. Effect sizes for dose responses are in Figure 1-source data 3.

### Linkage mapping

A total of 262 RIAILs were phenotyped in the HTA described previously for control and arsenic trioxide conditions [35,36]. The phenotype and genotype data were entered into R and scaled to have a mean of zero and a variance of one for linkage analysis (Figure 1-source data 8). Quantitative trait loci (QTL) were detected by calculating logarithm of odds (LOD) scores for each marker and each trait as −*n*(*ln*(1 − *r*^2^)/2*ln*(10)), where *r* is the Pearson correlation coefficient between RIAIL genotypes at the marker and phenotype trait values [75]. The maximum LOD score for each chromosome for each trait was retained from three iterations of linkage mappings (Figure 1-source data 11). We randomly permuted the phenotype values of each RIAIL while maintaining correlation structure among phenotypes 1000 times to estimate the significance threshold empirically. The significance threshold was set using a genome-wide error rate of 5%. Confidence intervals were defined as the regions contained within a 1.5 LOD drop from the maximum LOD score [76].

### Near-isogenic line (NIL) generation

NILs were generated by crossing N2xCB4856 RIAILs to each parental genotype. For each NIL, eight crosses were performed followed by six generations of selfing to homozygose the genome. Reagents used to generate NILs are detailed in the Key Resources Table. The NILs responses to 1000 µM arsenic trioxide were quantified using the HTA described above (Figure 1-source data 13). NIL whole-genome sequencing and analysis was performed as described previously [77] (Figure 1-source data 16).

### Genome-wide association mapping

Genome-wide association (GWA) mapping was performed using phenotype data from 86 *C. elegans* isotypes (Figure 2-source data 1). Genotype data were acquired from the latest VCF release (Release 20180527) from CeNDR that was imputed as described previously [32]. We used BCFtools [78] to filter variants that had any missing genotype calls and variants that were below 5% minor allele frequency. We used PLINK v1.9 [79,80] to LD-prune the genotypes at a threshold of *r*^2^ < 0.8, using *–indep-pairwise 50 10 0.8.* This resulting genotype set consisted of 59,241 markers that were used to generate the realized additive kinship matrix using the *A.mat* function in the *rrBLUP* R package [70] (Figure 2-source data 5). These markers were also used for genome-wide mapping. However, because these markers still have substantial LD within this genotype set, we performed eigen decomposition of the correlation matrix of the genotype matrix using *eigs_sym* function in Rspectra package [81]. The correlation matrix was generated using the *cor* function in the correlateR R package [82]. We set any eigenvalue greater than one from this analysis to one and summed all of the resulting eigenvalues [83]. This number was 500.761, which corresponds to the number of independent tests within the genotype matrix. We used the *GWAS* function in the rrBLUP package to perform genome-wide mapping with the following command: *rrBLUP::GWAS(pheno = PC1, geno = Pruned_Markers, K = KINSHIP, min.MAF = 0.05, n.core = 1, P3D = FALSE, plot = FALSE)*. To perform fine-mapping, we defined confidence intervals from the genome-wide mapping as +/-100 SNVs from the rightmost and leftmost markers above the Bonferroni significance threshold. We then generated a QTL region of interest genotype matrix that was filtered as described above, with the one exception that we did not perform LD pruning. We used PLINK v1.9 to extract the LD between the markers used for fine mapping and the peak QTL marker identified from the genome-wide scan. We used the same command as above to perform fine mapping, but with the reduced variant set. The workflow for performing GWA mapping can be found on https://github.com/AndersenLab/cegwas2-nf. All trait mapping results can be found on FigShare (https://figshare.com/articles/GWA_results_for_all_traits_mapped_in_manuscript/7458932).

### Generation of *dbt-1* allele replacement strains

Allele replacement strains were generated using CRISPR/Cas9-mediated genome editing, using the co-CRISPR approach [84] with Cas9 ribonucleoprotein delivery [85]. Alt-RTM crRNA and tracrRNA were purchased from IDT (Skokie, IL). tracrRNA (IDT, 1072532) was injected at a concentration of 13.6 µM. The *dpy-10* and the *dbt-1* crRNAs were injected at 4 µM and 9.6 µM, respectively. The *dpy-10* and the *dbt-1* single-stranded oligodeoxynucleotides (ssODN) repair templates were injected at 1.34 µM and 4 µM, respectively. Cas9 protein (IDT, 1074182) was injected at 23 µM. To generate injection mixes, the tracrRNA and crRNAs were incubated at 95°C for five minutes and 10°C for 10 minutes. Next, Cas9 protein was added and incubated for five minutes at room temperature. Finally, repair templates and nuclease-free water were added to the mixtures and loaded into pulled injection needles (1B100F-4, World Precision Instruments, Sarasota, FL). Individual injected *P_0_* animals were transferred to new 6 cm NGM plates approximately 18 hours after injections. Individual *F_1_* rollers were then transferred to new 6 cm plates to generate self-progeny. The region surrounding the desired S78C (or C78S) edit was then amplified from *F_1_* rollers using primers oECA1163 and oECA1165. The PCR products were digested using the *Sfc*I restriction enzyme (R0561S, New England Biolabs, Ipswich, MA). Differential band patterns signified successfully edited strains because the N2 S78C, which is encoded by the CAG codon, creates an additional *Sfc*I cut site. Non-Dpy, non-Rol progeny from homozygous edited *F_1_* animals were propagated. If no homozygous edits were obtained, heterozygous *F_1_* progeny were propagated and screened for the presence of the homozygous edits. *F_1_* and *F_2_* progeny were then Sanger sequenced to verify the presence of the proper edited sequence. The phenotypes of allele swap strains in control and arsenic trioxide conditions were measured using the HTA described above (Figure 1-source data 13). PCA phenotypes for allele-swap strains were generated the same way as described above for GWA mapping traits and are located in Figure 1-source data 13.

### Rescue with 13-methyltetradecanoic acid

Strains were grown as described for a standard HTA experiment. In addition to adding arsenic trioixde to experimental wells, we also added a range of C15iso (13-methyltetradecanoic acid, Matreya Catalog # 1605) concentrations to assay rescue of arsenic effects (Figure 4-source data 5).

### Growth conditions for metabolite profiling

For L1 larval stage assays, chunks (∼1 cm) were taken from starved plates and placed on multiple fresh 10 cm plates. Prior to starvation, animals were washed off of the plates using M9, and embryos were prepared by bleach synchronization. Approximately 40,000 embryos were resuspended in 25 ml of K medium and allowed to hatch overnight at 20°C. L1 larvae were fed 15 mg/ml of HB101 lysate the following morning and allowed to grow at 20°C for 72 hours. We harvested 100,000 embryos from gravid adults by bleaching. These embryos were hatched overnight in 50 ml of K medium in a 125 ml flask. The following day, we added arsenic trioxide to a final concentration of 100 µM and incubated the cultures for 24 hours. After 24 hours, we added HB101 bacterial lysate (2 mg/ml) to each culture. Finally, we transferred the cultures to 50 ml conical tubes, centrifuged the cultures at 3000 RPM for three minutes to separate the pellet and supernatant. The supernatant and pellets from the cultures were frozen at −80°C and prepared for analysis. For young adult stage assays, 45,000 animals per culture were prepared as described above but in S medium, at a density of three animals per microliter, and fed HB101 lysate (5 mg/mL). These cultures were shaken at 200 RPM, 20°C in 50 mL Erlenmeyer flasks for 62 h. For harvesting, we settled 15 mL of cultures for 15 minutes at room temperature and then pipetted the top 12 mL of solution off of the culture. The remaining 3 mL of animal pellet was washed with 10 mL of M9, centrifuged at 1000 g for one minute, and then the supernatant removed. This wash was repeated once more with M9 and again with water. The final nematode pellet was snap frozen in liquid nitrogen.

### Nematode extraction

Pellets were lyophilized 18-24 hours using a VirTis BenchTop 4 K Freeze Dryer until a chalky consistency was achieved. Dried pellets were transferred to 1.5 mL microfuge tubes and dry pellet weight recorded. Pellets were disrupted in a Spex 1600 MiniG tissue grinder after the addition of three stainless steel grinding balls to each sample. Microfuge tubes were placed in a Cryoblock (Model 1660) cooled in liquid nitrogen, and samples were disrupted at 1100 RPM for two cycles of 30 seconds. Each sample was individually dragged across a microfuge tube rack eight times, inverted, and flicked five times to prevent clumping. This process was repeated two additional rounds for a total of six disruptions. Pellets were transferred to 4 mL glass vials in 3 mL 100% ethanol. Samples were sonicated for 20 minutes (on/off pulse cycles of two seconds at power 90 A) using a Qsonica Ultrasonic Processor (Model Q700) with a water bath cup horn adaptor (Model 431C2). Following sonication, glass vials were centrifuged at 2750 RCF for five minutes in an Eppendorf 5702 Centrifuge using rotor F-35-30-17. The resulting supernatant was transferred to a clean 4 mL glass vial and concentrated to dryness in an SC250EXP Speedvac Concentrator coupled to an RVT5105 Refrigerated Vapor Trap (Thermo Scientific). The resulting powder was suspended in 100% ethanol according to its original dry pellet weight: 0.01 mL 100% ethanol per mg of material. The suspension was sonicated for 10 minutes (pulse cycles of two seconds on and three seconds off at power 90 A) followed by centrifugation at 20,817 RCF in a refrigerated Eppendorf centrifuge 5417R at 4°C. The resulting supernatant was transferred to an HPLC vial containing a Phenomenex insert (cat #AR0-4521-12) and centrifuged at 2750 RCF for five minutes in an Eppendorf 5702 centrifuge. The resulting supernatant was transferred to a clean HPLC vial insert and stored at −20°C or analyzed immediately.

### Mass spectrometric analysis

Reversed-phase chromatography was performed using a Dionex Ultimate 3000 Series LC system (HPG-3400 RS High Pressure pump, TCC-3000RS column compartment, WPS-3000TRS autosampler, DAD-3000 Diode Array Detector) controlled by Chromeleon Software (ThermoFisher Scientific) and coupled to an Orbitrap Q-Exactive mass spectrometer controlled by Xcalibur software (ThermoFisher Scientific). Metabolites were separated on a Kinetex EVO C18 column, 150 mm x 2.1 mm, particle size 1.7 µm, maintained at 40°C with a flow rate of 0.5 mL/min. Solvent A: 0.1% ammonium acetate in water; solvent B: acetonitrile (ACN). A/B gradient started at 5% B for 30 seconds, followed by a linear gradient to 95% B over 13.5 minutes, then a linear gradient to 100% B over three minutes. 100% B was maintained for one minute. Column was washed after each run with 5:1 isopropanol:ACN, flow rate of 0.12 mL/min for five minutes, followed by 100% ACN for 2.9 minutes, a linear gradient to 95:5 water:ACN over 0.1 minutes, and then 95:5 water:ACN for two minutes with a flow rate of 0.5 mL/min. A heated electrospray ionization source (HESI-II) was used for the ionization with the following mass spectrometer parameters: spray voltage: 3 kV; capillary temperature: 320°C; probe heater temperature: 300°C; sheath gas: 70 AU; auxiliary gas flow: 2 AU; resolution: 240,000 FWHM at m/z 200; AGC target: 5e6; maximum injection time: 300 ms. Each sample was analyzed in negative and positive modes with m/z range 200-800. Fatty acids and most ascarosides were detected as [M-H]-ions in negative ionization mode. Peaks of known abundant ascarosides and fatty acids were used to monitor mass accuracy, chromatographic peak shape, and instrument sensitivity for each sample. Processed metabolite measures can be found in Figure 4-source data 1–4 [86].

### Statistical analyses

All *p*-values testing the differences of strain phenotypes in the NIL, allele-replacement, and C15ISO experiments were performed in R using the *TukeyHSD* function with an ANOVA model with the formula (*phenotype ∼ strain*). *p*-values of individual pairwise strain comparisons are reported in each figure legend.

### CRISPR-Cas9 gene editing in human cells

Gene-editing experiments were performed in a single parallel culture experiment using human 293 T cells (ATCC) grown in DMEM with 10% FBS. On day zero, 300,000 cells were seeded per well in a six-well plate format. The following day, two master mixes were prepared: a) LT-1 transfection reagent (Mirus) was diluted 1:10 in Opti-MEM and incubated for five minutes; b) a DNA mix of 500 ng Cas9-sgRNA plasmid (Supplementary File 1-2) with 250 pmol repair template oligonucleotide was diluted in Opti-MEM in a final volume of 100 μL. 250 μL of the lipid mix was added to each of the DNA mixes and incubated at room temperature for 25 minutes. Following incubation, the full 350 μL volume of DNA and lipid mix was added dropwise to the cells. These six-well plates were then centrifuged at 1000 x g for 30 minutes. After six hours, the media on the cells was replaced. For the next six days, cells were expanded and passaged as needed. On day seven, one million cells were taken from each set of edited and unedited cells and placed into separate T75s with either media-only or 5 µM arsenic-containing media. Days seven to fourteen, arsenic and media-only conditions were maintained at healthy cell densities. Days fourteen to eighteen, arsenic exposed cell populations were maintained off arsenic to allow the populations to recover prior to sequencing. Media-only conditions were maintained in parallel. On day eighteen, all arsenic and media-only conditions were pelleted for genomic DNA extraction.

### Analysis of CRISPR-Cas9 editing in human cells

Genomic DNA was extracted from cell pellets using the QIAGEN (QIAGEN, Hilden, Germany) Midi or Mini Kits based on the size of the cell pellet (51183, 51104) according to the manufacturer’s recommendations. DBT1 loci were first amplified with 17 cycles of PCR using a touchdown protocol and the NEBnext 2x master mix (New England Biolabs M0541). The resulting product served as input to a second PCR, using primers that appended a sample-specific barcode and the necessary adaptors for Illumina sequencing. The resulting DNA was pooled, purified with SPRI beads (A63880, Beckman Coulter, Brea, CA), and sequenced on an Illumina MiSeq with a 300-nucleotide single-end read with an eight nucleotide index read. For each sample, the number of reads exactly matching the wild-type and edited DBT1 sequence were determined (Figure 5-source data 1).

### Preparing human cells for Mass Spectroscopy

Mass spectroscopy experiments used human 293 T cells (ATCC) grown in DMEM with 10% FBS. On day zero, 150,000 cells were seeded into 15 cm tissue cultures dishes with 15 mL of either 2.5 µM arsenic or no arsenic media. Each condition had five replicates. On day three, the no arsenic cells were approaching confluence and required passaging. Arsenic conditions were at ∼30% confluence and received a media change. On day seven, both conditions were near confluence, media was removed, and plates were rinsed with ice cold PBS, remaining liquid removed. Plates were frozen at −80°C before processing for mass spectrometric analysis. Cells were scraped off the plates with PBS and pelleted in microfuge tubes. Cell pellets were lyophilized 18-24 hours using a VirTis BenchTop 4 K Freeze Dryer and extracted in 100% ethanol using the same sonication program as described for nematode extraction. Following sonication, samples were centrifuged at 20,817 RCF in a refrigerated Eppendorf centrifuge 5417R at 4 °C. Clarified supernatant was aliquoted to a new tube and concentrated to dryness in an SC250EXP Speedvac Concentrator coupled to an RVT5105 Refrigerated Vapor Trap (Thermo Scientific). The resulting material was suspended in 0.01 mL 100% ethanol and analyzed by LC-MS as described. Metabolite measurements can be found in Figure 5-source data 3.

### Tajima’s D calculation

We used the VCF corresponding to CeNDR release 20160408 (https://elegansvariation.org/data/release/20160408) to calculate Tajima’s D. Tajima’s D was calculated using the *tajimas_d* function in the *cegwas* package using default parameters (window size = 500 SNVs, sliding window distance = 50 SNVs, outgroup = N2) (Figure 2-source data 9). Isolation locations of strains can be found in Figure 2-source data 10.

## Acknowledgments

The authors would like to thank Samuel Rosenberg for assistance on early mappings of drug sensitivities, Mudra Hegde of the Broad Institute for assistance with sequence analysis, and members of the Andersen laboratory for critical reading of this manuscript.

## Supplementary Figures

**Figure 1-figure supplement 1:**
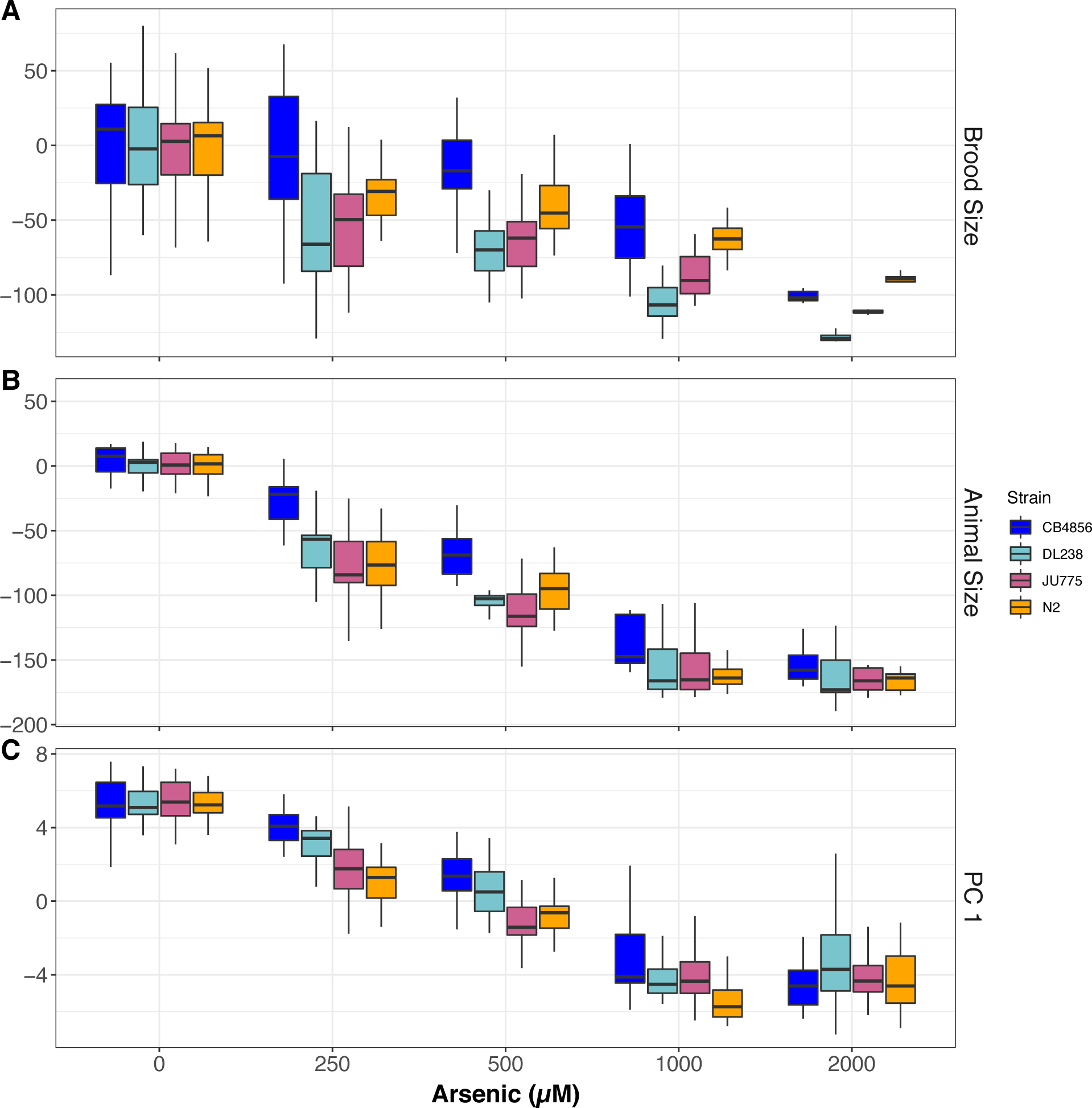
Arsenic trioxide dose response of four diverged *C. elegans* strains. Arsenic trioxide concentration in µM is plotted on the x-axis and the (A) normalized brood size, (B) median progeny length, or (C) the first principal component are plotted on the y-axis. For panels A-B, the y-axis values represent individual phenotypic measurements subtracted from the mean value in 0 µM arsenic trioxide. At least 15 replicates for each strain and condition are represented by Tukey box plots. Box plots are colored by strain (CB4856:blue, DL238:teal, JU775:pink, and N2:orange).

**Figure 1-figure supplement 2:**
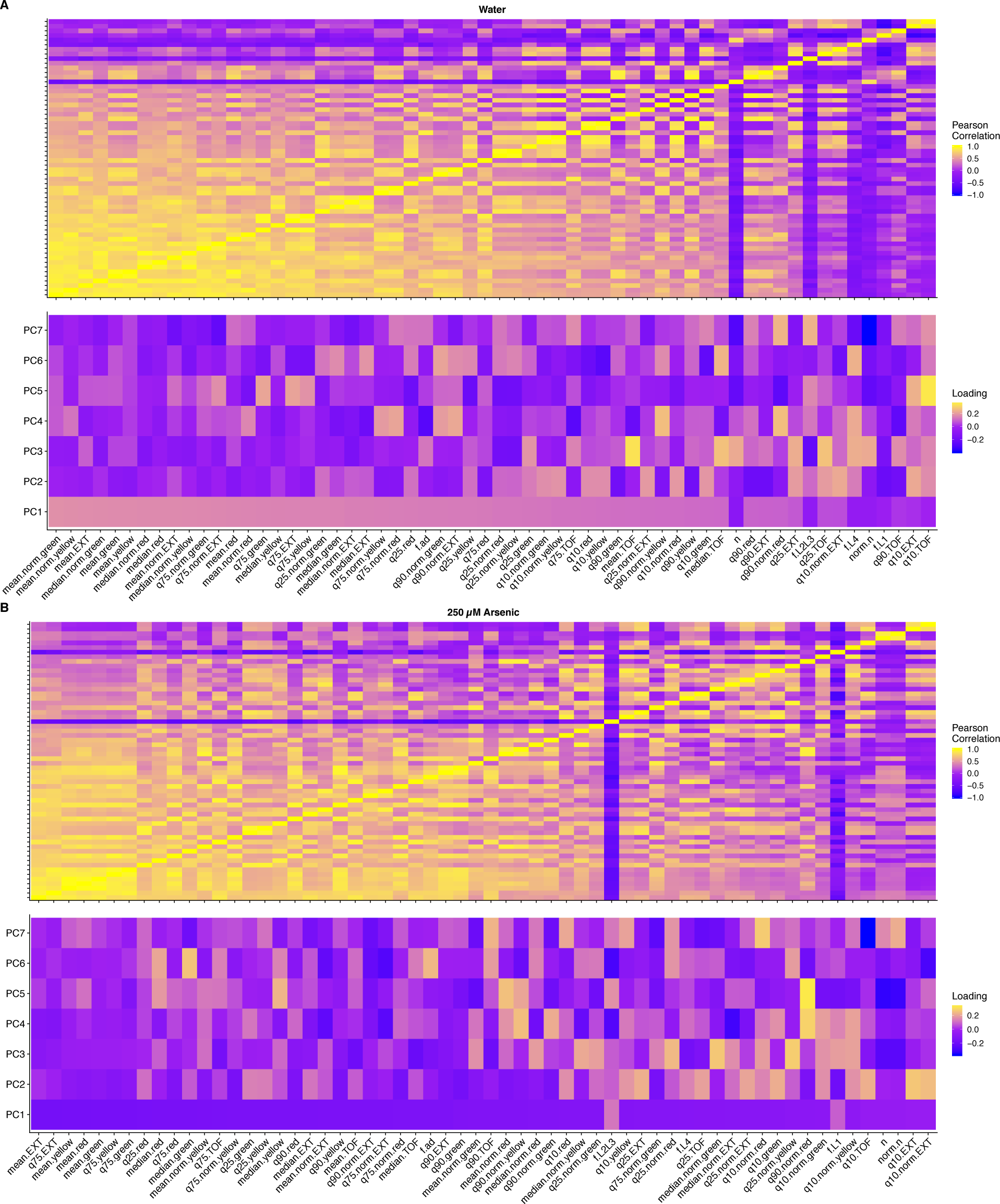
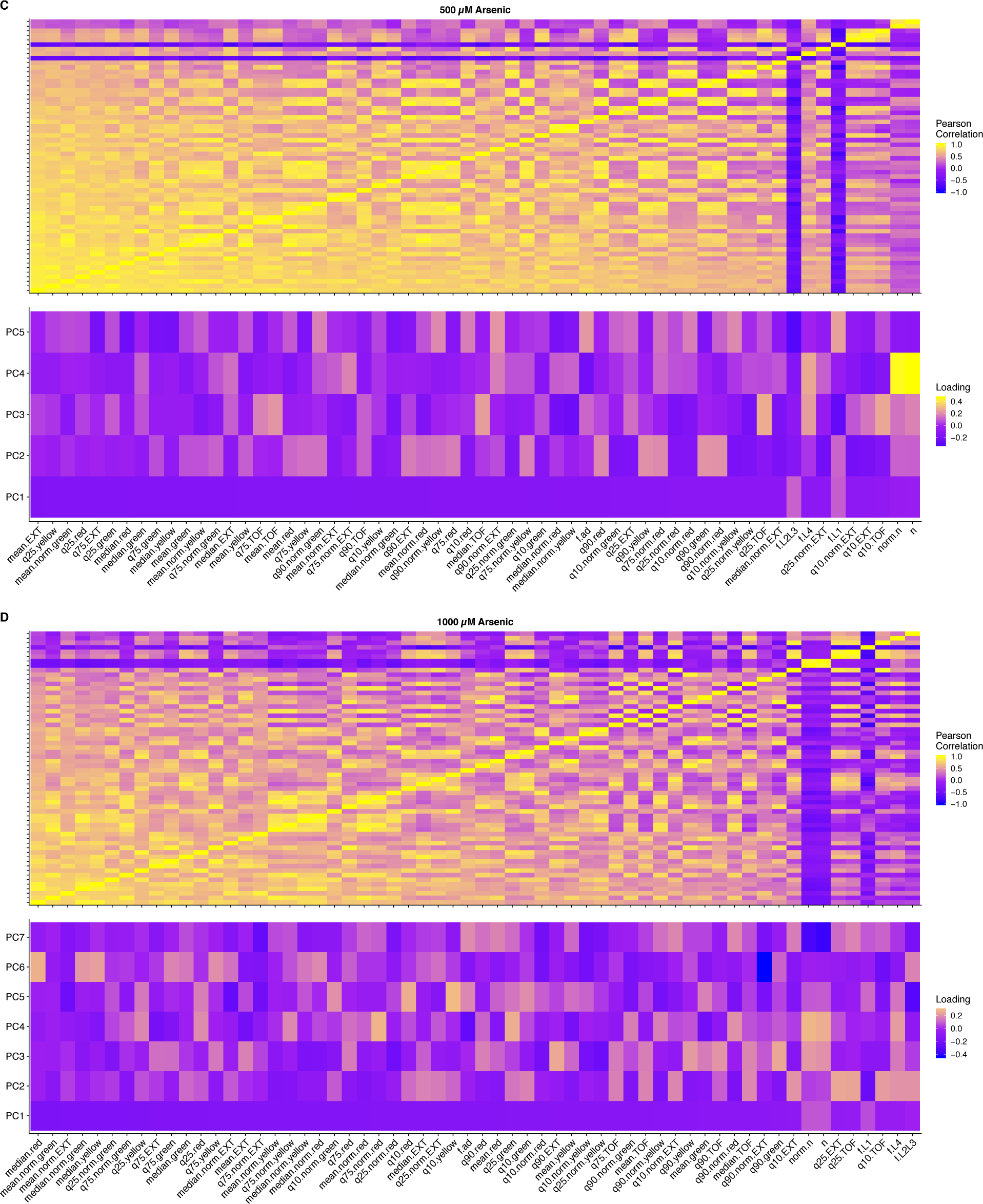
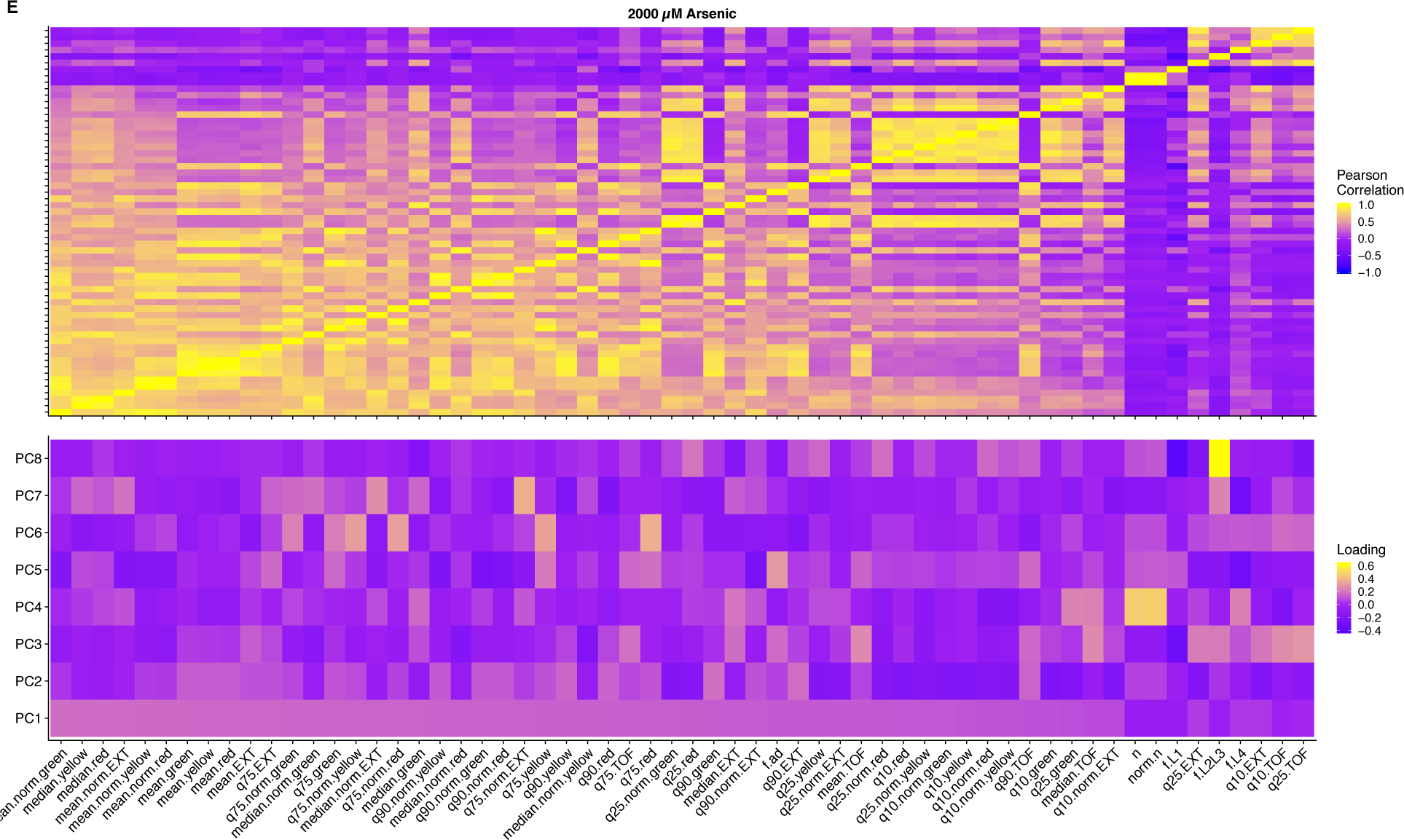
Trait correlations and principal component loadings of arsenic trioxide dose response. The top panels for A-E represent the trait correlations of the measured traits. The bottom panels for A-E represent the contribution of each trait to the principal components that explain 90% of the total variance in the experiment for each condition: (A) Water, (B) 250 µM, (C) 500 µM, (D) 1000 µM, and (E) 2000µM of arsenic trioxide. For each plot, the tile color corresponds to the value corresponding to trait Pearson’s correlation coefficient for the top panels and principal component loading value for the bottom panels, where yellow colors represent higher values.

**Figure 1-figure supplement 3:**
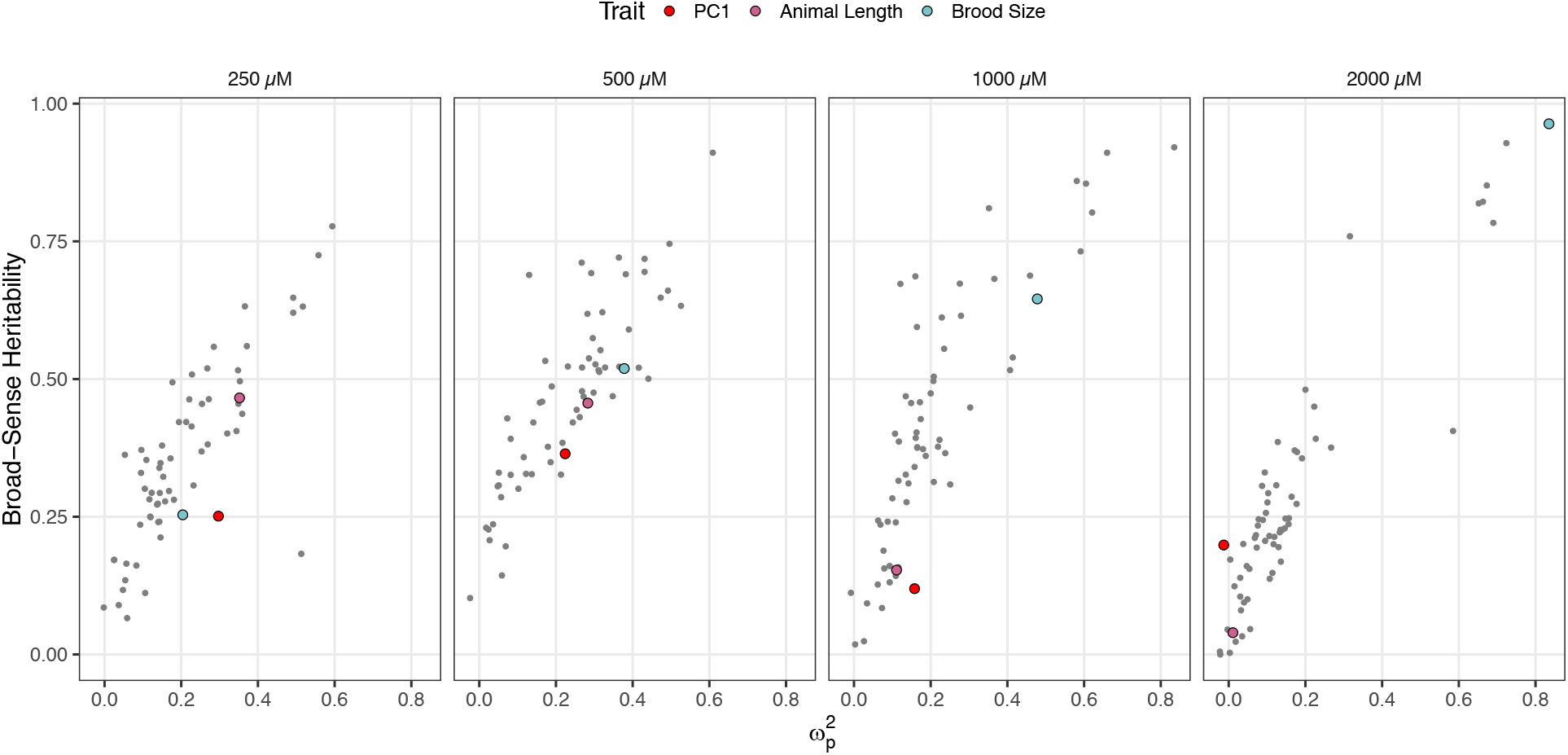
Effect size and broad-sense heritability estimates for the arsenic trioxide dose response. Each panel corresponds to a concentration of arsenic trioxide concentration, which is indicated above the plot panel. The x-axis represents the partial omega squared (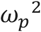) effect-size estimate and the y-axis represent the broad-sense heritability estimate (*H*^2^). Each dot represents a different measured or principal component trait. The three traits discussed throughout the manuscript are highlighted — red: first principal component, pink: animal length, and blue: brood size.

**Figure 1-figure supplement 4:**
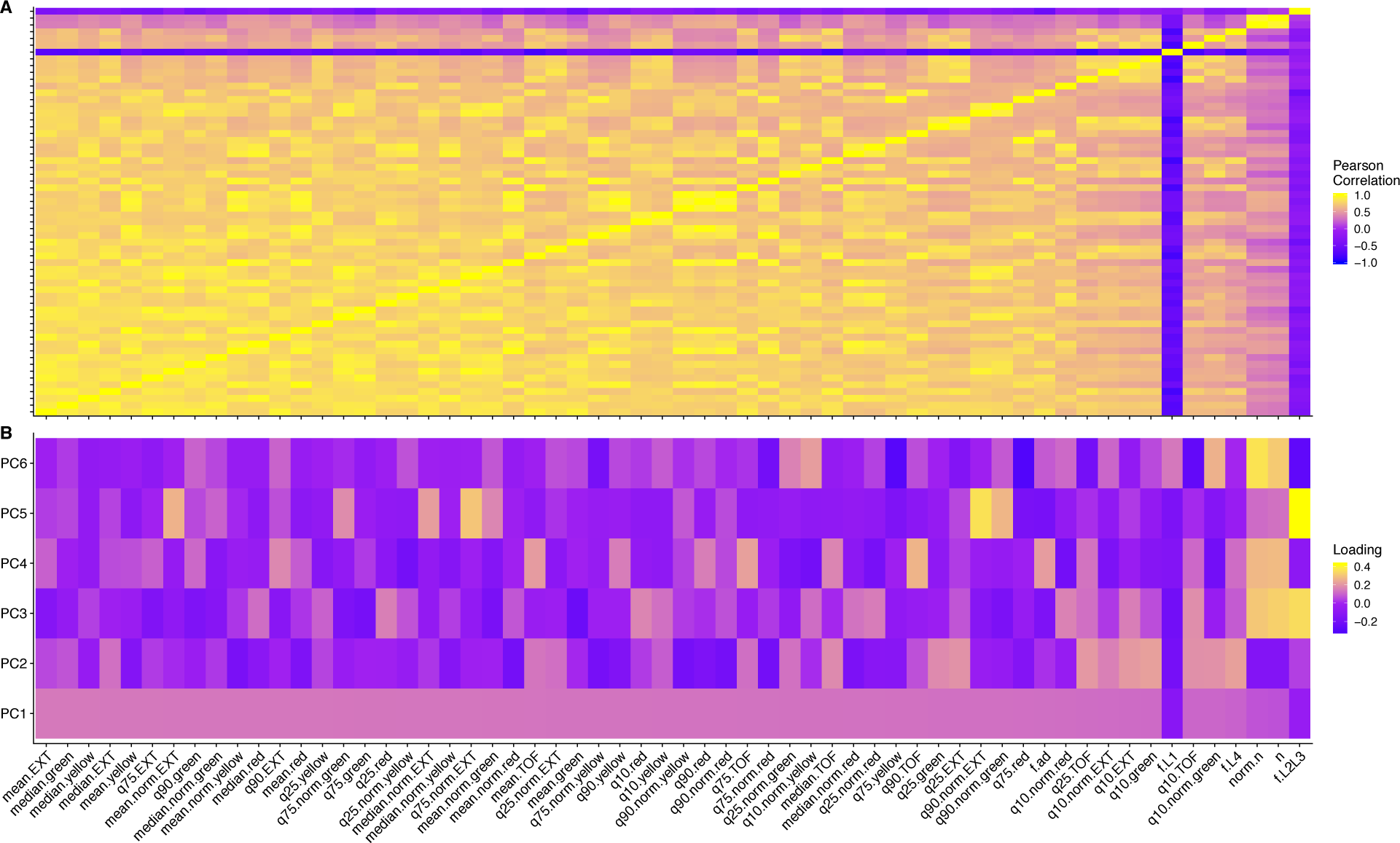
Trait correlations and principal component loadings of linkage mapping experiment. (A) The trait Pearson’s correlation coefficient of the assay- and control-regressed measured traits. (B) The contribution of each measured trait to the principal components that explain 90% of the total variance in the linkage mapping experiment, which was performed at 1000 µM. For each plot, the tile color corresponds to the value, where yellow colors represent higher values.

**Figure 1-figure supplement 5:**
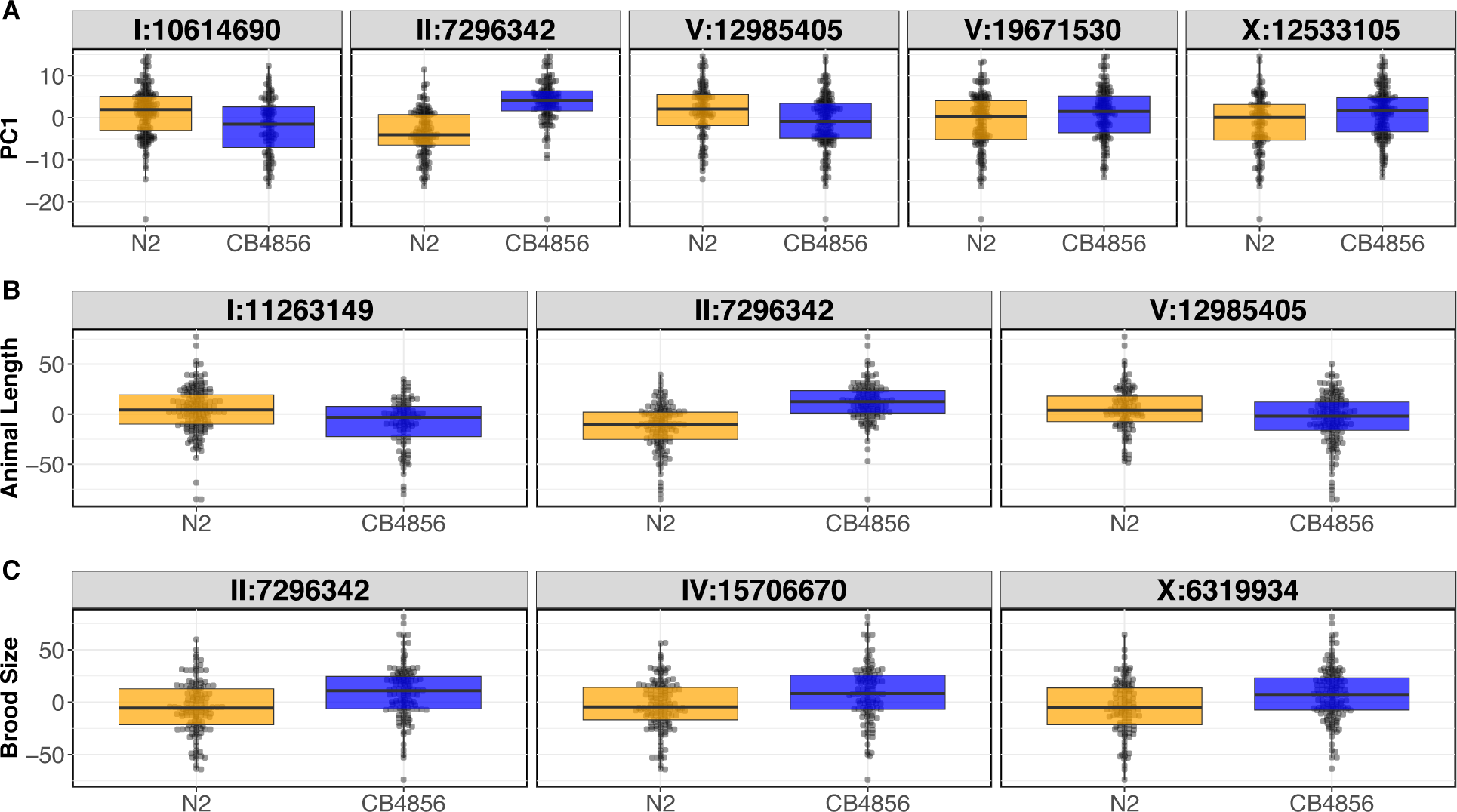
RIAIL phenotypes from the linkage mapping experiment Tukey box plots of the first principal component. (A), assay- and control-regressed brood sizes and (B) median animal lengths of the N2 and CB4856 RIAIL panel after exposure to arsenic trioxide. Each dot corresponds to the phenotype for a single RIAIL. The RIAILs are separated by the N2 (orange) or CB4856 (blue) genotype at each QTL detected by linkage mapping.

**Figure 1-figure supplement 6:**
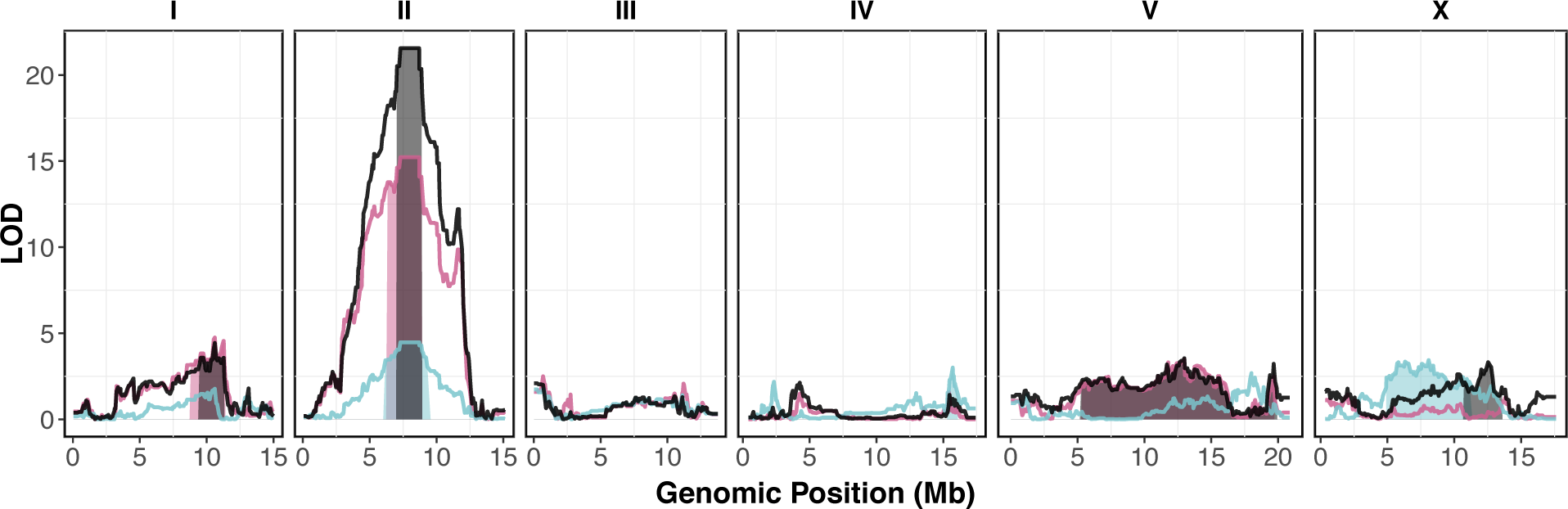
Linkage mapping results for brood size, animal length, and the first principal component. Linkage mapping plots for regressed brood size (teal), median animal length (pink), and the first principal component (black) in the presence of 1000 µM arsenic trioxide are shown. The significance values (logarithm of odds, LOD, ratio) for 1454 markers between the N2 and CB4856 strains are on the y-axis, and the genomic position (Mb) separated by chromosome is plotted on the x-axis. The associated 1.5 LOD-drop confidence intervals are represented by teal, pink, and boxes for brood size, median animal length, and the first principal component respectively.

**Figure 1-figure supplement 7:**
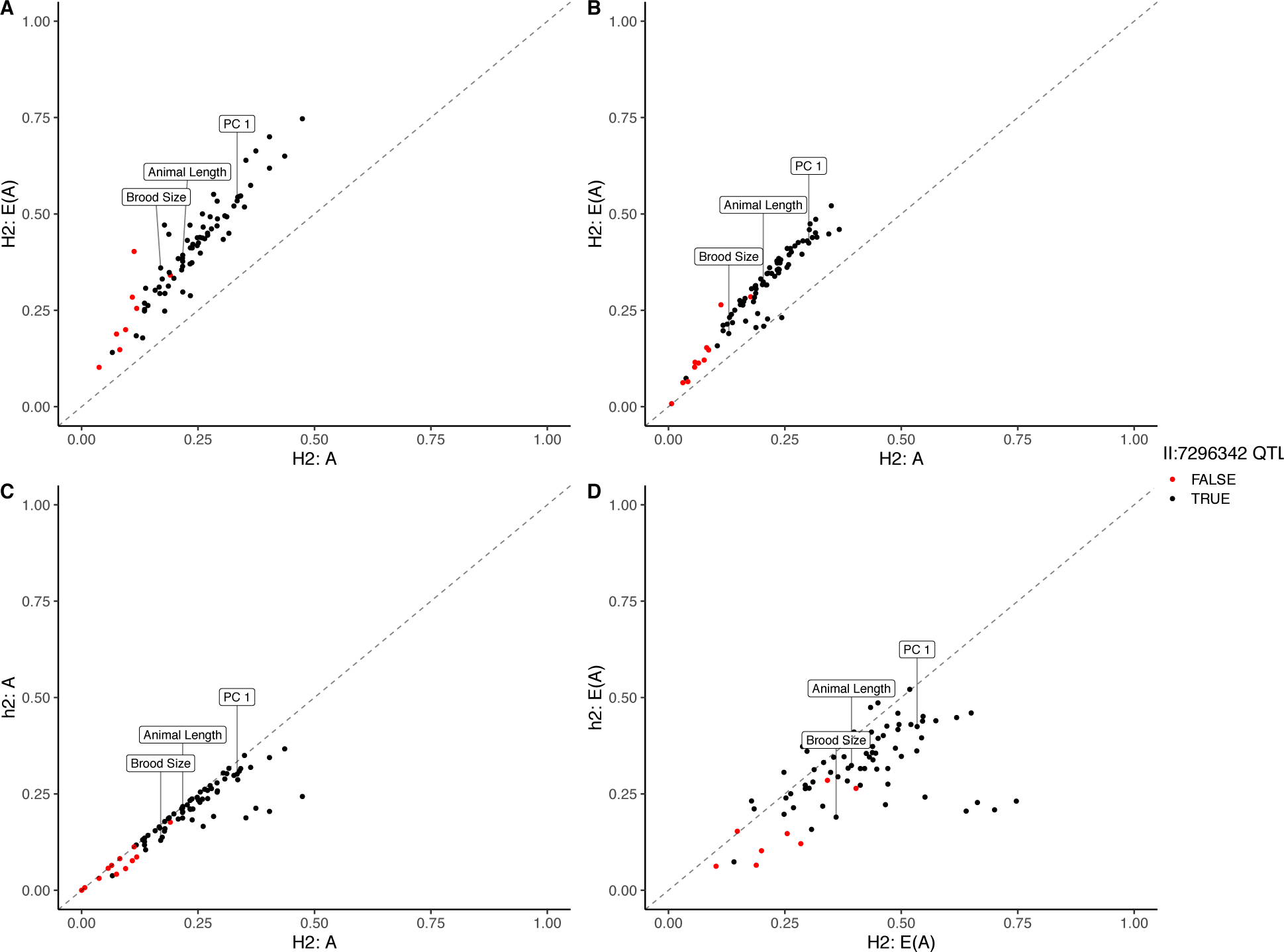
Genomic-heritability estimates of linkage mapping traits. The genomic broad (*H*^2^)- and narrow (*h*^2^)-sense heritability estimates calculated using the expectation (E(A)) of the realized relatedness matrix or the realized relatedness matrix (A). Each dot represents a measured or principal component trait. Dots are colored black if that trait mapped to the center of chromosome II and red if no QTL was detected on the center of chromosome II. The brood size, animal length, and first principal component traits are marked for clarity.

**Figure 1-figure supplement 8:**
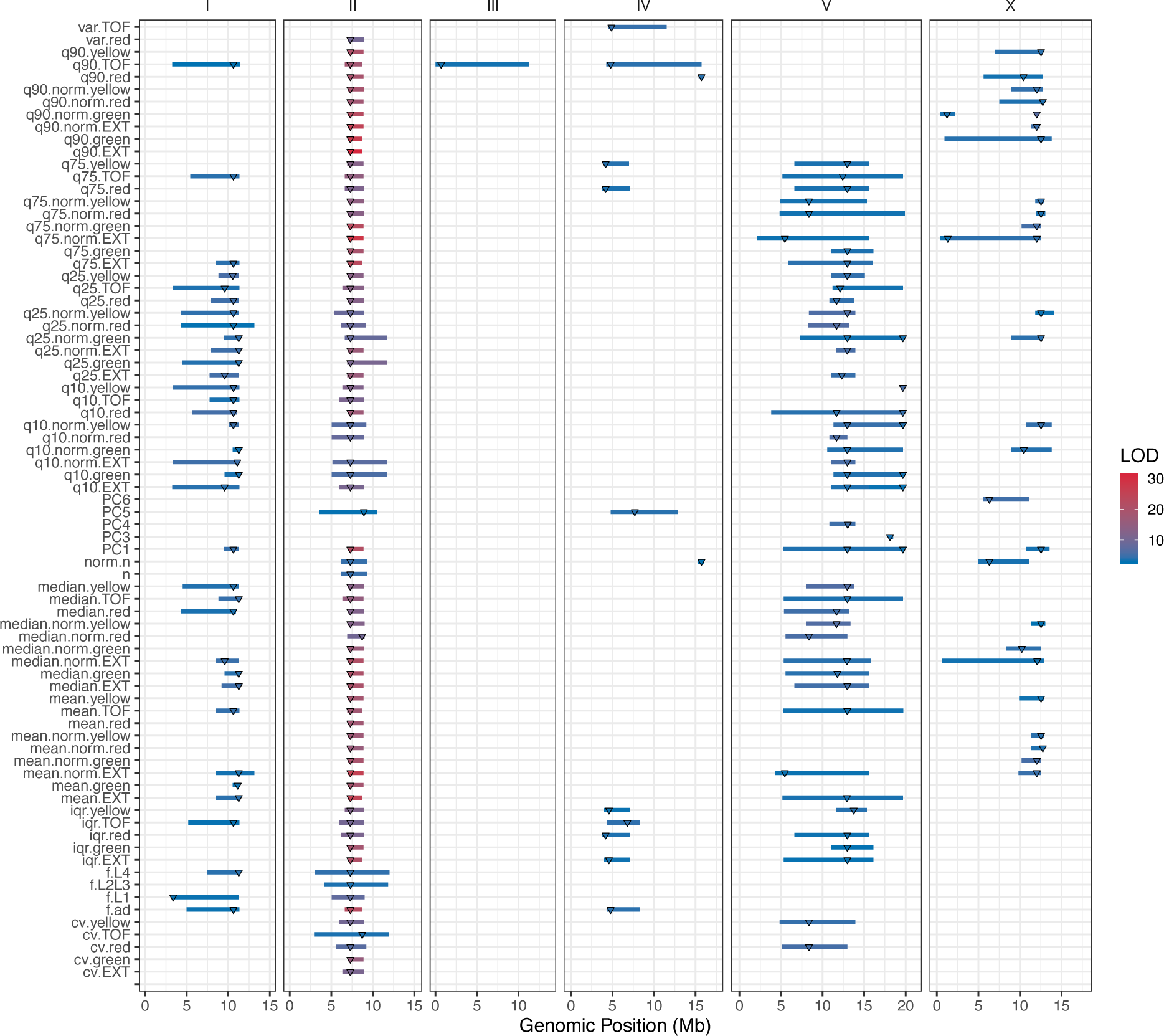
Linkage mapping QTL summary. All QTL identified by linkage mapping are shown. Traits are labeled on the y-axis and the genomic position in Mb is plotted on the x-axis. Triangles represent the peak QTL position and bars represent the associated 1.5-LOD drop QTL confidence interval. Triangles and bars are colored based on the associated LOD score, where red colors correspond to higher LOD values.

**Figure 1-figure supplement 9:**
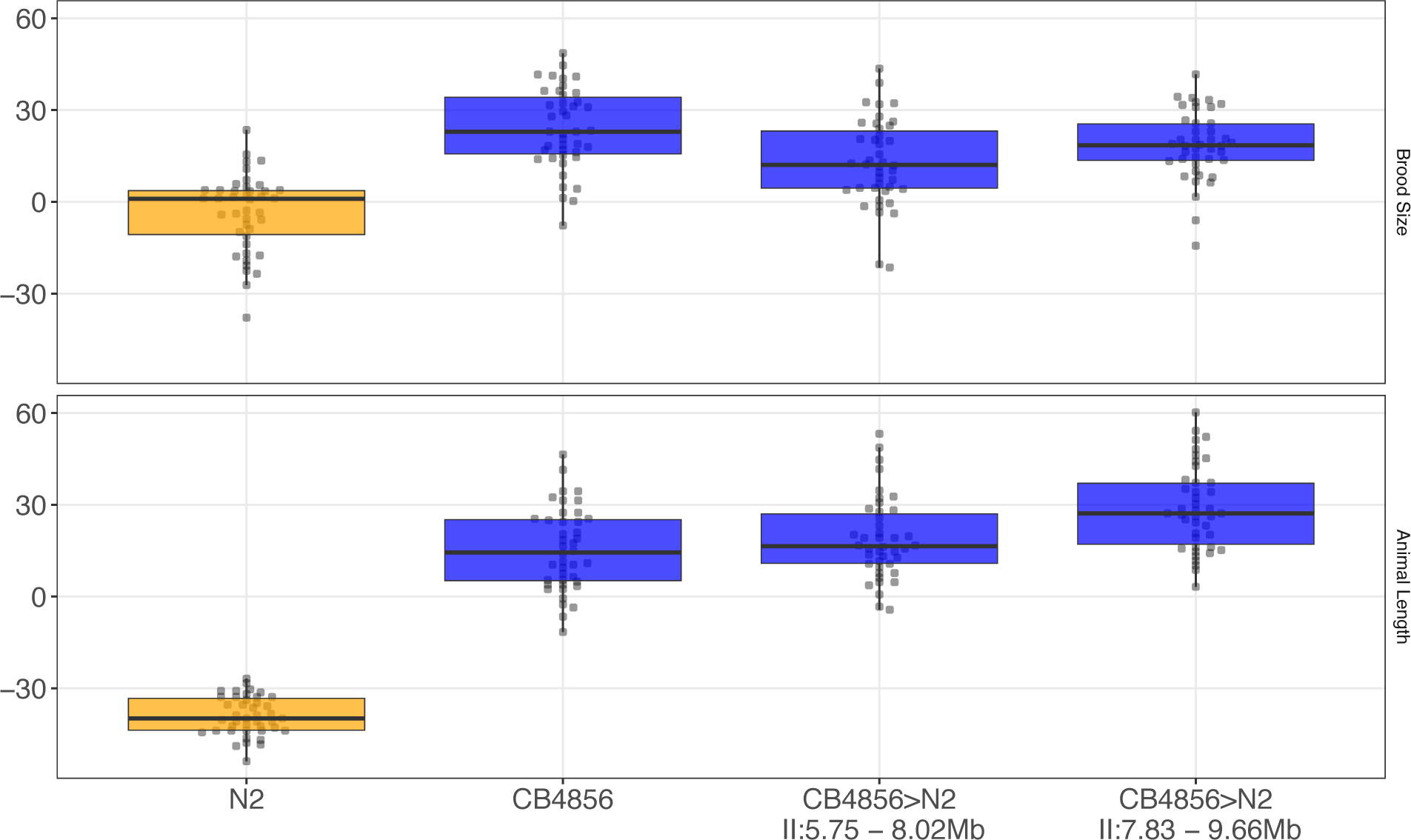
NIL recapitulation of chromosome II QTL. Tukey box plots of near-isogenic line (NIL) phenotype values for the brood size (top panel) and animal length (bottom panel) traits in the presence of 1000 µM arsenic trioxide are shown. NIL genotypes are indicated below the plot as genomic ranges. The N2 brood sizes and progeny lengths are significantly different from all other strains (Brood size: Tukey HSD *p*-value < 1.56E-7; Animal Length: Tukey HSD *p*-value < 3.0E-14).

**Figure 1-figure supplement 10:**
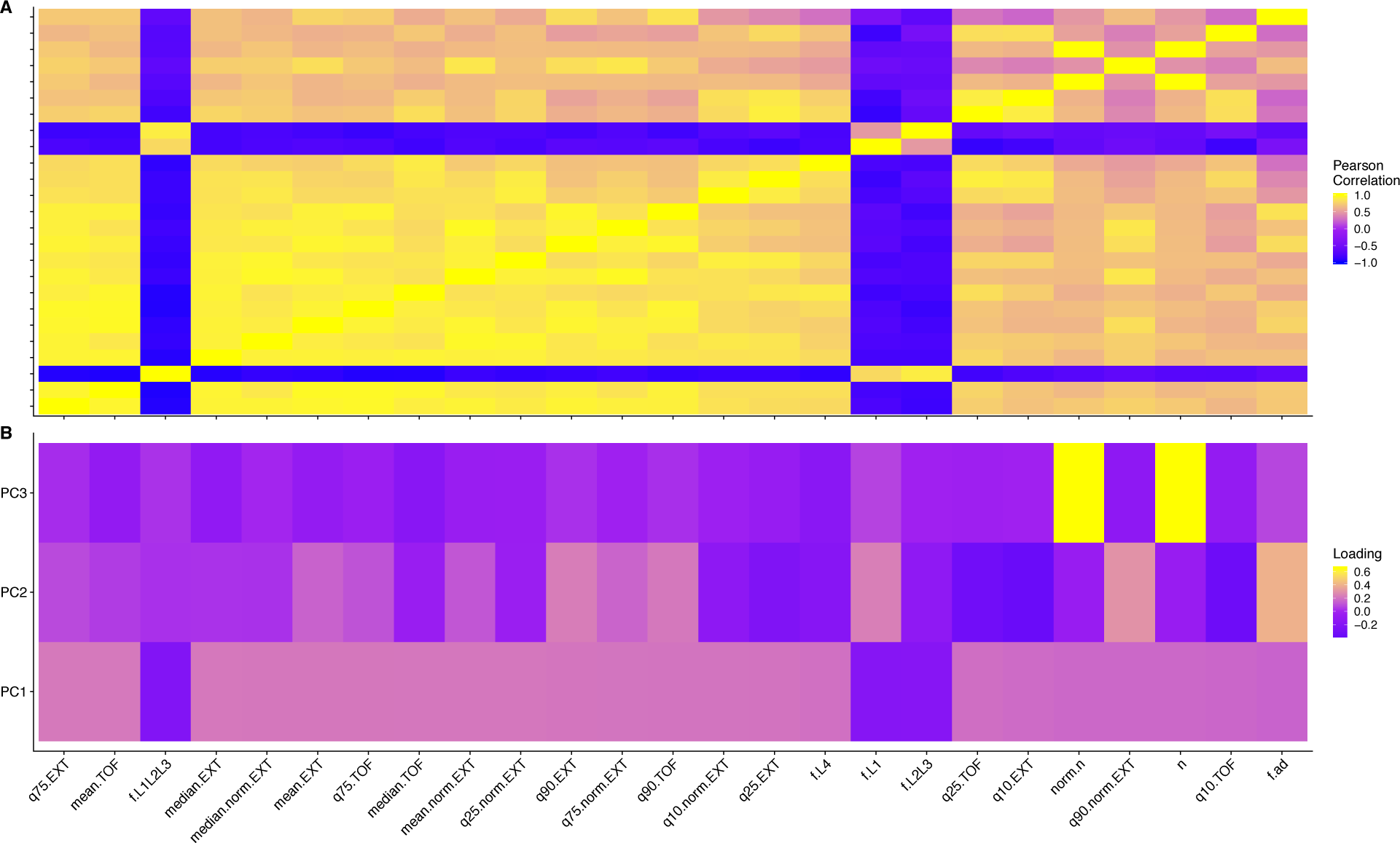
Trait correlations and principal component loadings of NIL and allele-swap recapitulation experiment. (A) The trait Pearson’s correlation coefficient of the assay- and control-regressed measured traits. (B) The contribution of each measured trait to the principal components that explain 90% of the total variance in the NIL and allele swap-recapitulation experiment, which was performed at 1000 µM. For each plot, the tile color corresponds to the value, where yellow colors represent higher values.

**Figure 1-figure supplement 11:**
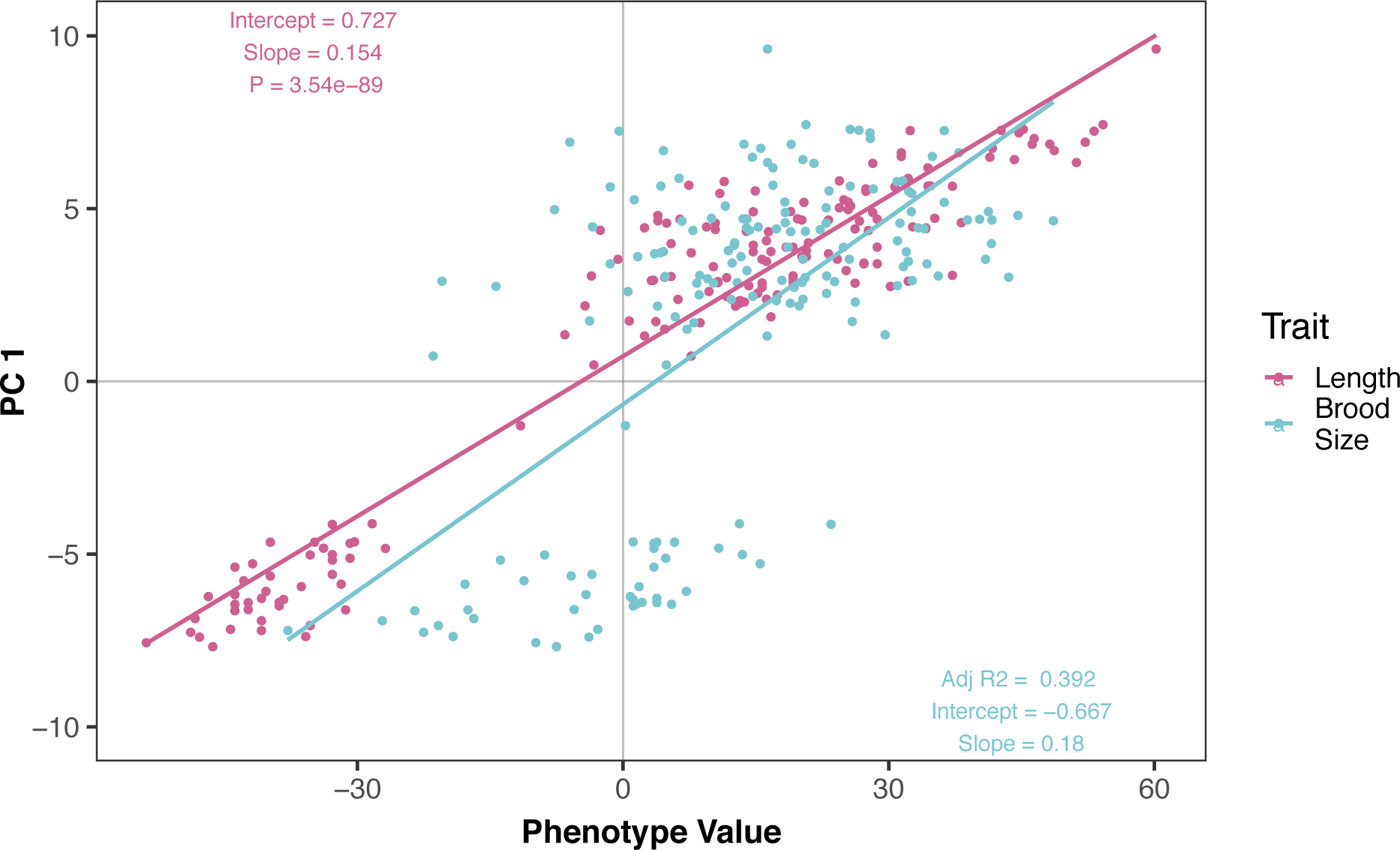
Brood size and animal length are correlated with the first principal component for the NIL recapitulation experiment. The correlation between brood size (blue) or animal length (pink) with the first principal component trait for the NIL recapitulation experiment. Each dot represents an individual NIL or parental strain replicate phenotype, with the animal length and brood size phenotype values on the x-axis and the first principal component phenotype on the y-axis.

**Figure 2-figure supplement 1:**
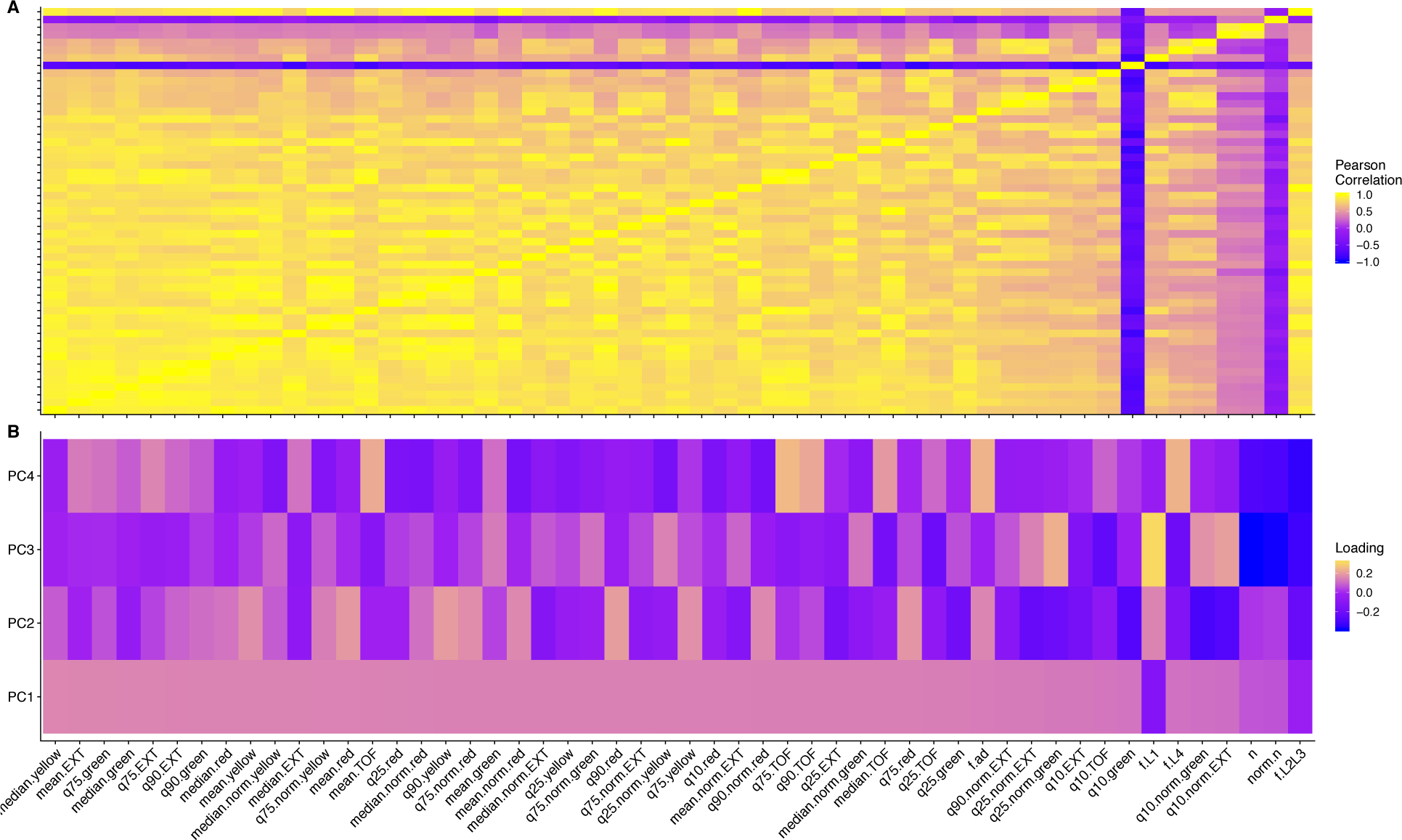
Trait correlations and principal component loadings of GWA mapping experiment. A) The trait Pearson’s correlation coefficient of the assay- and control-regressed Measured traits. (B) The contribution of each measured traits to the principal components that explain 90% of the total variance in the GWA mapping experiment, which was performed at 1000 µM. For each plot, the tile color corresponds to the value, where yellow colors represent higher values.

**Figure 2-figure supplement 2:**
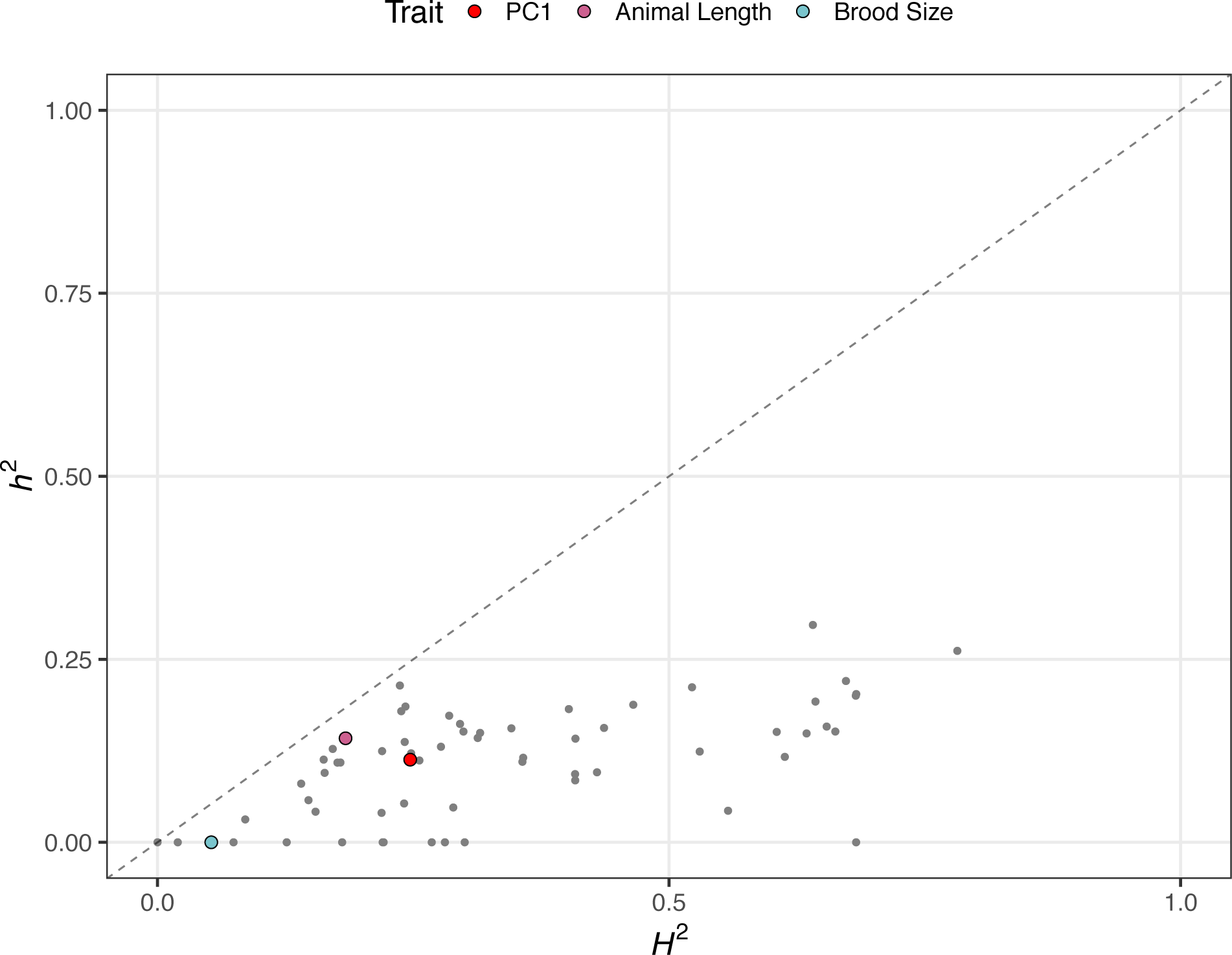
Genomic-heritability estimates of GWA mapping traits. The genomic broad (*H*^2^)- and narrow (*h*^2^)-sense heritability estimates calculated using the the realized relatedness matrix. Each dot represents a measured or principal component trait. The three traits discussed throughout the manuscript are highlighted — red: first principal component, pink: animal length, and blue: brood size.

**Figure 2-figure supplement 3:**
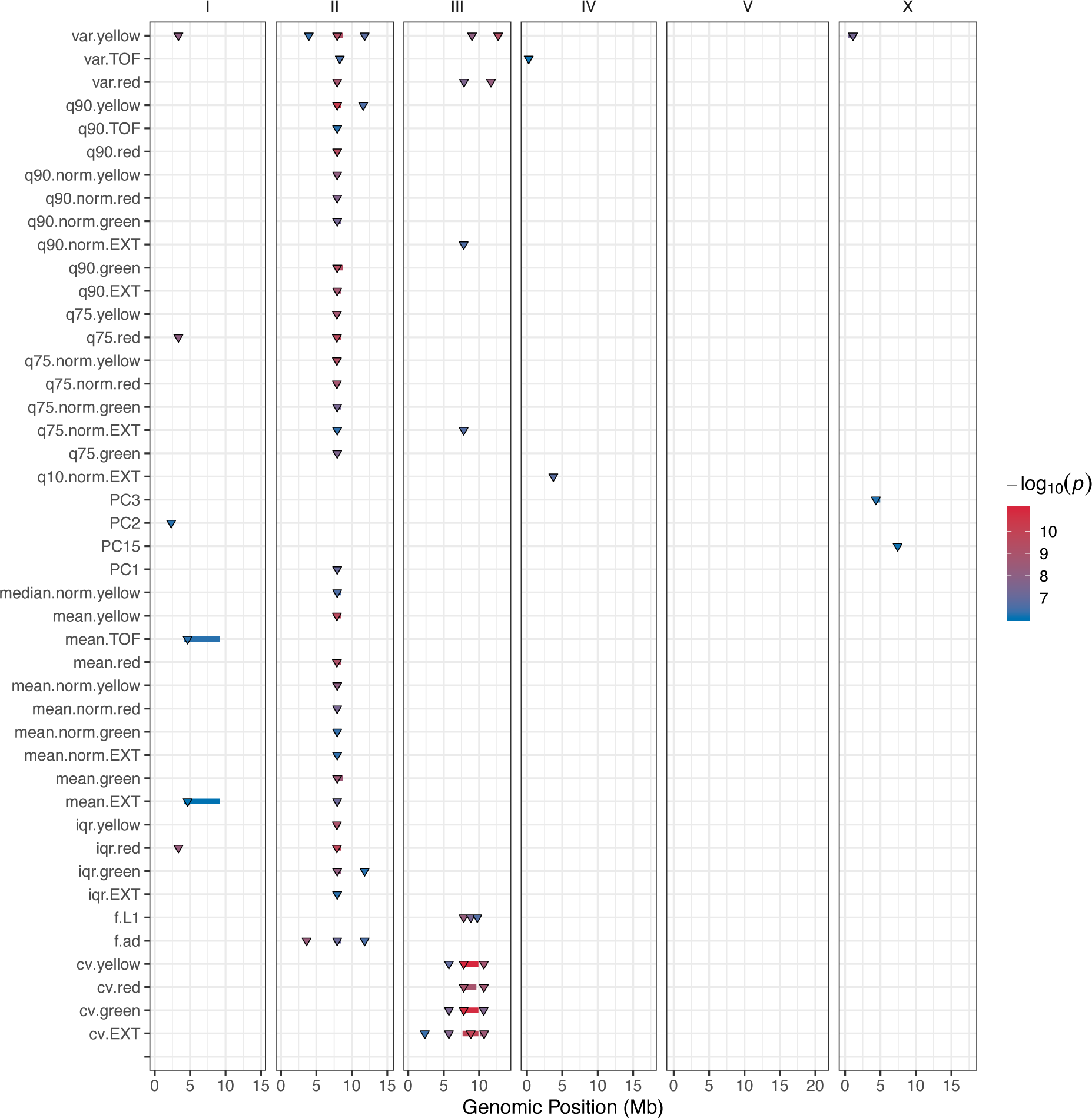
GWA mapping QTL summary. All QTL identified by GWA mapping are shown. Traits are labeled on the y-axis and the genomic position in Mb is plotted on the x-axis. Triangles represent the peak QTL position and bars represent the associated QTL region of interest. Triangles and bars are colored based on the associated significance value, where red colors correspond to higher significance values.

**Figure 2-figure supplement 4:**
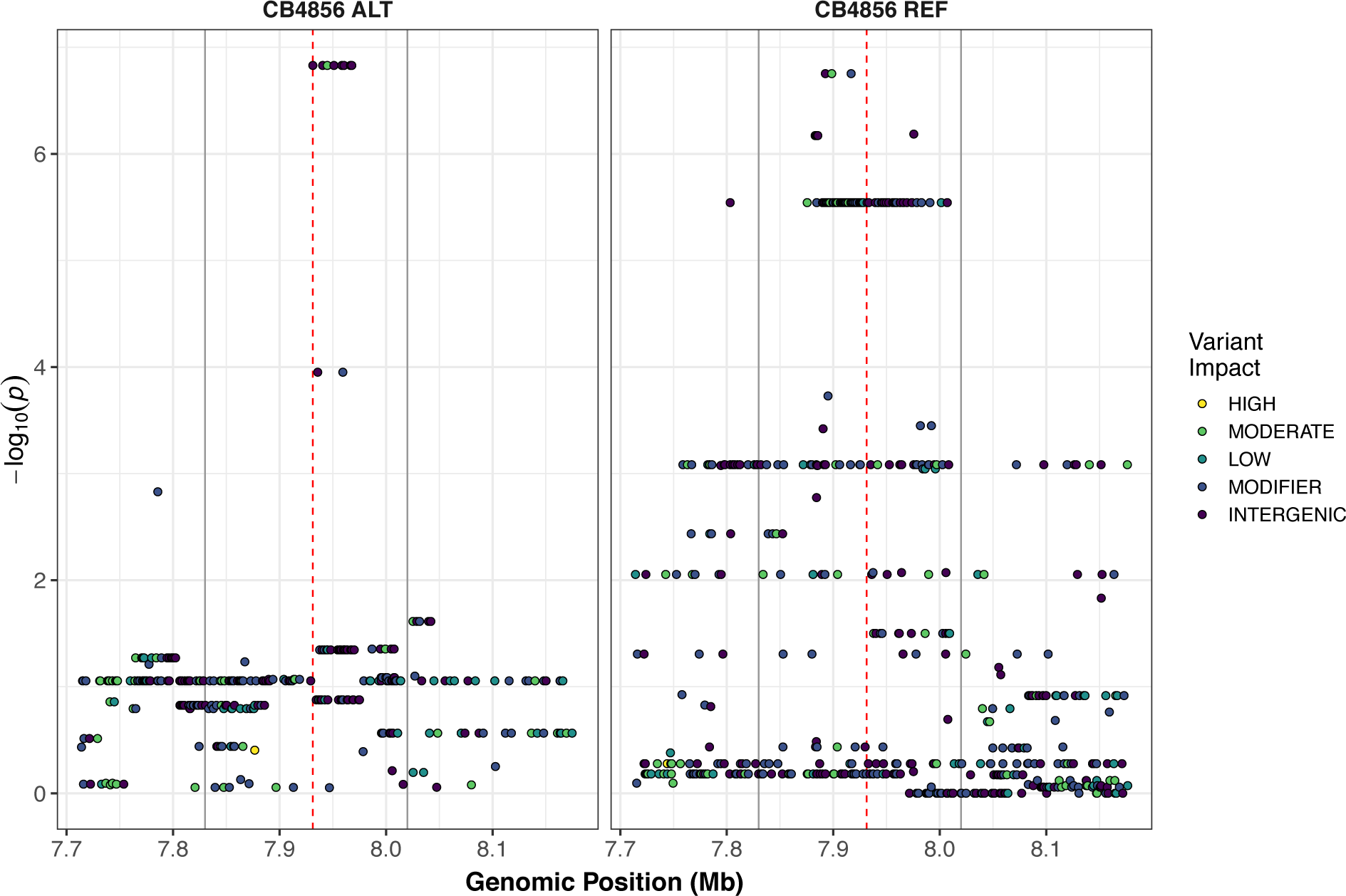
Fine-mapping of the chromosome II QTL identified by GWA mapping. Fine mapping of the chromosome II region of interest (cyan region from panel A, 7.71 - 8.18 Mb) is shown. Each dot represents an SNV present in the phenotyped population. SNVs present in the CB4856 strain are shown in the left panel and SNVs present in other phenotyped strains, but REF in CB4856, are shown in the right panel. The association between the SNV and first principal component is shown on the y-axis and the genomic position of the SNV is shown on the x-axis. Dots are colored by their SnpEff predicted effect.

**Figure 3-figure supplement 1:**
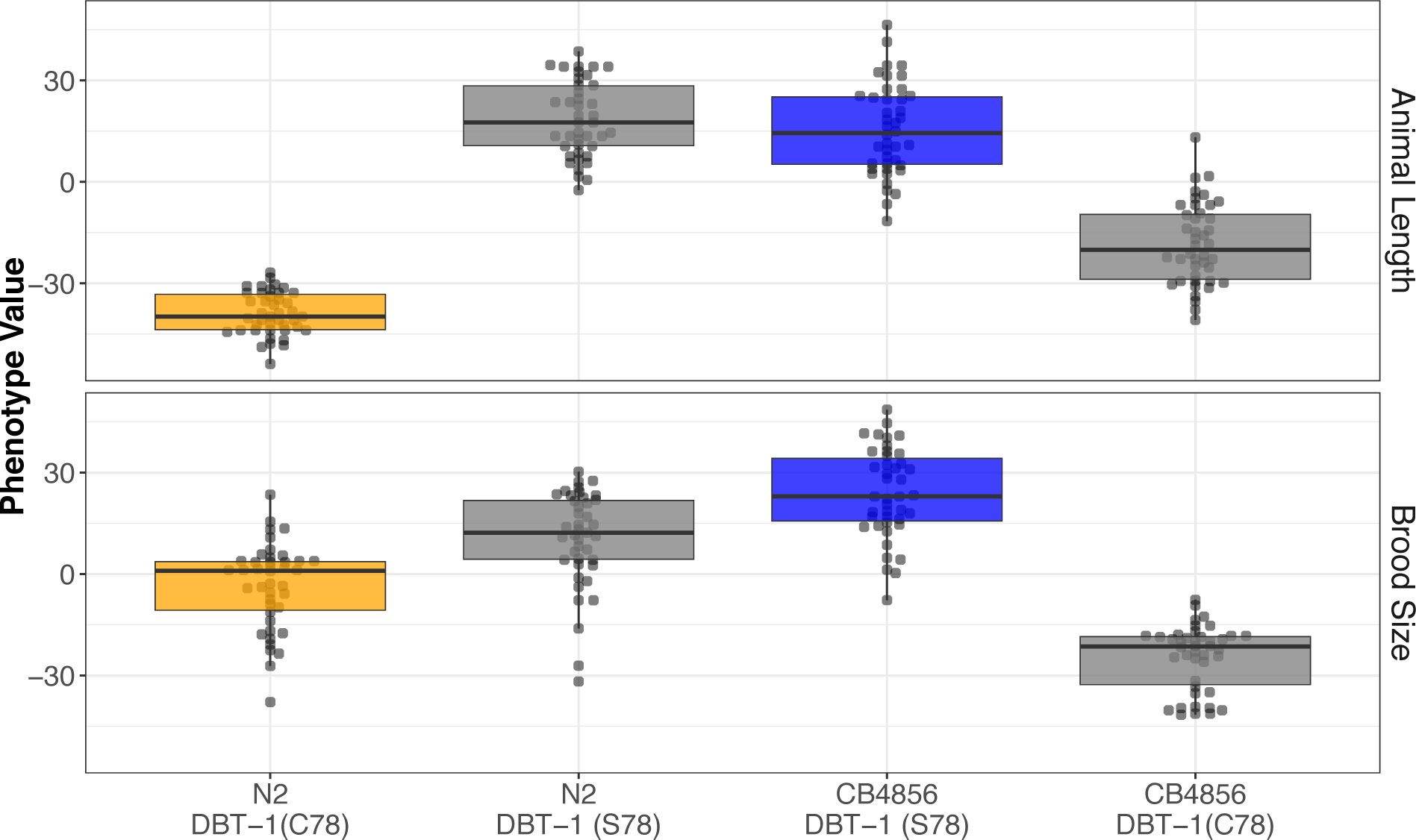
The DBT-1 C78S variant underlies arsenic trioxide sensitivity in *C. elegans.* Tukey box plots of residual animal length (top panel) and median animal length (bottom panel) after exposure to 1000 µM arsenic trioxide are shown (N2, orange; CB4856, blue; allele replacement strains, gray). Labels correspond to the genetic background and the corresponding residue at position 78 of DBT-1 (C for cysteine, S for serine). Every pair-wise strain comparison is significant except for the N2 DBT-1(S78) - CB4856 comparison from animal length (Brood size: Tukey’s HSD *p*-value < 1.55E-5; Animal length: *p*-value << 1E-7).

**Figure 3-figure supplement 2:**
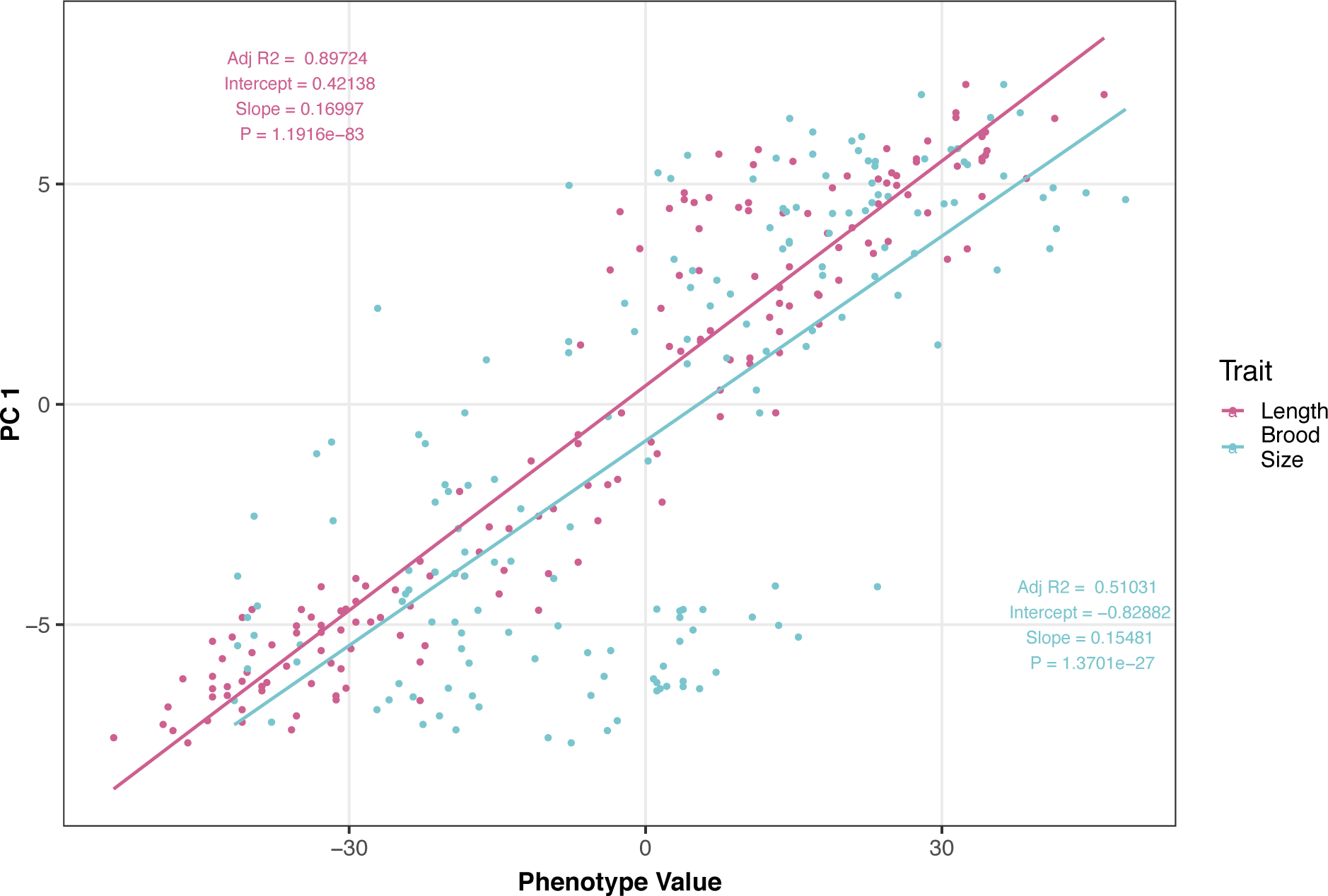
Brood size and animal length are correlated with the first principal component for the allele-swap recapitulation experiment. The correlation between brood size (blue) or animal length (pink) with the first principal component trait for the allele-swap recapitulation experiment. Each dot represents an individual allele-swap or parental strain replicate phenotype, with the animal length and brood size phenotype values on the x-axis and the first principal component phenotype on the y-axis.

**Figure 4-figure supplement 1:**
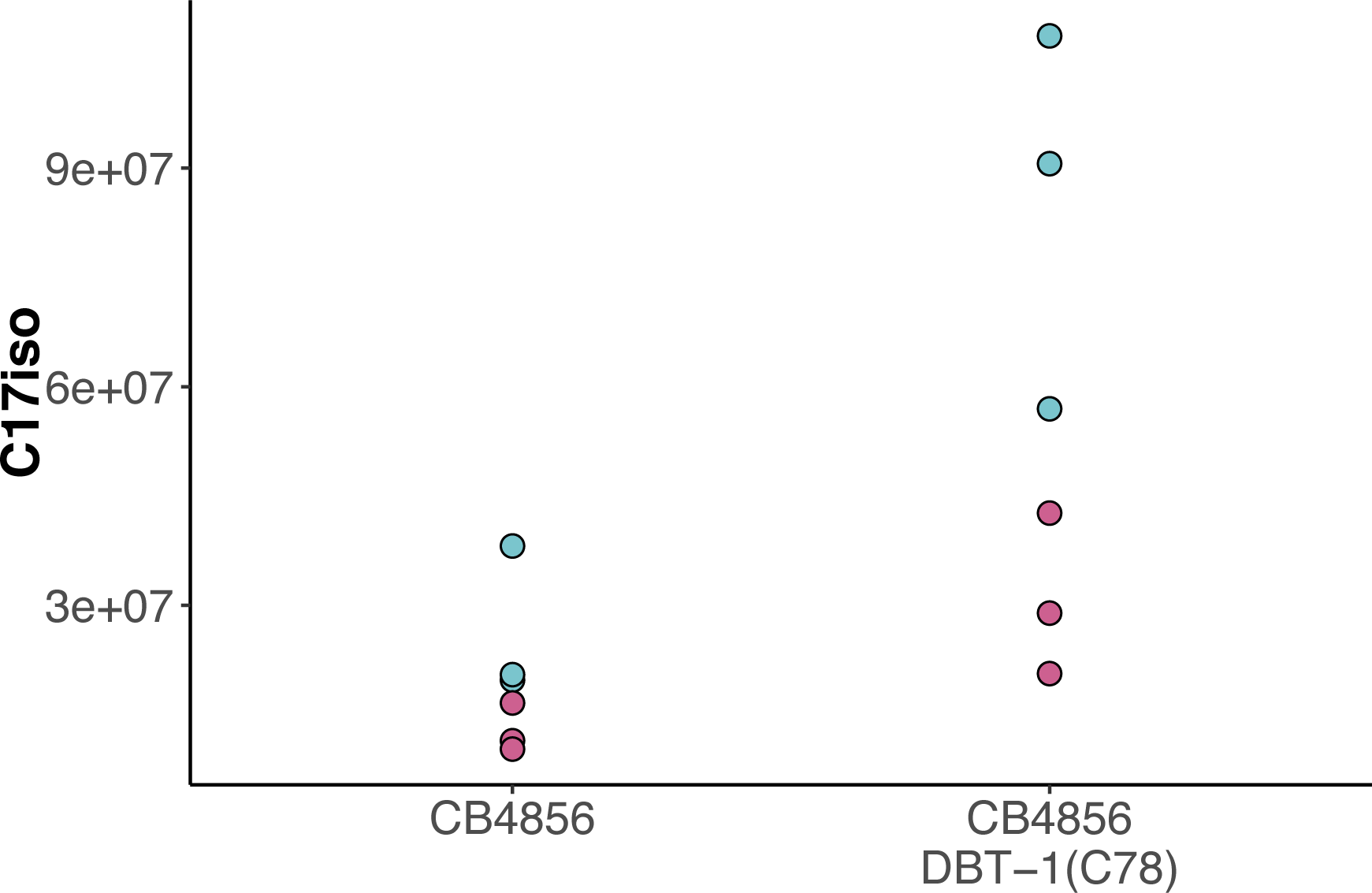
Raw abundance of C17ISO for CB4856 and CB4856 allele swap. The raw abundance of C17ISO is plotted on the y-axis for three independent replicates of the CB4856 and CB4856 allele swap strains exposed to control (teal) or 100 µM arsenic trioxide (pink) conditions. The difference between CB4856 swap mock and arsenic conditions is significant (Tukey HSD *p*-value = 0.029), but the difference between CB4856 mock and arsenic conditions is not (Tukey HSD *p*-value = 0.10).

**Figure 4-figure supplement 2:**
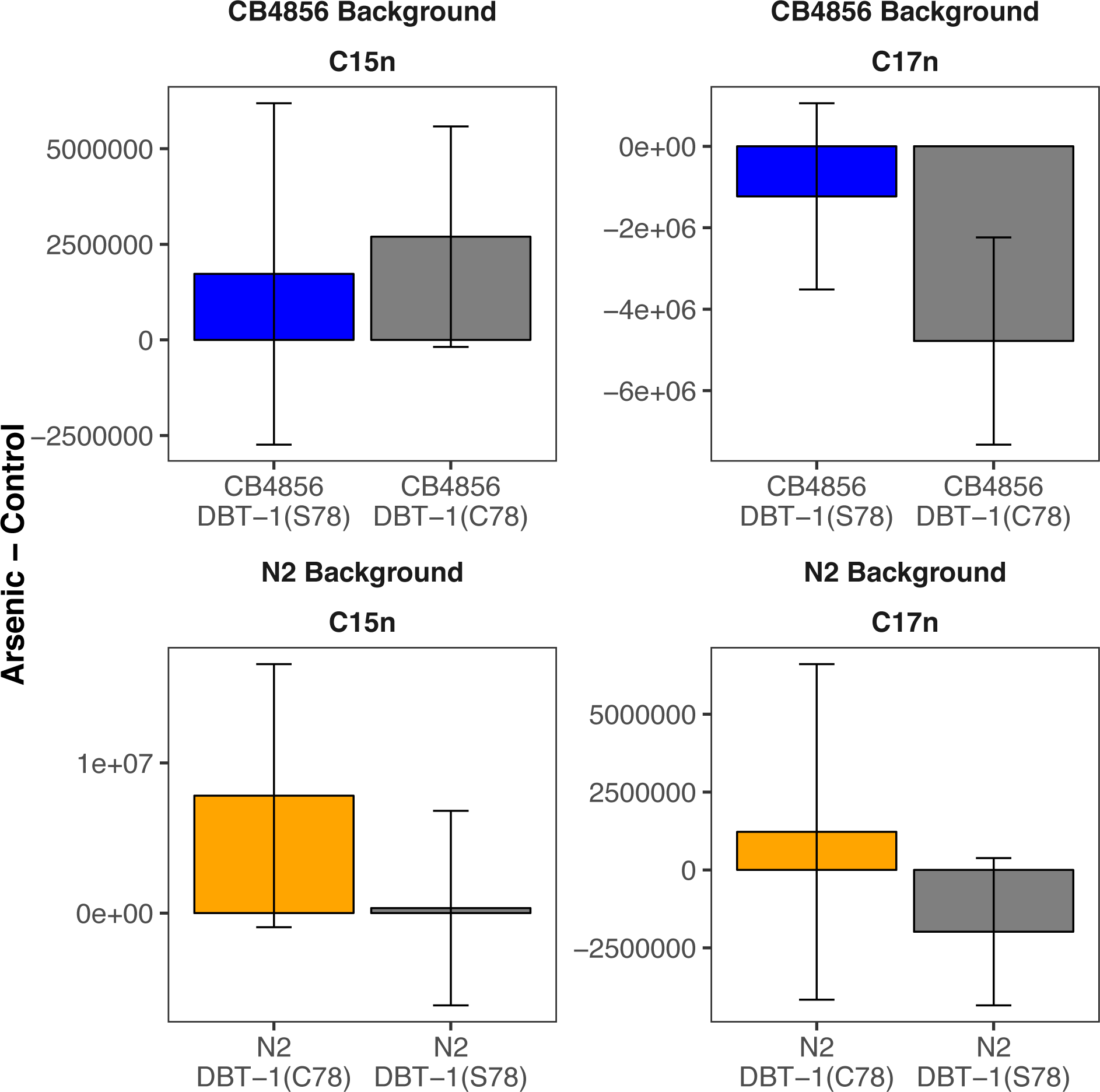
Straight-chain fatty acids are not affected by arsenic trioxide. The difference in raw C15SC (left panel) or C17SC (right panel) abundances between 100 µM arsenic trioxide and control conditions is plotted on the y-axis for three independent replicates of the CB4856 and CB4856 allele swap strains and six independent replicates of the N2 and N2 allele swap strains. There are no significant differences when comparing the abundances between parental and allele-swap strains.

**Figure 4-figure supplement 3:**
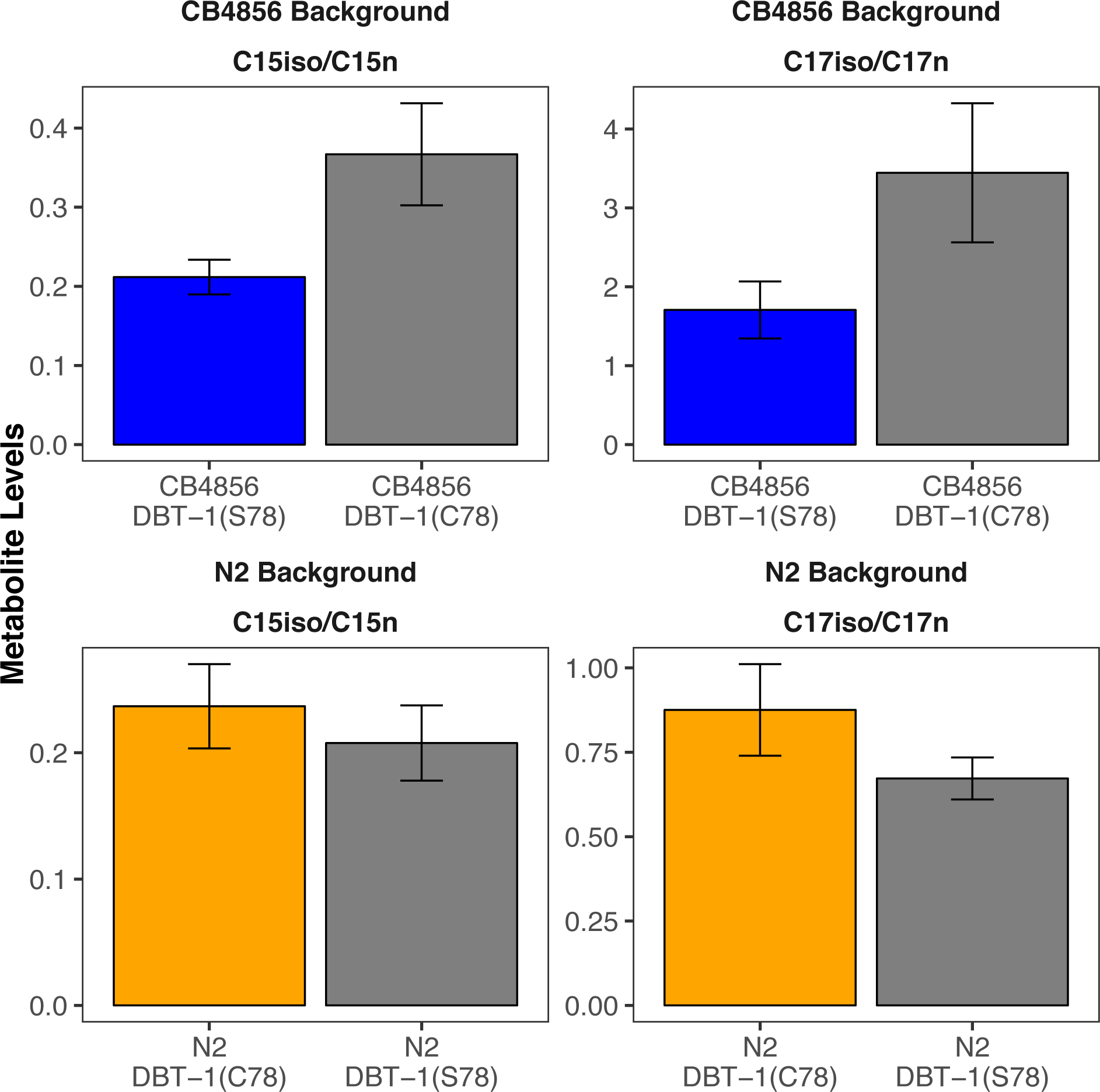
C15ISO and C17ISO to strait-chain ratios in control conditions. The C15ISO/C15SC (left panel) or C17ISO/C17SC (right panel) ratios in control conditions are plotted on the y-axis for three independent replicates of the CB4856 and CB4856 allele swap strains and six independent replicates of the N2 and N2 allele swap strains. The C15 and C17 ratios for the CB4856-CB4856 allele swap comparison are significant (C15: Tukey HSD *p-*value = 0.0168749; C17: Tukey HSD *p-*value = 0.0342525). The difference between the C17 ratio for the N2-N2 allele swap comparison is significant (Tukey HSD *p-*value = 0.0044667), but the difference in the C15 ratio is not significant (Tukey HSD *p-*value = 0.1239674).

**Figure 4-figure supplement 4:**
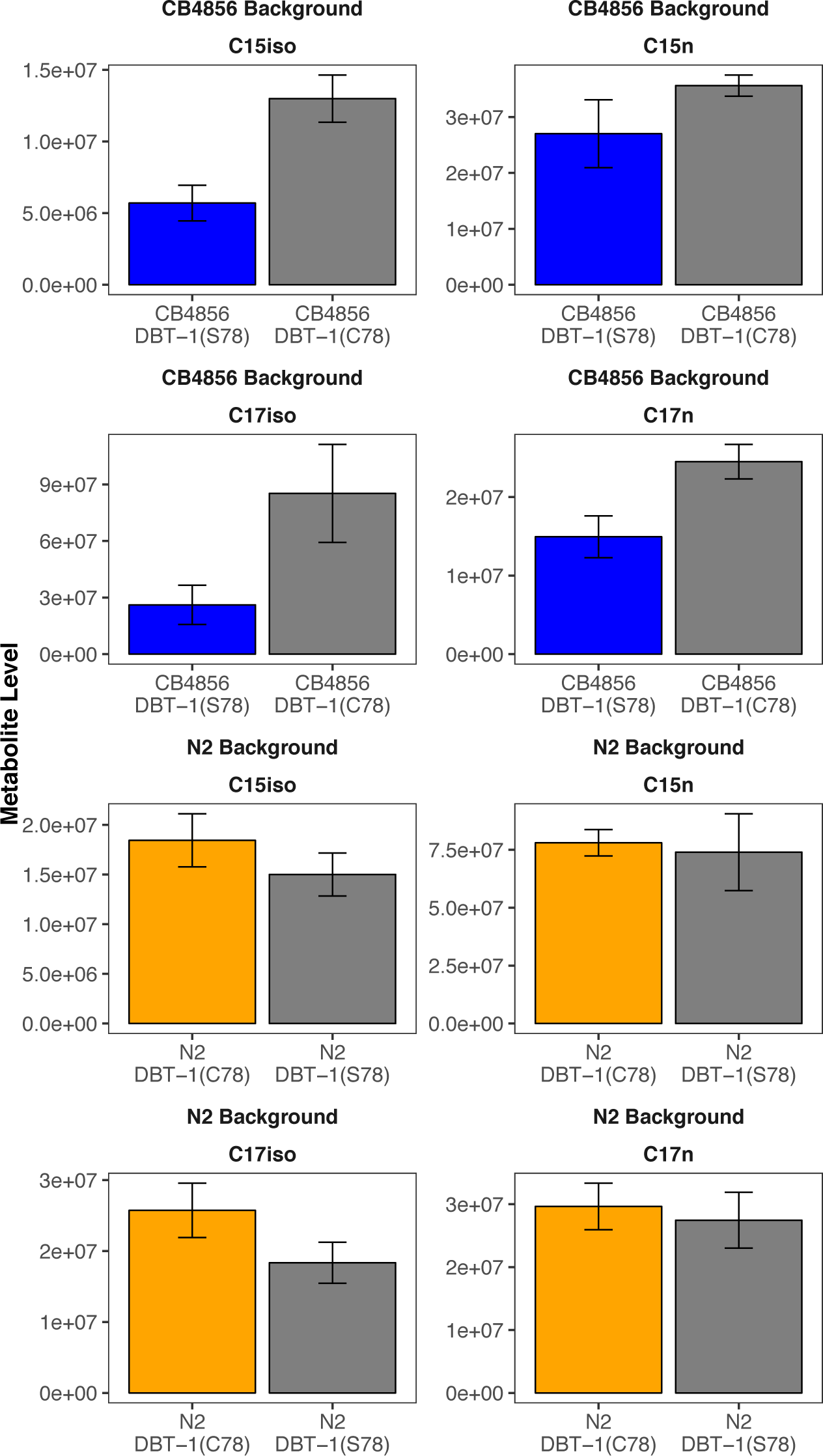
Strains with the DBT-1(C78) allele produce more branched chain fatty acids in the L1 larval stage in control conditions. Branched chain (left panel) and straight chain (right panel) fatty acid measurements in L1 animals are represented on the y-axis. There are significant differences in abundances when comparing all parental and allele-swap strains for C15ISO and C17ISO chain fatty acids (CB4856-C15ISO DBT-1(C78): Tukey HSD *p-*value = 0.0036201, n = 3; N2-C15ISO DBT-1(C78): Tukey HSD *p-*value = 0.0265059, n = 6; CB4856-C17ISO DBT-1(C78): Tukey HSD *p-*value = 0.0086572, n = 3; N2-C17ISO DBT-1(C78): Tukey HSD *p-*value = 0.0022501, n = 6). Conversely, we observe no significant differences in straight chain fatty production except for C17n production in the CB4856 background (CB4856-C15n DBT-1(C78): Tukey HSD *p-*value = 0.0787388, n = 3; N2-C15n DBT-1(C78): Tukey HSD *p-*value = 0.5817993, n = 6; CB4856-C17n DBT-1(C78): Tukey HSD *p-*value = 0.0086572, n = 3; N2-C17n DBT-1(C78): Tukey HSD *p-*value = 0.35827, n = 6).

**Figure 4-figure supplement 5:**
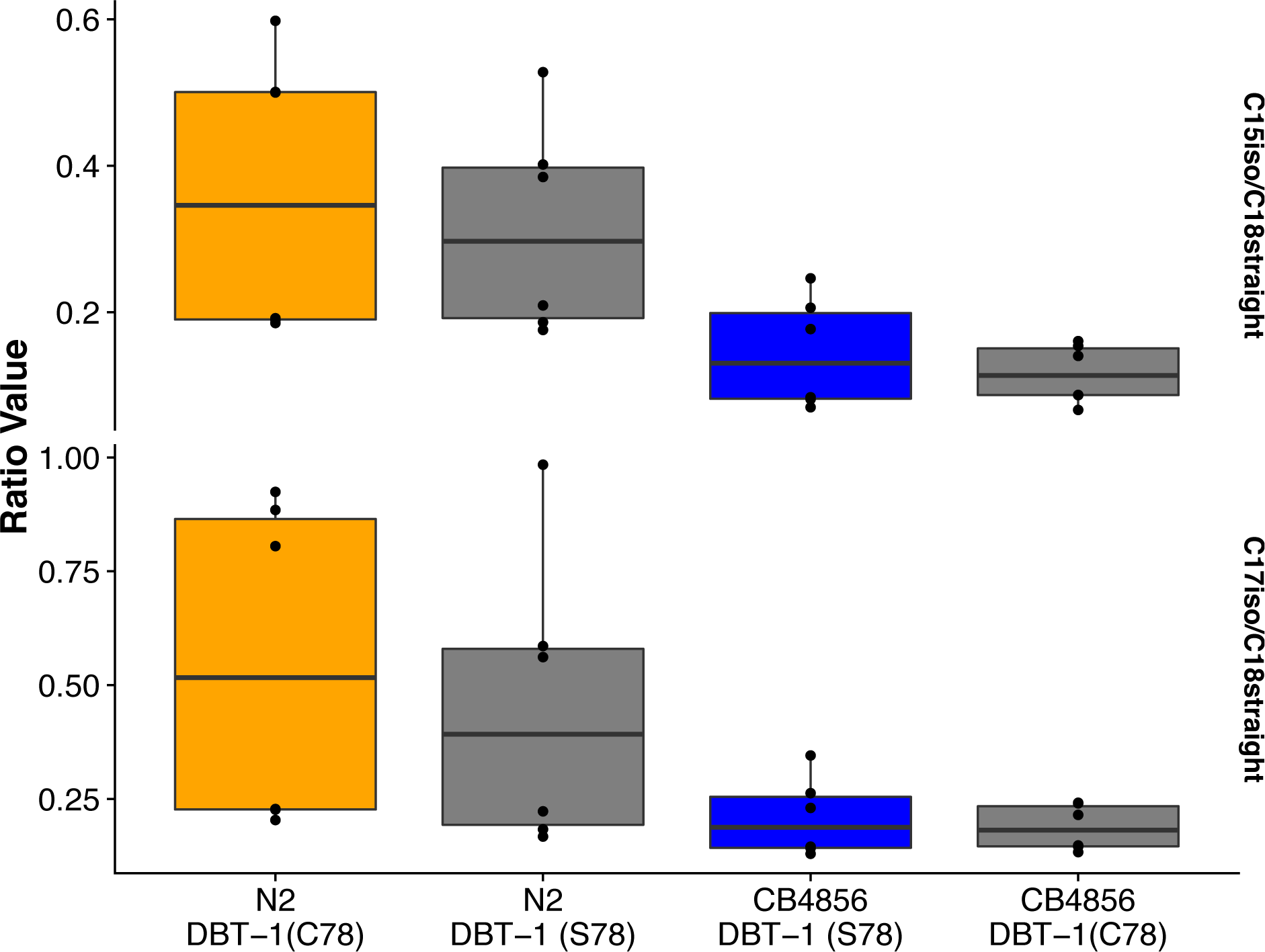
Young adult C15ISO and C17ISO to strait-chain ratios in control conditions. The C15ISO/C18SC (top panel) or C17ISO/C18SC(bottom panel) ratios of Young adult animals in control conditions are plotted on the y-axis for six independent replicates for all strains. There are no significant differences when comparing the abundances between parental and allele-swap strains.

**Figure 4-figure supplement 6:**
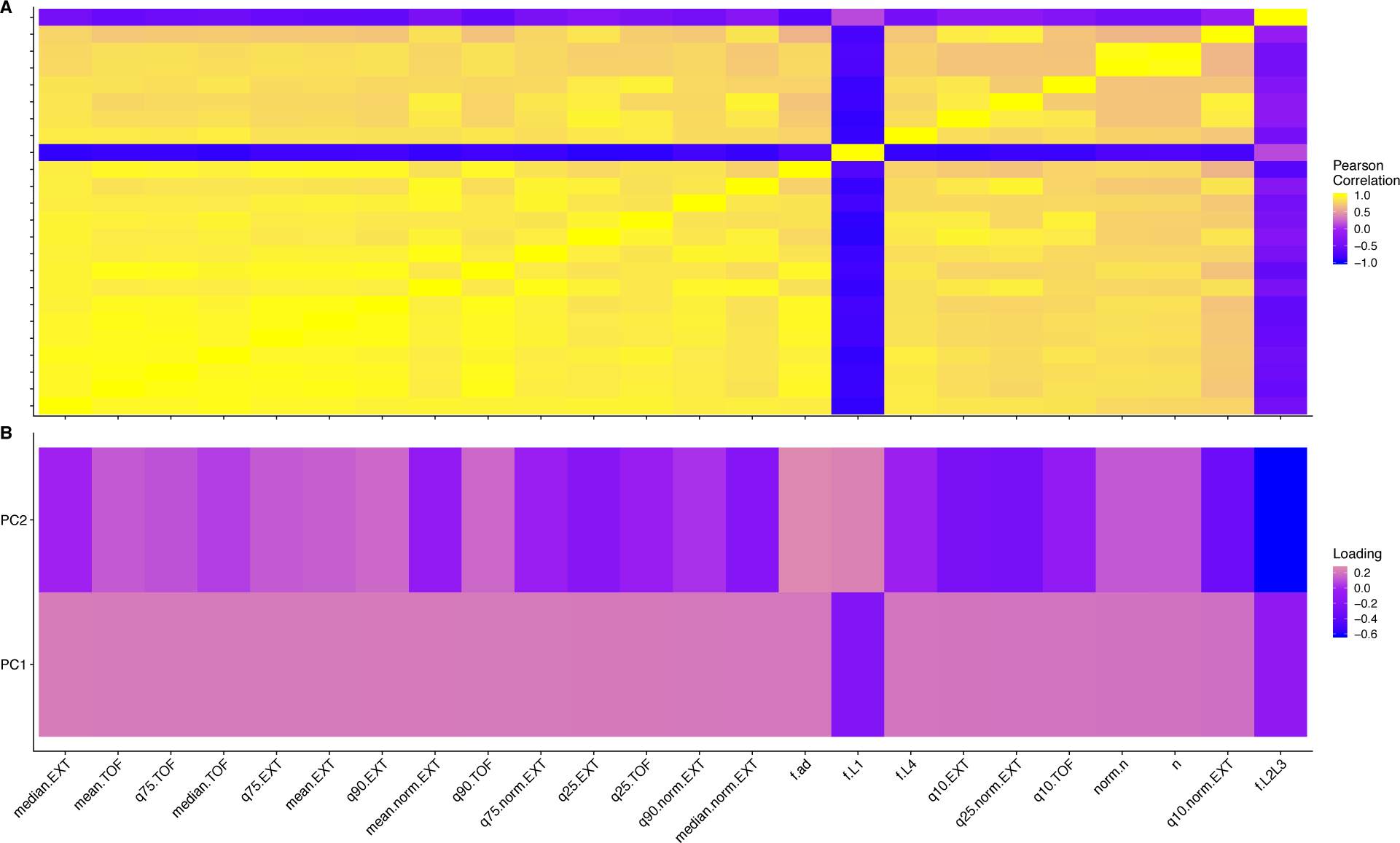
Trait correlations and principal component loadings of C15ISO rescue experiment. (A) The trait Pearson’s correlation coefficient of the assay- and control-regressed measured traits. (B) The contribution of each measured trait to the principal components that explain 90% of the total variance in the GWA mapping experiment, which was performed at 1000 µM. For each plot, the tile color corresponds to the value, where yellow colors represent higher values.

**Figure 4-figure supplement 7:**
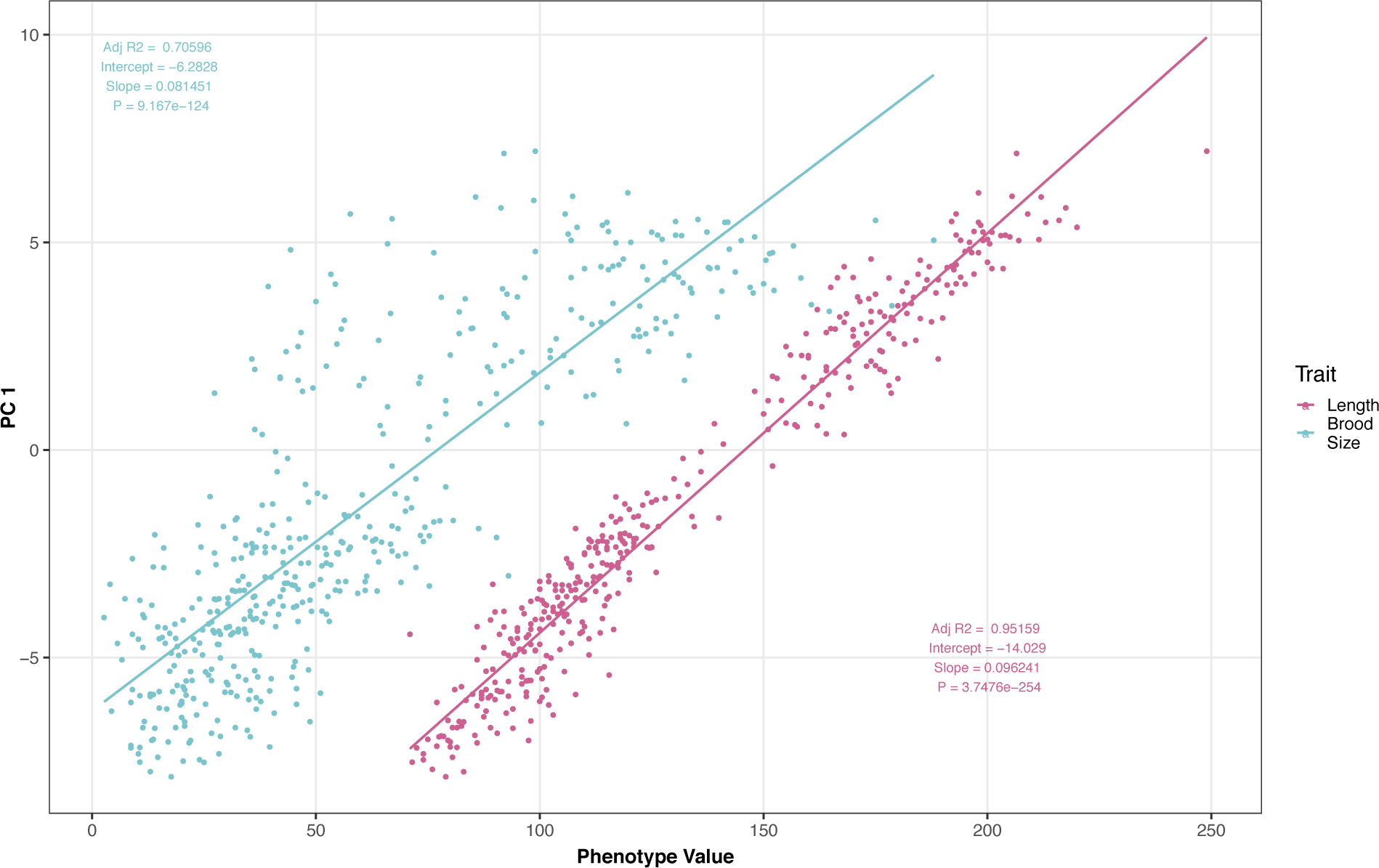
Brood size and animal length are correlated with the first principal component for the C15ISO rescue experiment. The correlation between brood size (blue) or animal length (pink) with the first principal component trait for the C15ISO rescue experiment. Each dot represents an individual allele-swap or parental strain replicate phenotype, with the animal length and brood size phenotype values on the x-axis and the first principal component phenotype on the y-axis.

**Figure 4-figure supplement 8:**
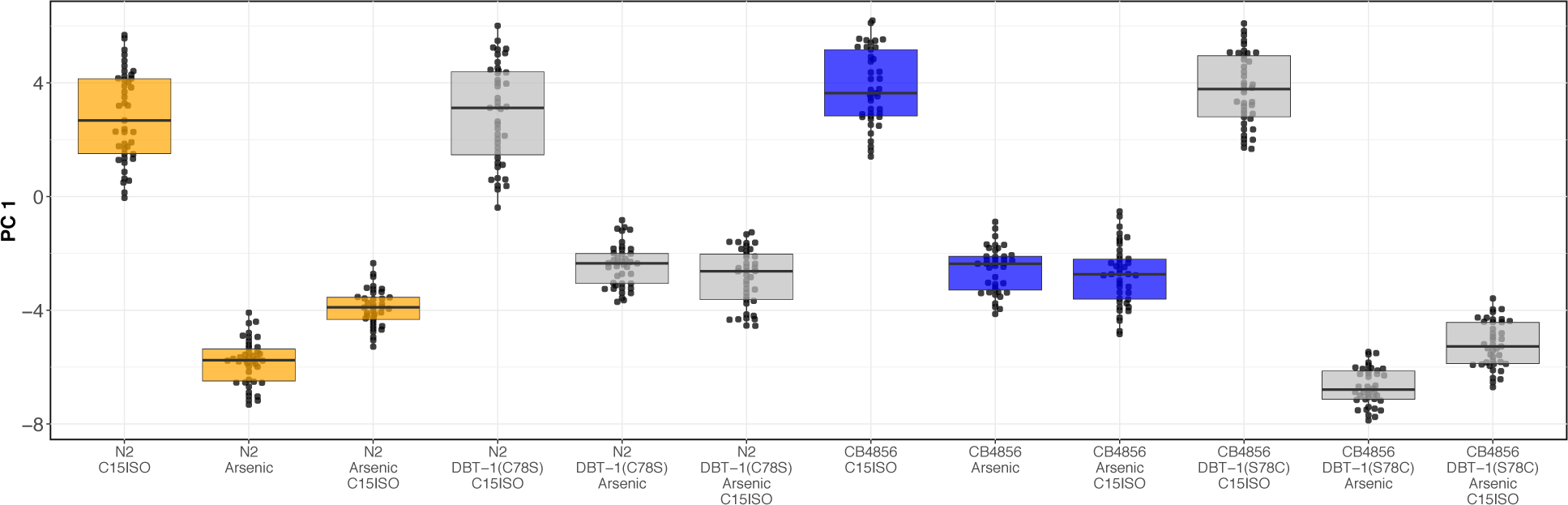
Complete results from C15iso rescue experiment. Tukey box plots median animal length after C15ISO, arsenic trioxide, or arsenic trioxide and 0.64 µM C15ISO exposure are shown (N2, orange; CB4856, blue; allele replacement strains, gray). Labels correspond to the genetic background and the corresponding residue at position 78 of DBT-1 (C for cysteine, S for serine). Every pairwise strain comparison is significant except for the N2 DBT-1(S78) - CB4856 comparisons (Tukey’s HSD *p*-value < 1.43E-6).

**Figure 5-figure supplement 1:**
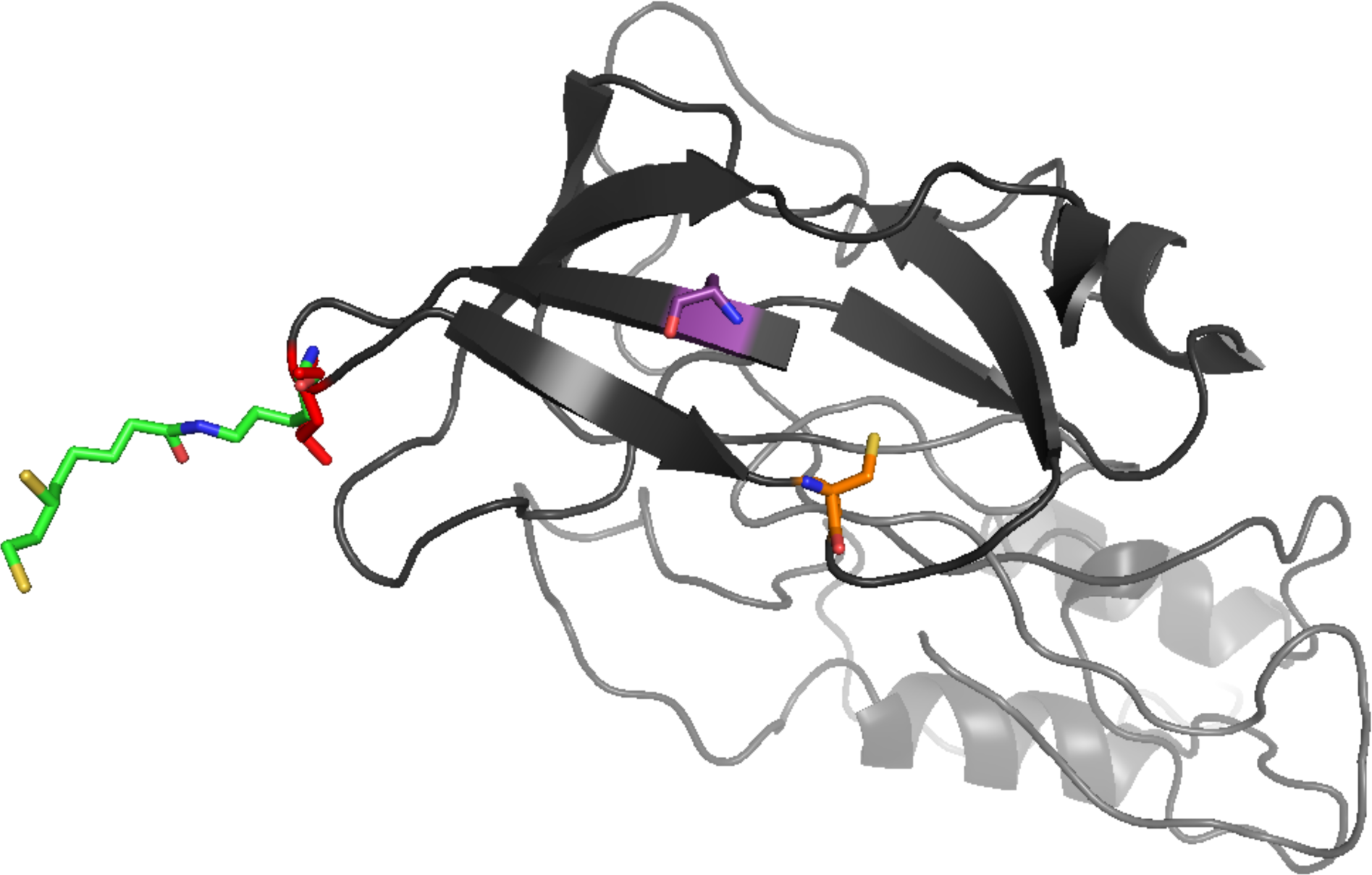
Three-dimensional homology model of *C. elegans* DBT-1. A three-dimensional homology model of *C. elegans* DBT-1 (black) aligned to human pyruvate dehydrogenase lipoyl domain (PDB:1Y8N) is shown. The C78 residue that confers resistance to arsenic trioxide is highlighted in orange, and the C65 residue is highlighted in purple. The *C. elegans* K71 residue is highlighted in red, and the human lipoylated lysine is highlighted in green.

**Figure 2-figure supplement 5:**
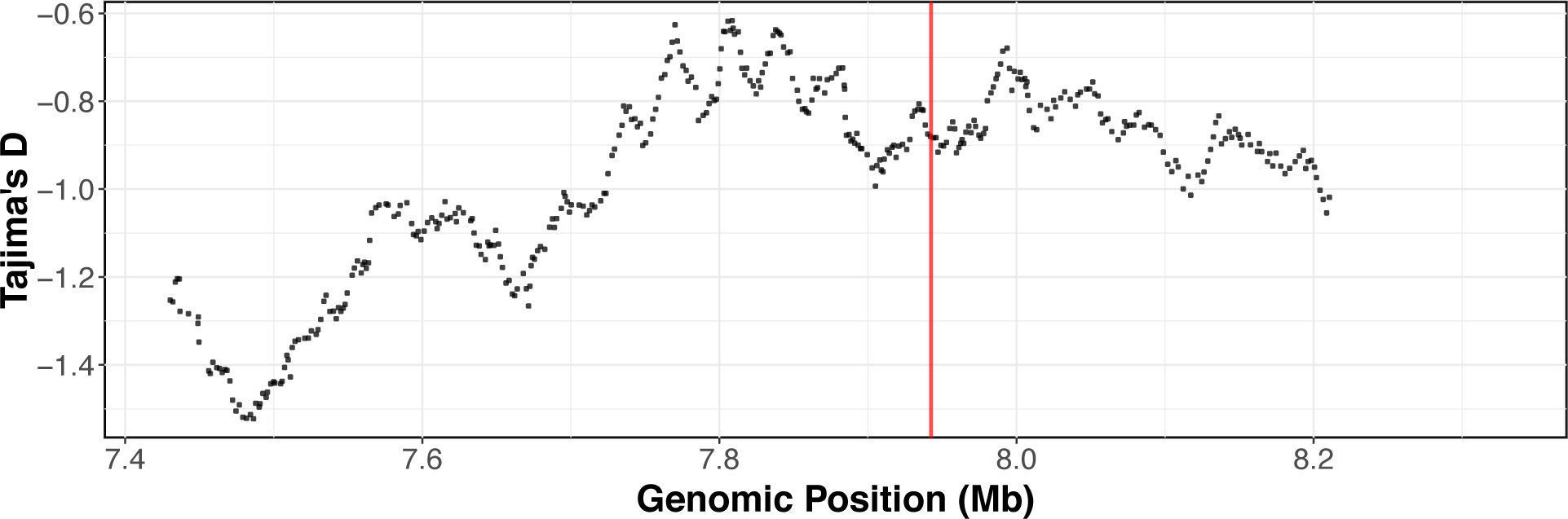
Tajima’s *D* across the arsenic trioxide QTL confidence interval. Divergence, as measured by Tajima’s D, is shown across the arsenic trioxide QTL confidence interval (II:7,430,000-8,330,000). The whole-genome SNV data set [24,49] was used for Tajima’s D calculations. Window size for the calculations was 500 SNVs with a 10 SNV sliding window size. The vertical red line marks the position of the *dbt-1* locus.

**Figure 2-figure supplement 6:**
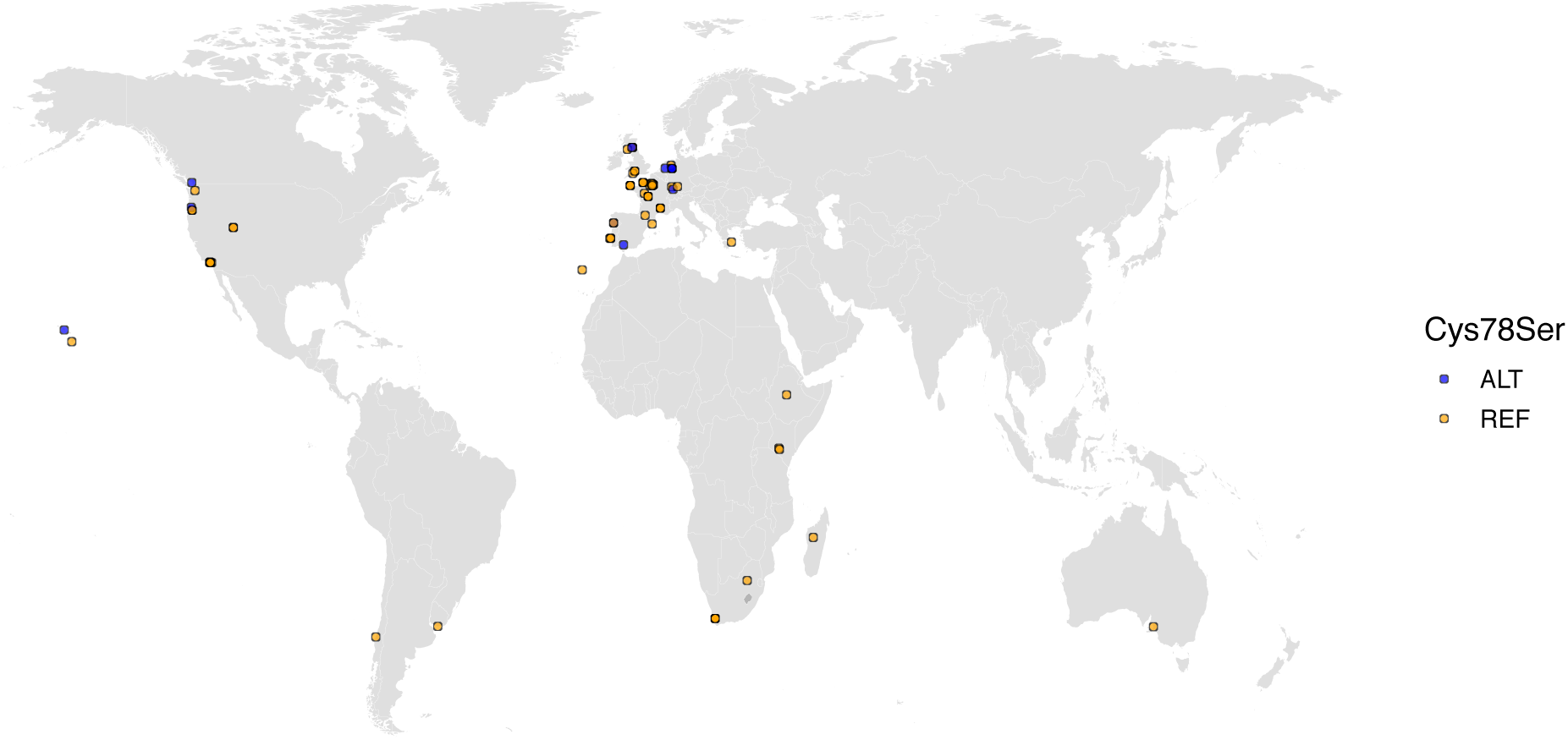
The worldwide distribution of the DBT-1(C78S) allele. Cysteine (REF) is shown in orange and serine (ALT) is shown in blue. Latitude and longitude coordinates of sampling locations were used to plot individual strains on the map.

## Supplementary Files

**Figure 1-source data 1:** Arsenic dose response trait data for four strains in 0, 250, 500, 1000, or 2000µM arsenic, labelled as water, arsenic1, arsenic2, arsenic3, arsenic4 in the Condition column. (Used to generate Supplementary Figure 1A-B)

**Figure 1-source data 2:** Broad-sense heritability estimates from arsenic dose response data. Estimates were calculated using the lmer function of the lme4 R package by fitting the mixed effect model equation lmer(value ∼ (1|strain)), where value is the trait value and strain is the strain name. (Used to generate Supplementary Figure 3)

**Figure 1-source data 3:** Arsenic dose response estimates of effect sizes estimated by fitting the linear model: trait value ∼ strain. (Used to generate Supplementary Figure 3)

**Figure 1-source data 4:** Arsenic dose response loadings of principal components (PCs) for the PCs that explain up to 90% of the total variance in the trait data. Condition names correspond to those in Supplementary File 1. (Used to generate Supplementary Figure 2A-D)

**Figure 1-source data 5:** Arsenic dose response principal component (PC) eigenvectors for the PCs that explain up to 90% of the total variance in the data set. PCA was performed all arsenic concentrations together in order to look at the relative PC value for each concentration. Condition names correspond to those in Supplementary File 1. (Used to generate Supplementary Figure 1C)

**Figure 1-source data 6:** Arsenic dose response loadings of principal components (PCs) for the PCs that explain up to 90% of the total variance in the trait data. PCA was performed all arsenic concentrations together in order to look at the relative PC value for each concentration.

**Figure 1-source data 7:** Arsenic dose response trait correlations where each row corresponds to the Pearson correlation coefficient for two traits. Condition names correspond to those in Supplementary File 1. (Used to generate Supplementary Figure 1A-D)

**Figure 1-source data 8:** RIAIL phenotype data used for linkage mapping. (Used to generate Figure 1B and Supplementary Figure 5)

**Figure 1-source data 9:** RIAIL trait correlations where each row corresponds to the Pearson correlation coefficient for two traits. (Used to generate Supplementary Figure 4A)

**Figure 1-source data 10:** RIAIL loadings of principal components (PCs) for the PCs that explain up to 90% of the total variance in the trait data. (Used to generate Supplementary Figure 4B)

**Figure 1-source data 11:** Results from linkage mapping experiment. (Used to generate Figure 1A, Supplementary Figure 6, and Supplementary Figure 8)

**Figure 1-source data 12:** Genomic heritability estimates from RIAIL phenotype data. (Used to generate Supplementary Figure 7)

**Figure 1-source data 13:** Phenotypes of NILs and CRISPR allele swap strains in the presence of 1000 µM arsenic trioxide after correcting for strain differences in control conditions. (Used to generate Figure 1C, Figure 3, Supplementary Figure 9, and Supplementary Figure 16)

**Figure 1-source data 14:** NIL and CRISPR allele swap trait correlations. (Used to generate Supplementary Figure 10A)

**Figure 1-source data 15:** NIL and CRISPR allele swap trait loadings of principal components (PCs) for the PCs that explain up to 90% of the total variance in the trait data. (Used to generate Supplementary Figure 10B)

**Figure 1-source data 16:** NIL genotypes generated from whole-genome sequencing.

**Figure 2-source data 1:** All wild-isolate traits used for genome-wide association mapping.

**Figure 2-source data 2:** Wild isolate trait loadings of principal components (PCs) for the PCs that explain up to 90% of the total variance in the trait data. (Used to generate Supplementary Figure 12B)

**Figure 2-source data 3:** Wild isolate trait correlations. (Used to generate Supplementary Figure 12A).

**Figure 2-source data 4:** GWA mapping results for PC1. (Used to generate Figure 1A-B)

**Figure 2-source data 5:** Genotype matrix used for genome-wide mapping.

**Figure 2-source data 6:** Genomic heritability estimates of wild isolate traits. (Used to generate Supplementary Figure 13)

**Figure 2-source data 7:** All QTL identified by GWA mapping (Used to generate Supplementary Figure 14).

**Figure 2-source data 8:** Fine-mapping results for PC1. (Used to generate Figure 1C and Supplementary Figure 15)

**Figure 2-source data 9:** Tajima’s D of GWA mapping confidence interval. (Used to generate Supplementary Figure 26)

**Figure 2-source data 10:** Isolation locations of strains used in GWA mapping. (Used to generate Supplementary Figure 27)

**Figure 4-source data 1:** Metabolite measurements for the CB4856 and CB4856 allele swap strains (Used for Figure 4B and Supplementary Figures 18-20)

**Figure 4-source data 2:** Processed metabolite measurements for the CB4856 and CB4856 allele swap strains (Used for Figure 4B and Supplementary Figures 18-20)

**Figure 4-source data 3:** Metabolite measurements for the N2 and N2 allele swap strains (Used for Figure 4B and Supplementary Figures 18-20)

**Figure 4-source data 4:** Metabolite measurements for N2, CB4856, and both allele-swap strains at the L4 larval stage. (Used to generate Supplementary Figure 21)

**Figure 4-source data 5:** Processed phenotype data for the C15ISO rescue experiment (Used to generate Figure 4C)

**Figure 4-source data 6:** Trait correlations for the for the C15ISO rescue experiment (Used to generate Supplementary Figure 22A)

**Figure 4-source data 7:** C15ISO rescue trait loadings of principal components (PCs) for the PCs that explain up to 90% of the total variance in the trait data. (Used to generate Supplementary Figure 2B)

**Figure 5-source data 1:** Human cell line read data for CRISPR swap experiment in 293 T cells.(Used to generate Figure 5B)

**Figure 5-source data 2:** Results from Fisher’s exact test of human cell line read data for CRISPR swap experiment in 293 T cells.

**Figure 5-source data 3:** Metabolite measurements from human cell line experiments.

**Figure 2-source data 9:** Tajima’s D of GWA mapping confidence interval. (Used to generate Supplementary Figure 26)

**Figure 2-source data 10:** Isolation locations of strains used in GWA mapping. (Used to generate Supplementary Figure 27)

**Supplementary File 1:** Plasmid used for editing human cells with the S112C and R113C edits.

**Supplementary File 2:** Plasmid used for editing human cells with the W84C edit.

## Format of Supplementary Files

**Figure 1-source data 1:**
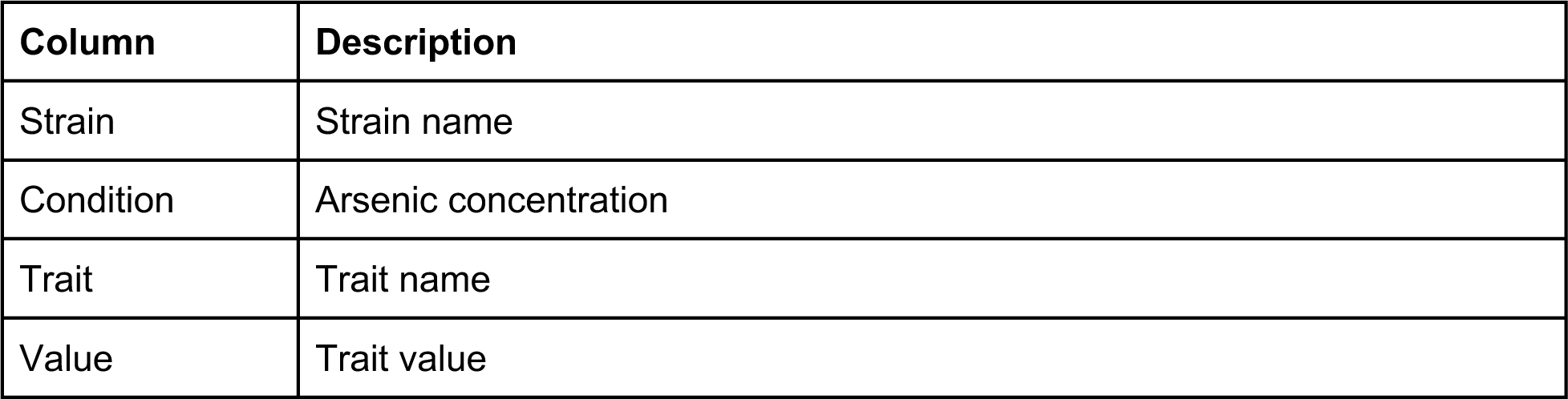
Arsenic dose response trait data for four strains in 0, 250, 500, 1000, or 2000µM arsenic, labelled as water, arsenic1, arsenic2, arsenic3, arsenic4 in the Condition column.

**Figure 1-source data 2:**
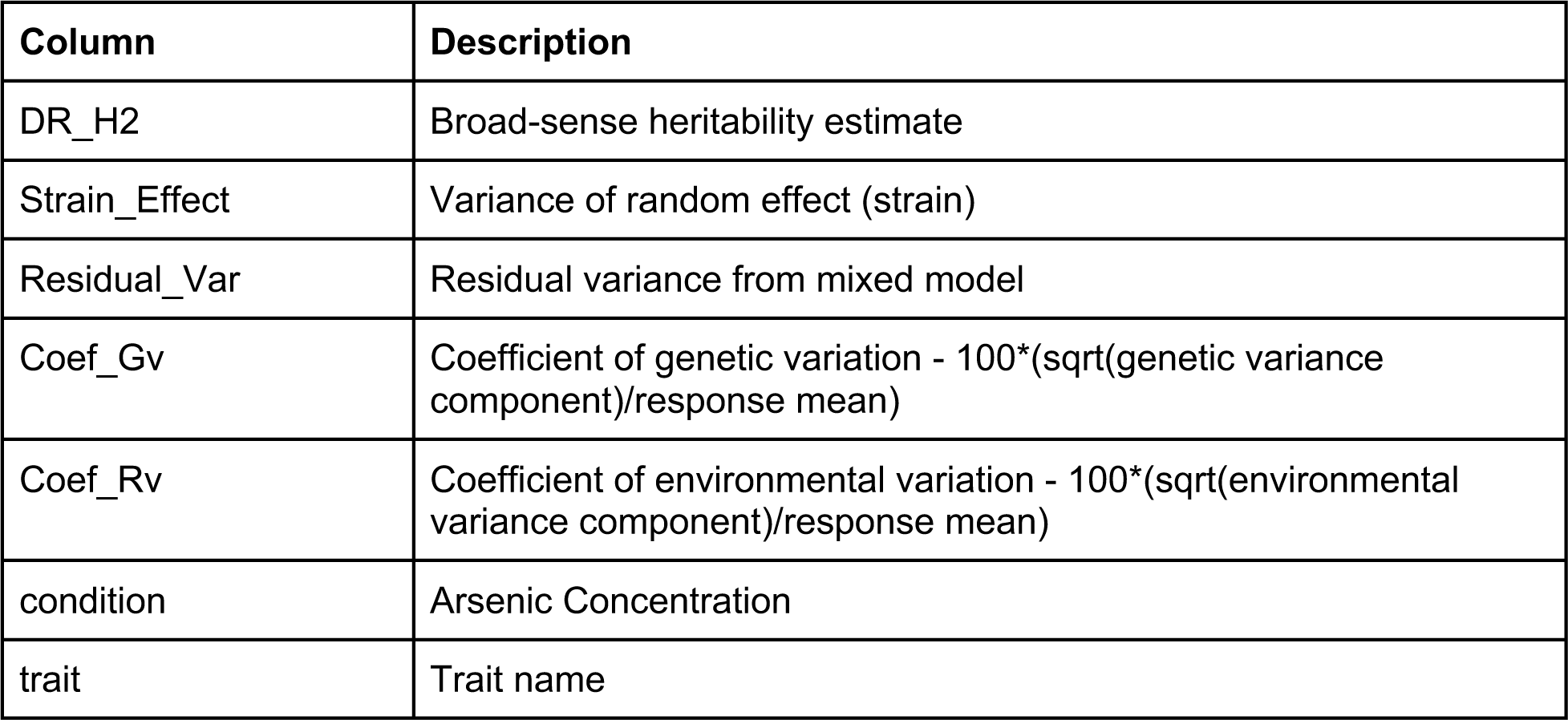
Broad-sense heritability estimates from arsenic dose response data. Estimates were calculated using the lmer function of the lme4 R package by fitting the mixed effect model equation lmer(value ∼ (1|strain)), where value is the trait value and strain is the strain name.

**Figure 1-source data 3:**
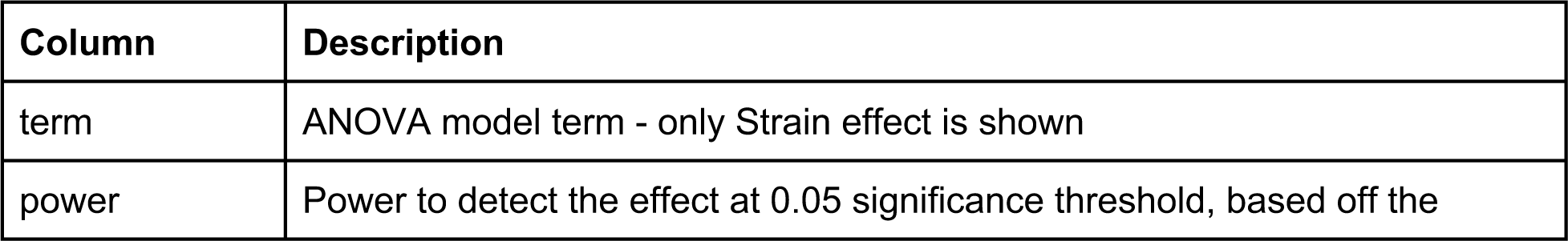
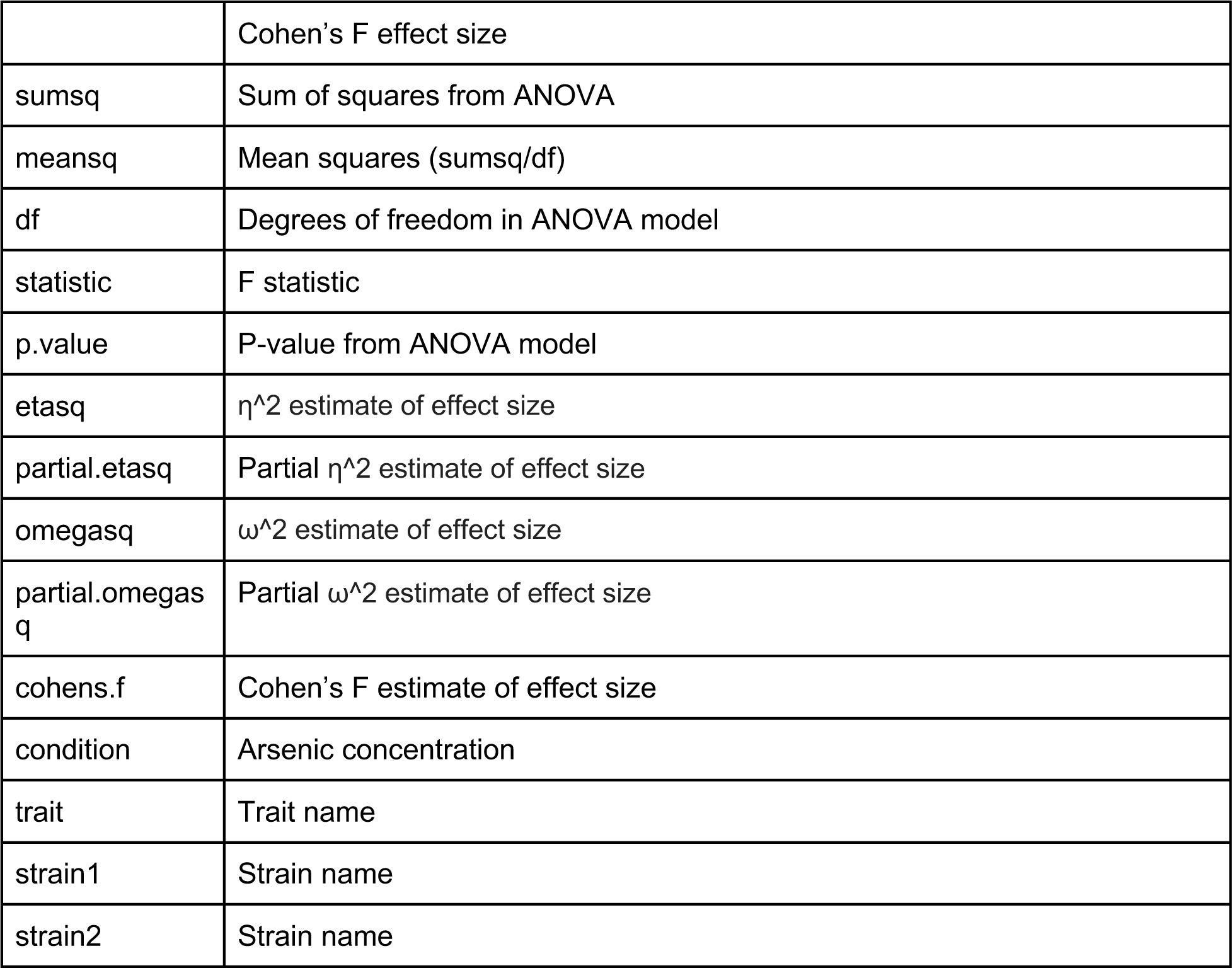
Arsenic dose response estimates of effect sizes estimated by fitting the linear model: trait value ∼ strain. Estimates were performed for pairs of strains and for all strains. When all strains were used for the estimates, the strain1 and strain2 column values are “ALL”. Condition names correspond to those in SuppData1.

**Figure 1-source data 4:**
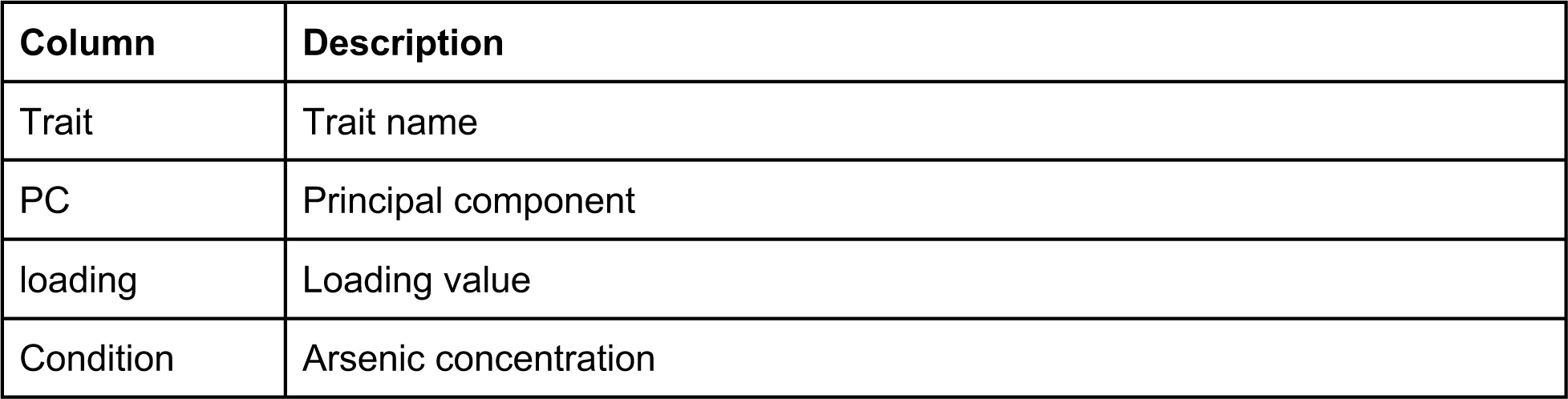
Arsenic dose response loadings of principal components (PCs) for the PCs that explain up to 90% of the total variance in the trait data. Condition names correspond to those in SuppData1

**Figure 1-source data 5:**
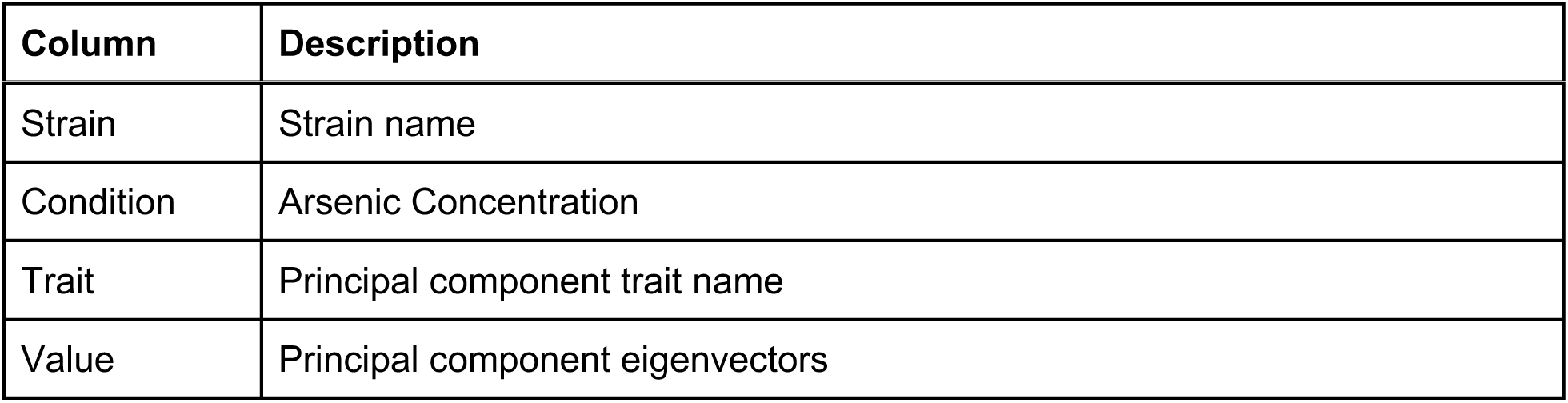
Arsenic dose response principal component (PC) eigenvectors for the PCs that explain up to 90% of the total variance in the data set. Condition names correspond to those in SuppData1.

**Figure 1-source data 6:**
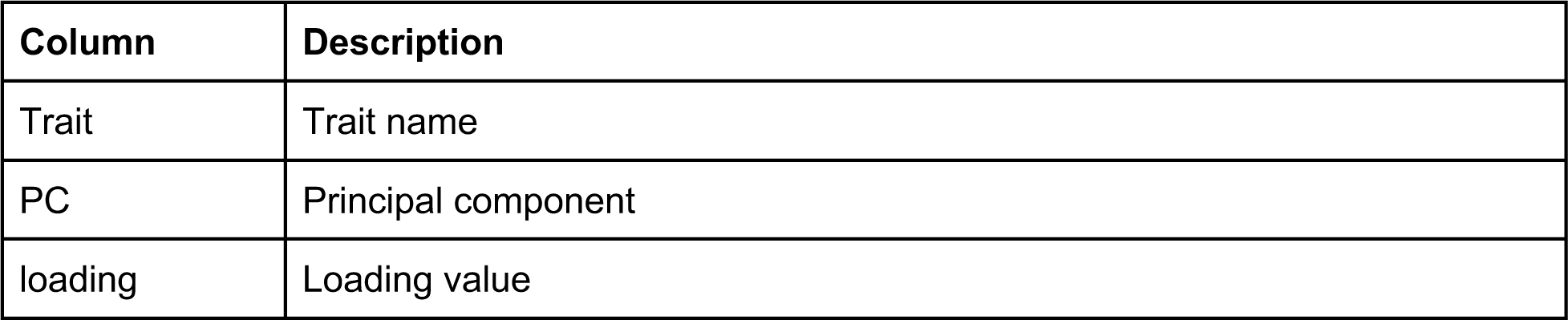
Arsenic dose response loadings of principal components (PCs) for the PCs that explain up to 90% of the total variance in the trait data. PCA was performed all arsenic concentrations together in order to look at the relative PC value for each concentration

**Figure 1-source data 7:**
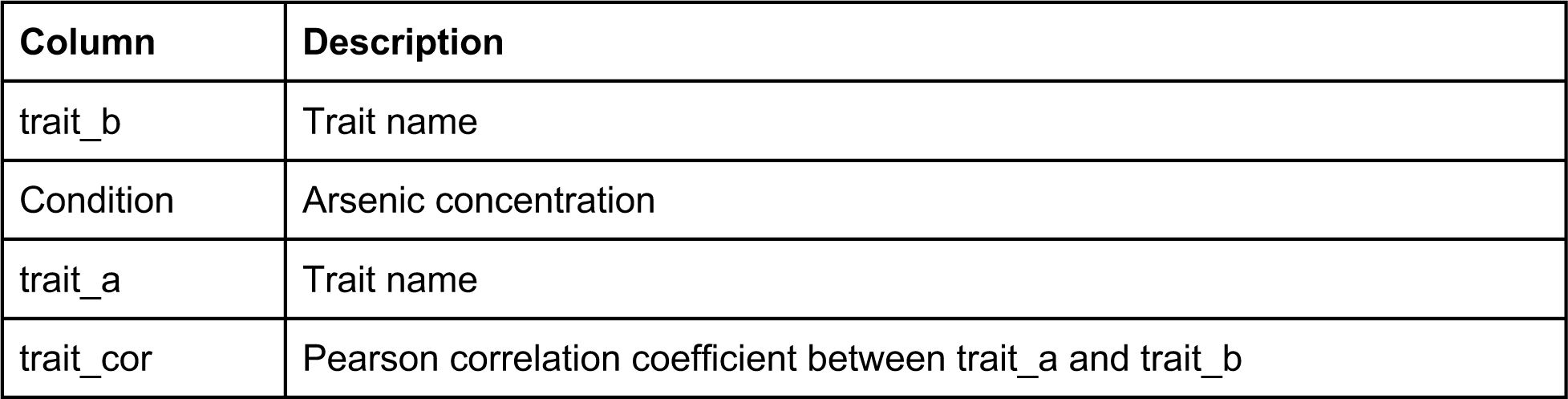
Arsenic dose response trait correlations where each row corresponds to the Pearson correlation coefficient for two traits. Condition names correspond to those in SuppData1.

**Figure 1-source data 8:**
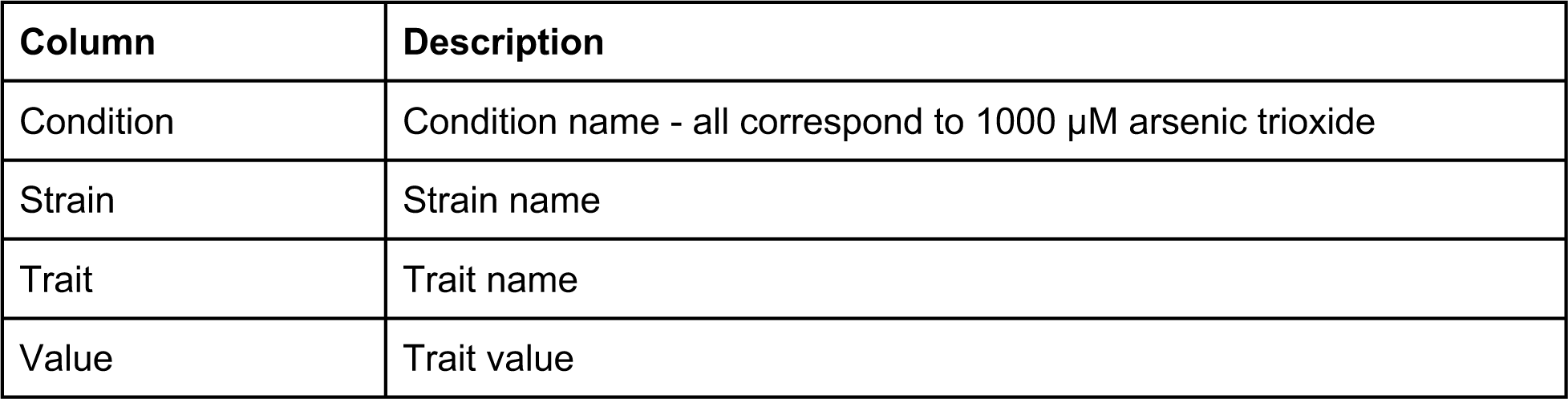
RIAIL phenotype data used for linkage mapping

**Figure 1-source data 9:**
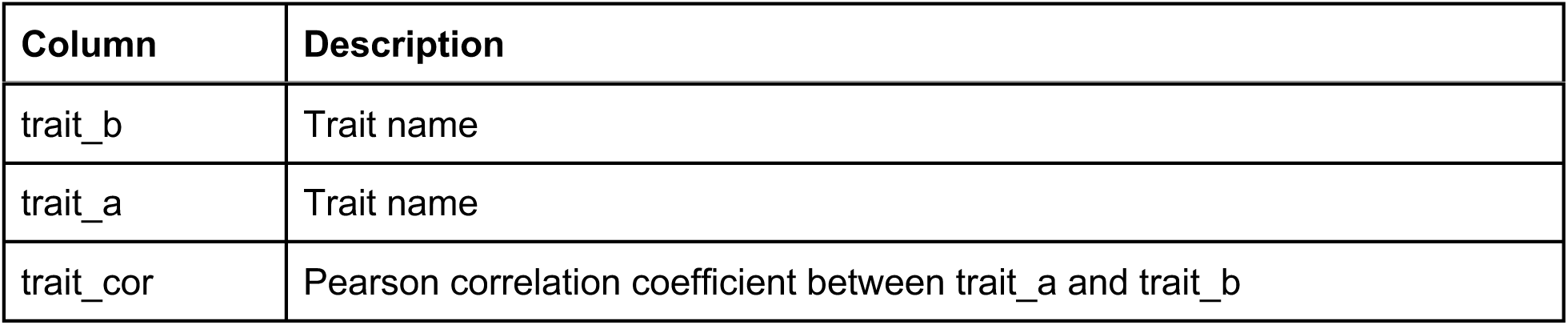
RIAIL trait correlations where each row corresponds to the Pearson correlation coefficient for two traits.

**Figure 1-source data 10:**
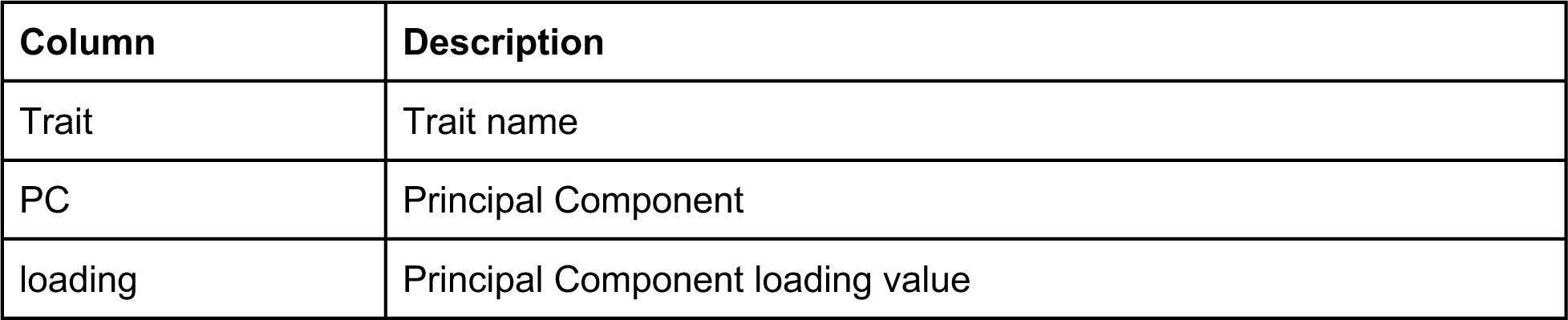
RIAIL loadings of principal components (PCs) for the PCs that explain up to 90% of the total variance in the trait data.

**Figure 1-source data 11:**
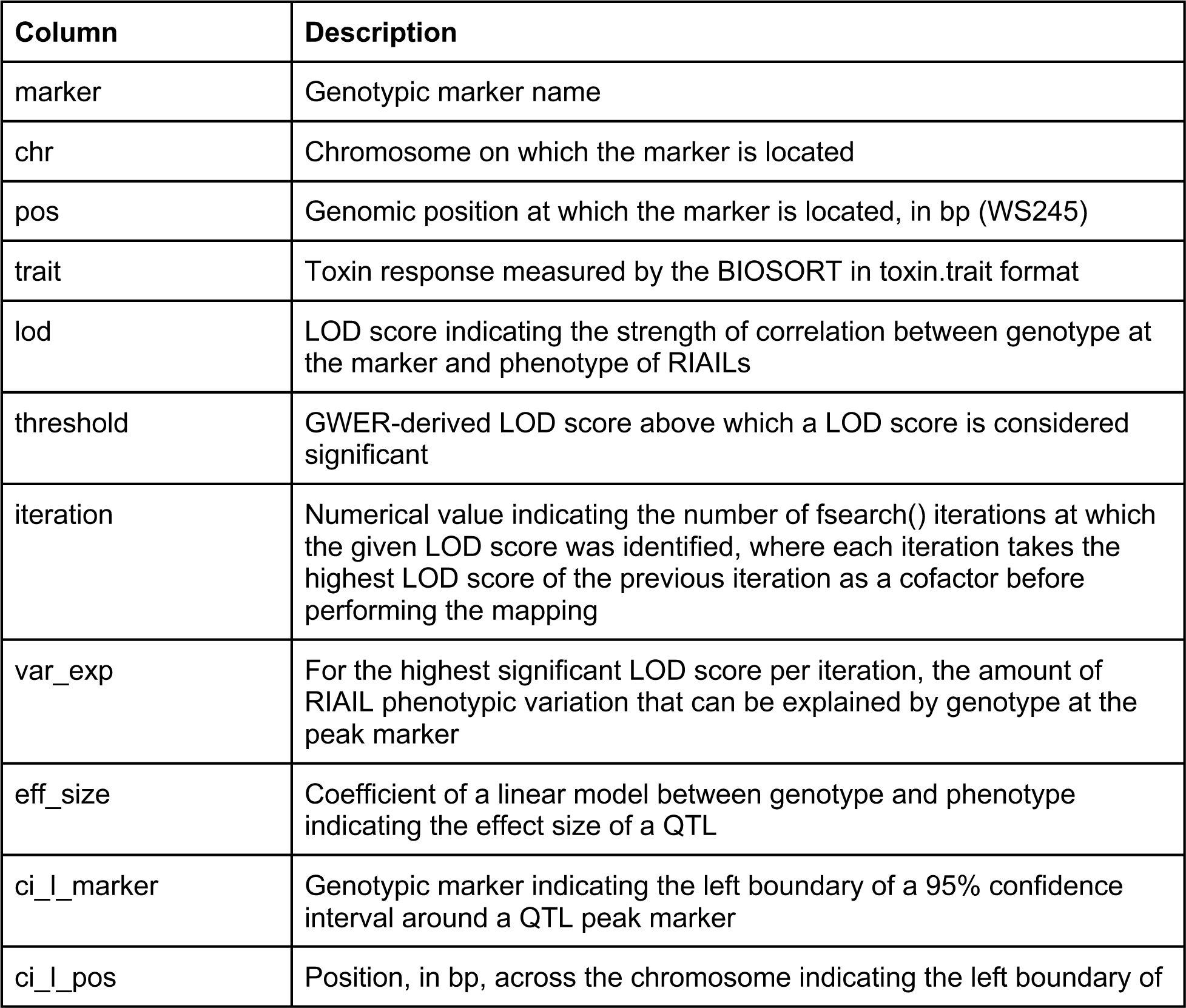
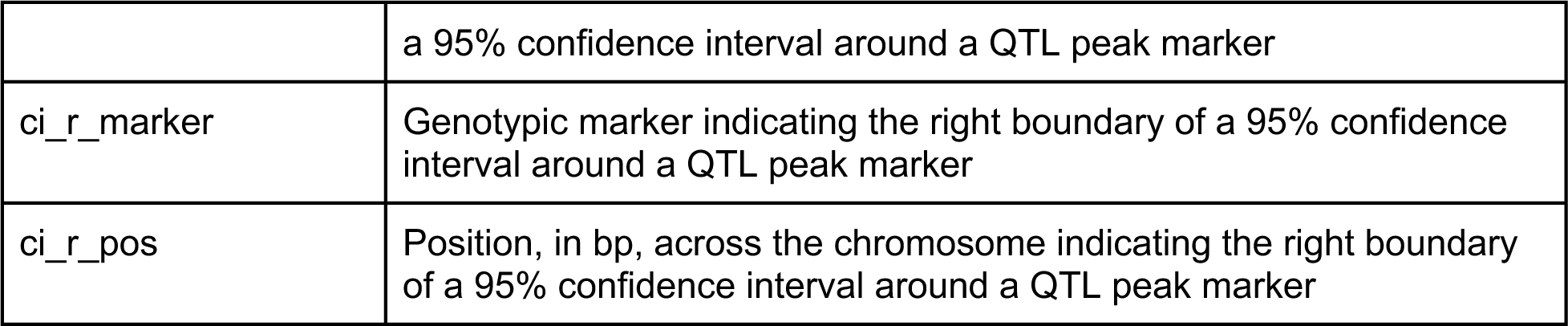
Results from linkage mapping experiment.

**Figure 1-source data 12:**
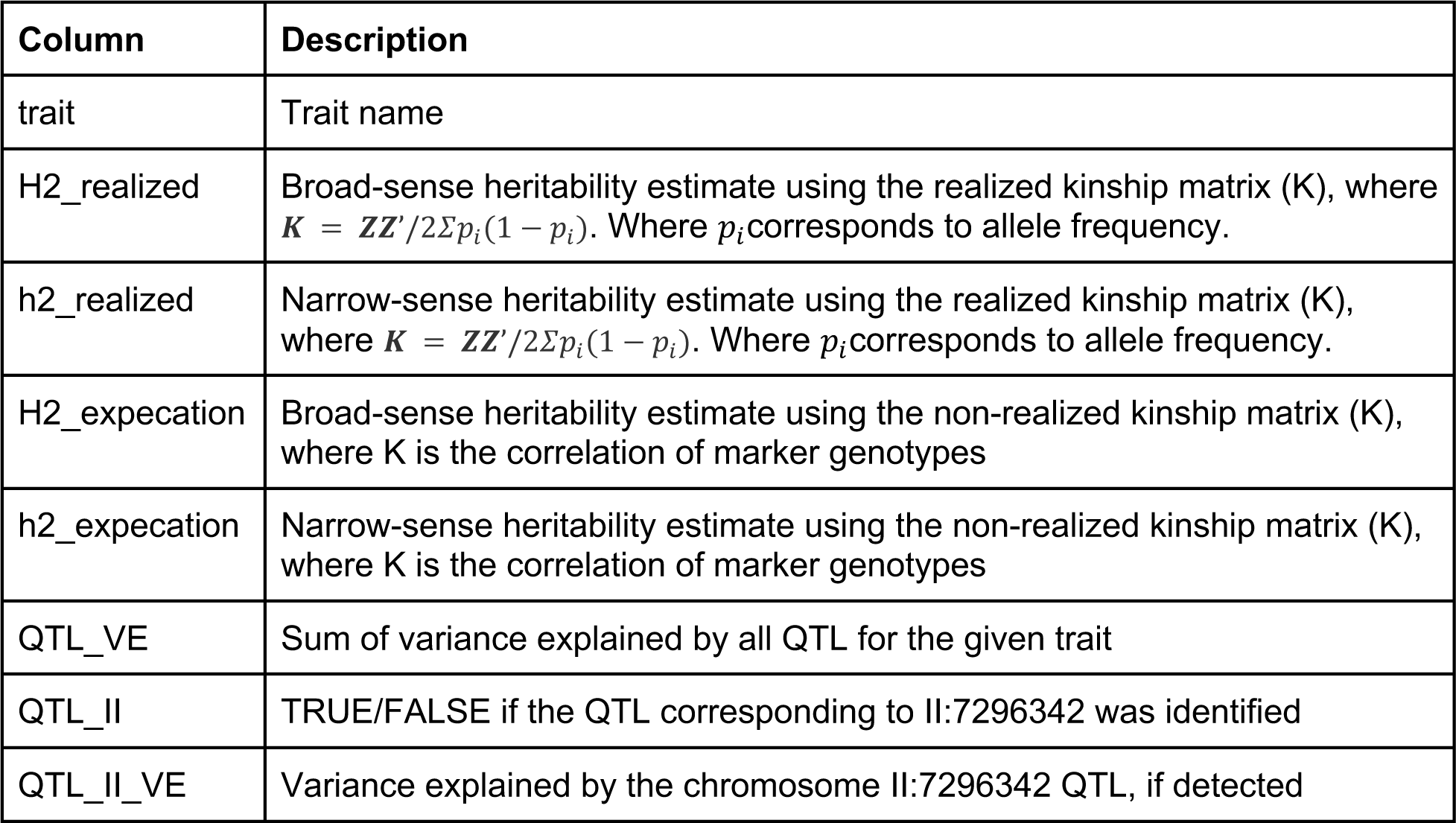
Genomic heritability estimates from RIAIL phenotype data. Two estimates are provided that both utilize a linear mixed effect model with the equation *y* = *Xβ* + *Zu* + *∊* to estimate heritability. The estimates differ in their formulation for the strain additive relatedness matrix, realized relatedness matrices correct for allele frequencies and the expectation matrix does not. For both estimates, the Hadamard product of the additive relatedness matrix was used to calculate the epistatic relatedness matrix.

**Figure 1-source data 13:**
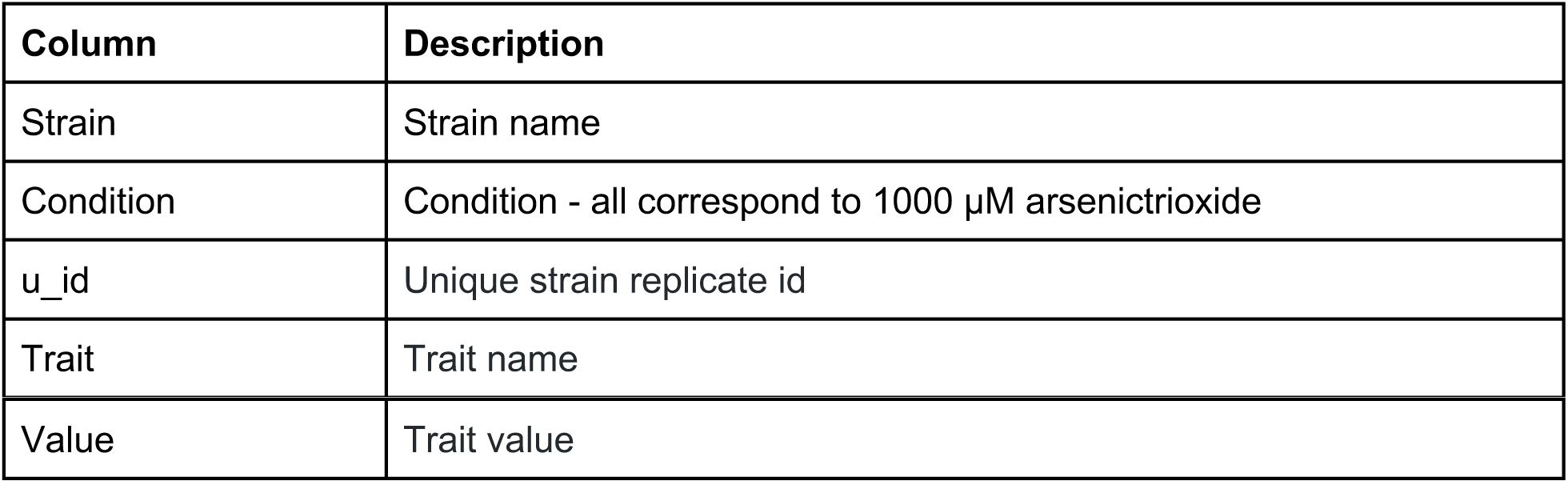
Phenotypes of NILs and CRISPR allele swap strains in the presence of 1000 µM arsenic trioxide after correcting for strain differences in control conditions.

**Figure 1-source data 14:**
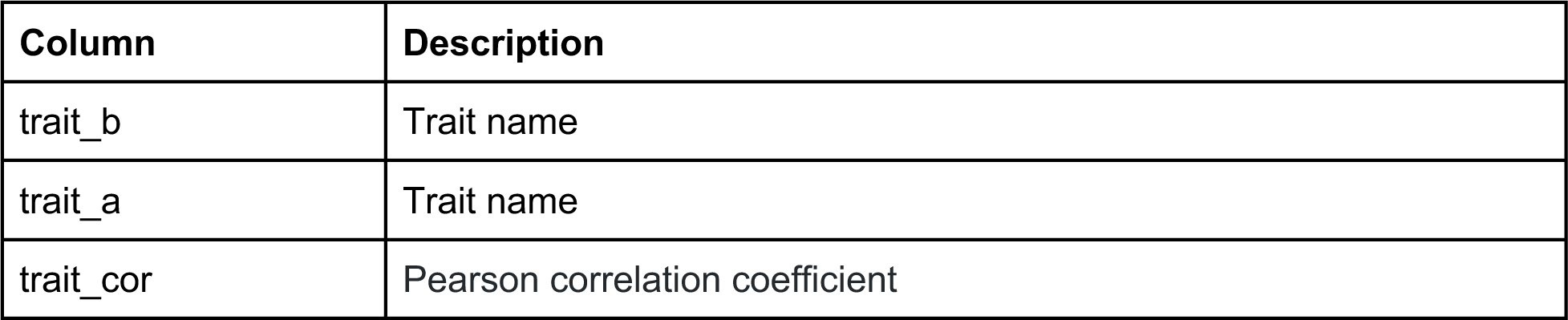
NIL and CRISPR allele swap trait correlations, where each row corresponds to the Pearson correlation coefficient for two traits. All traits correspond to those in 1000 µM arsenic after correcting for strain differences in control conditions

**Figure 1-source data 15:**
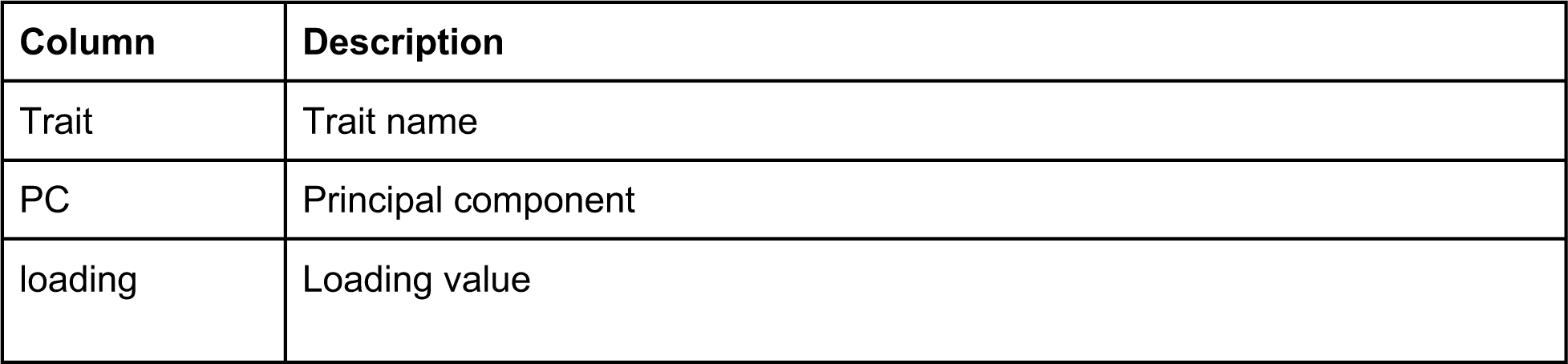
NIL and CRISPR allele swap trait loadings of principal components (PCs) for the PCs that explain up to 90% of the total variance in the trait data. All traits correspond to those in 1000 µM arsenic after correcting for strain differences in control conditions

**Figure 1-source data 16:**
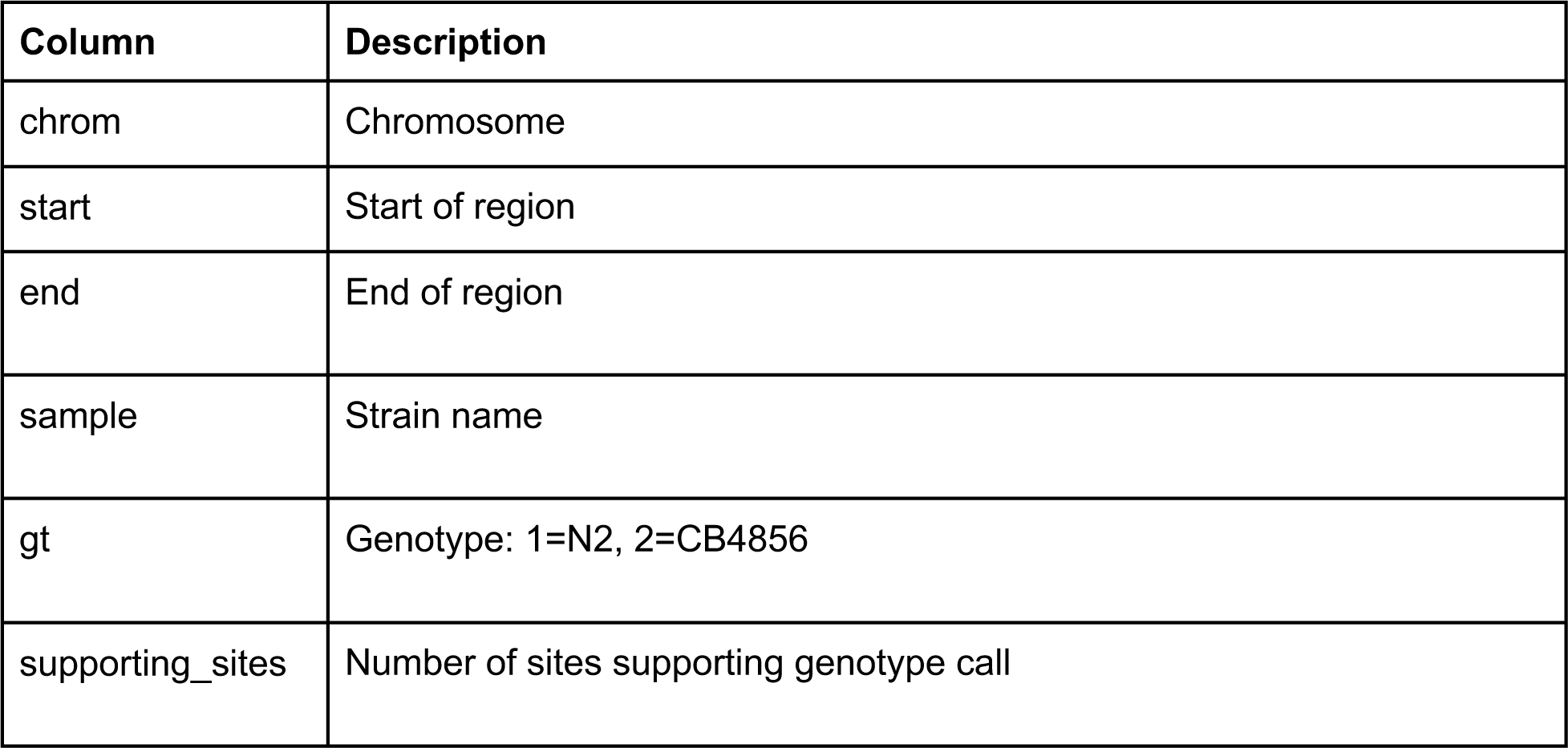
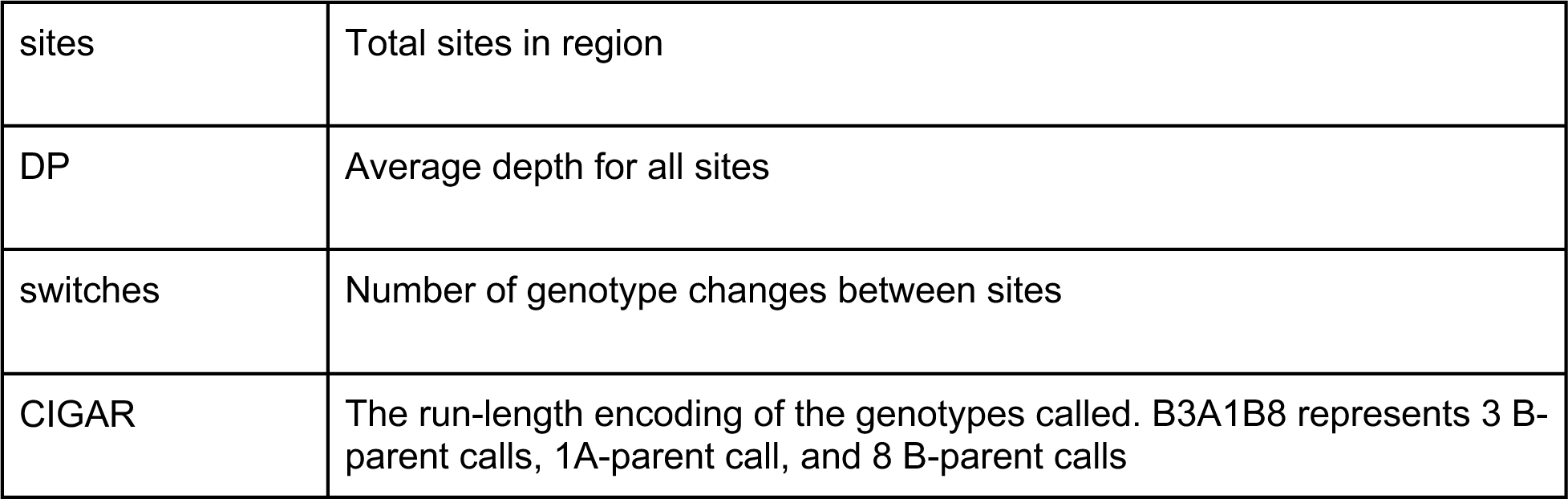
NIL genotypes generated from whole-genome sequencing.

**Figure 2-source data 1:**
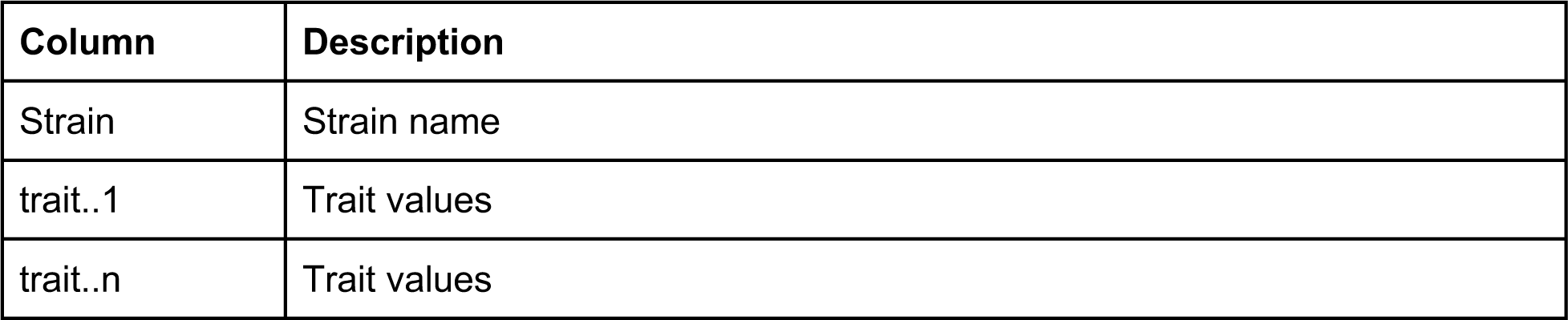
Wild isolate phenotype data in the presence of 1000 µM arsenic trioxide after correcting for growth in control conditions. There is a column for each trait.

**Figure 2-source data 2:**
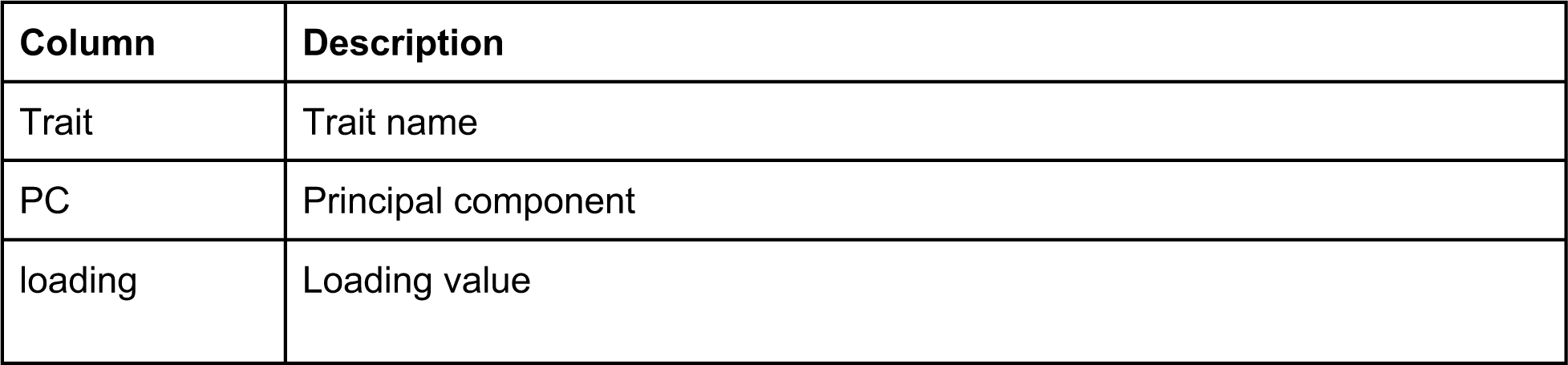
Wild isolates loadings of principal components (PCs) for the PCs that explain up to 90% of the total variance in the trait data. All traits correspond to those in 1000 µM arsenic after correcting for strain differences in control conditions

**Figure 2-source data 3:**
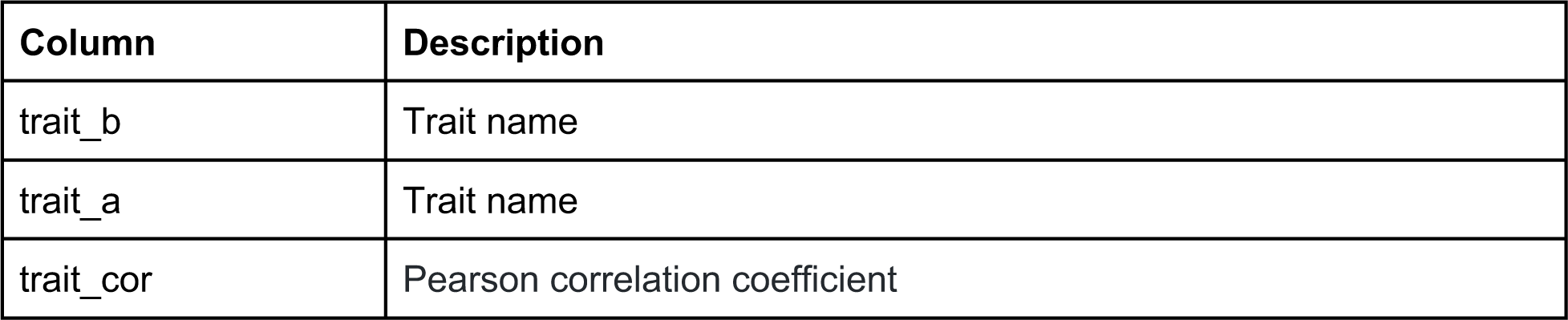
Wild isolate trait correlations where each row corresponds to the Pearson correlation coefficient for two traits. All traits correspond to those in 1000 µM arsenic after correcting for strain differences in control conditions

**Figure 2-source data 4:**
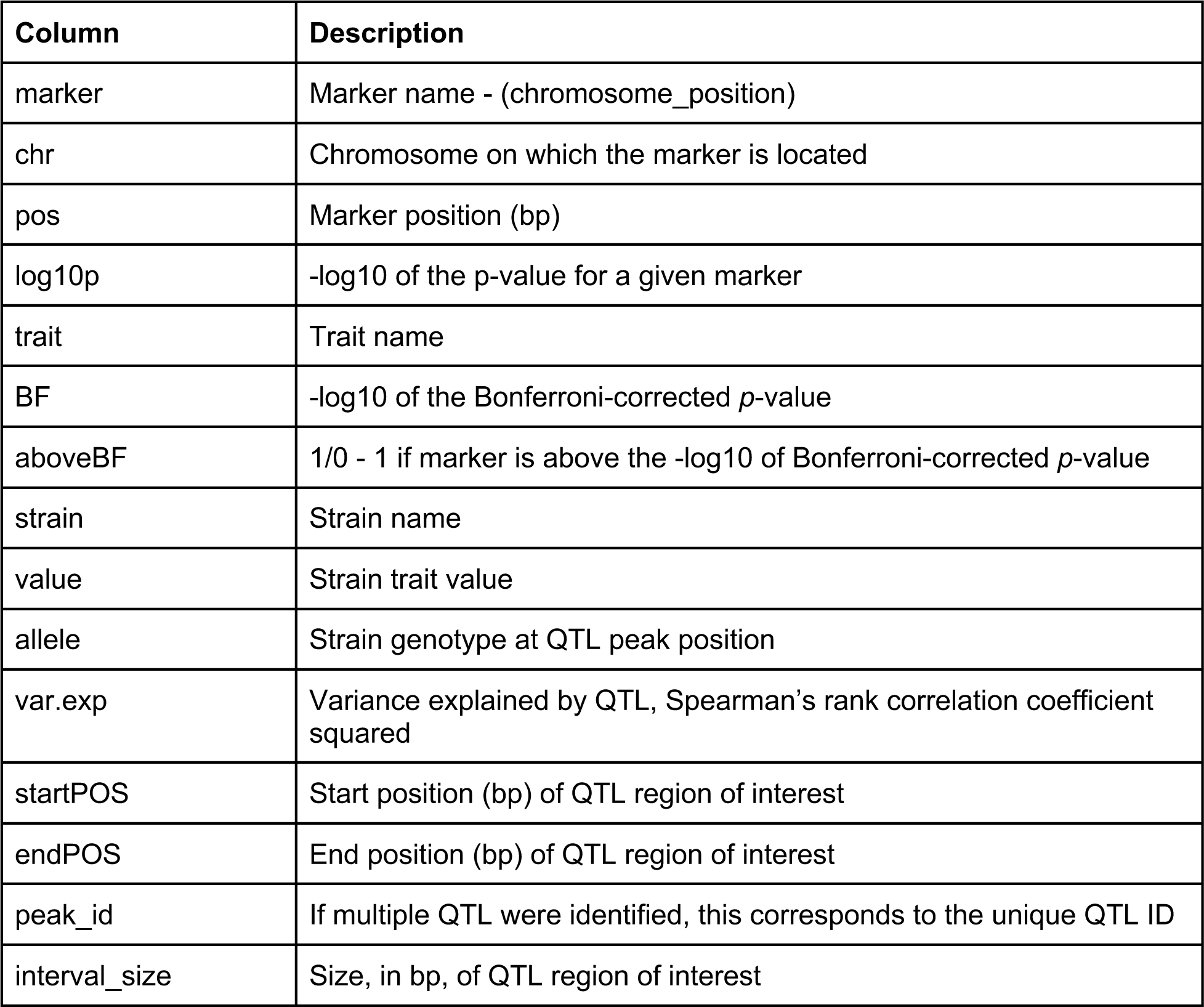
GWA mapping results of PC 1 trait in the presence of 1000 µM arsenic trioxide. Mapping was performed using the EMMA algorithm.

**Figure 2-source data 5:**
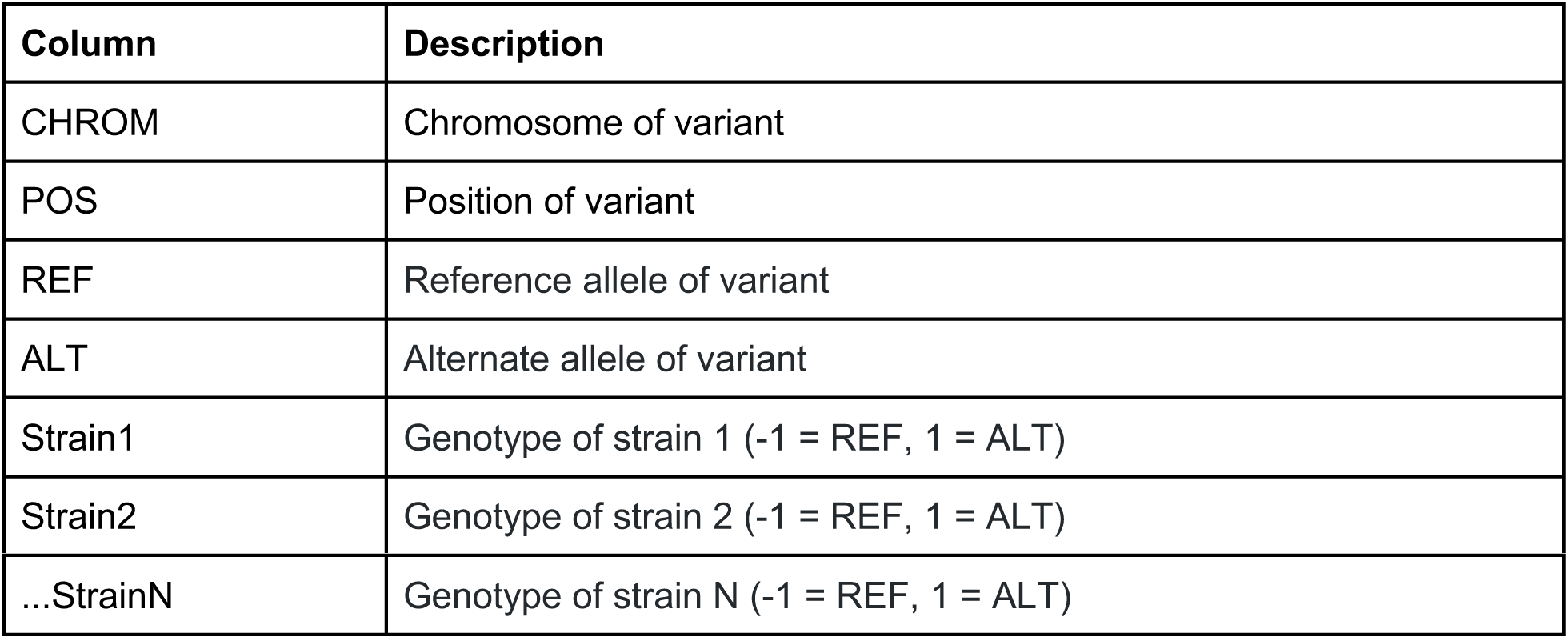
Genotype matrix used for genome-wide mapping. Each row corresponds to a single-nucleotide variant.

**Figure 2-source data 6:**
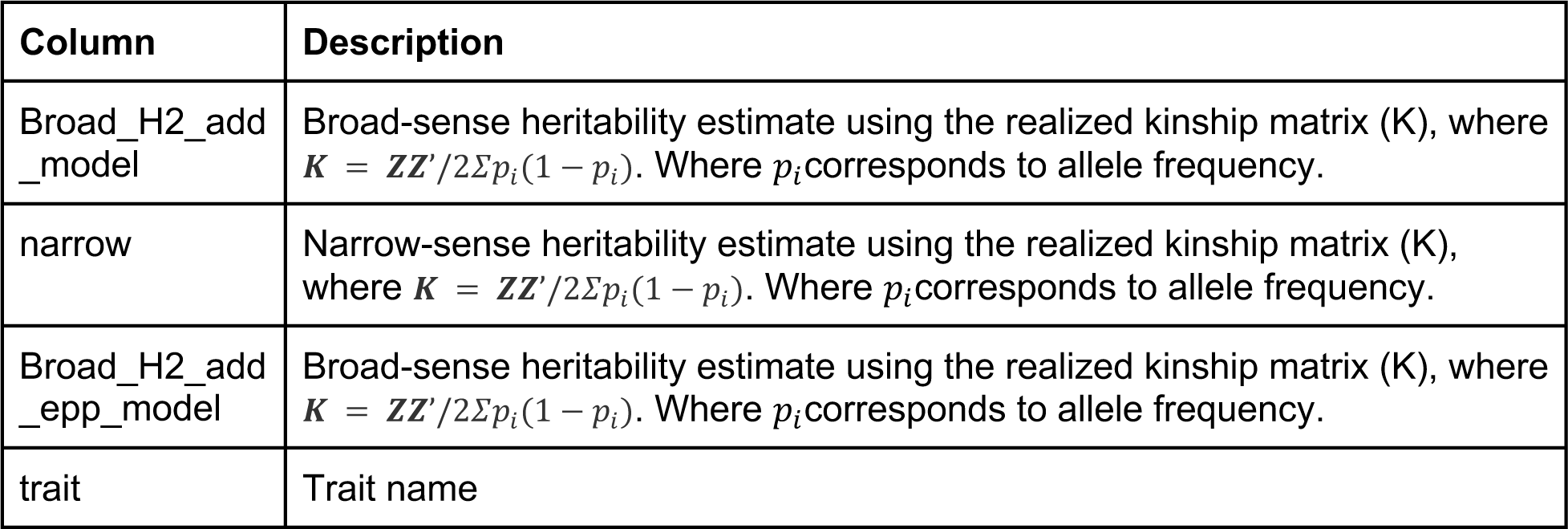
Genomic heritability estimates from wild isolate phenotype data. Estimates were calculated by fitting a linear mixed effect model with the equation *y* = *Xβ* + *Zu* + *∊*. Two models were fitted, one using only additive random effects and one using an additive and epistatic random effects. The Hadamard product of the additive relatedness matrix was used to calculate the epistatic relatedness matrix.

**Figure 2-source data 7:**
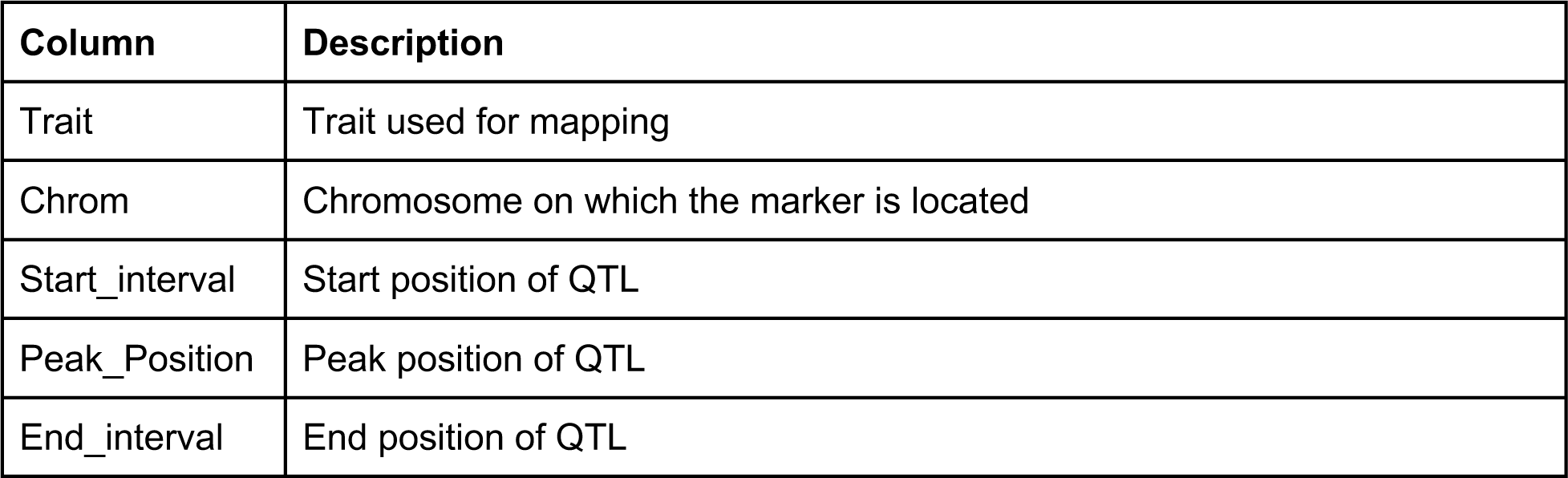
All QTL detected by GWA mapping

**Figure 2-source data 8:**
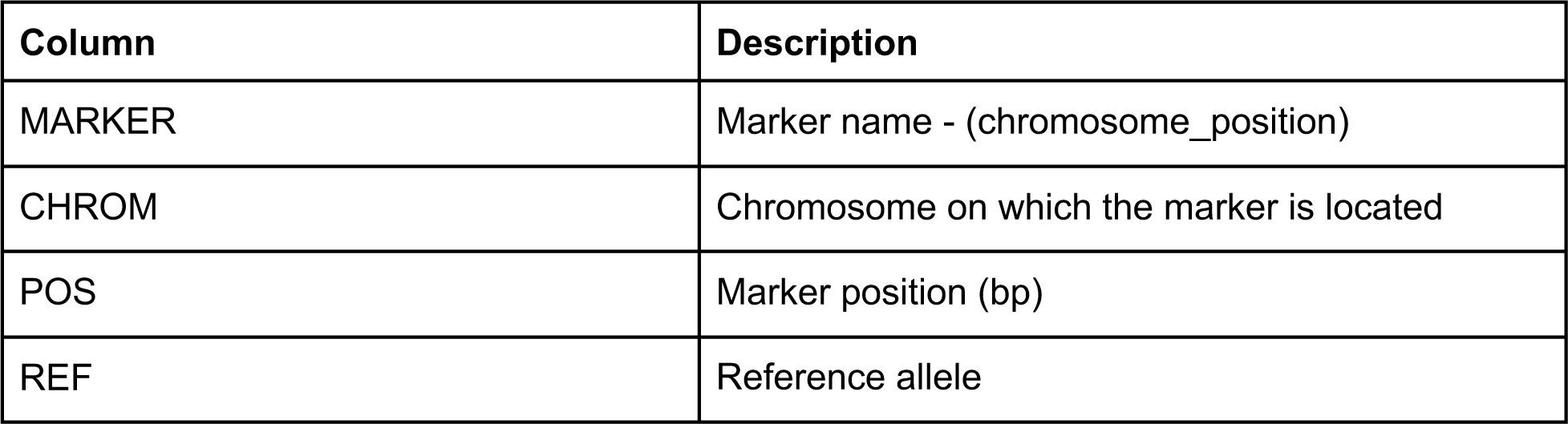
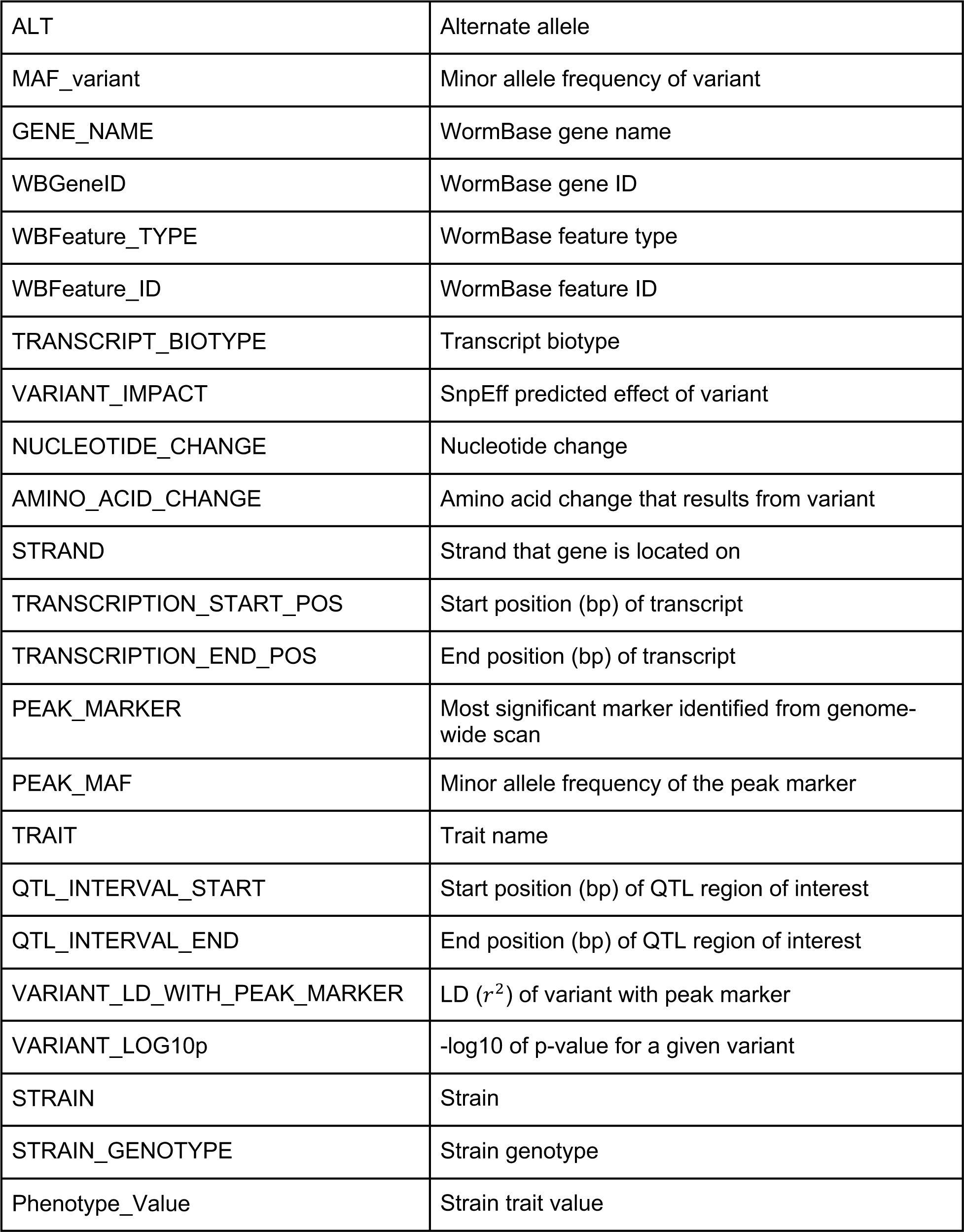
Fine mapping of QTL identified from mapping the first principal component of wild-isolate phenotype data. Each row corresponds to a variant within the QTL region of interest identified from the genome-wide scan.

**Figure 4-source data 1:**
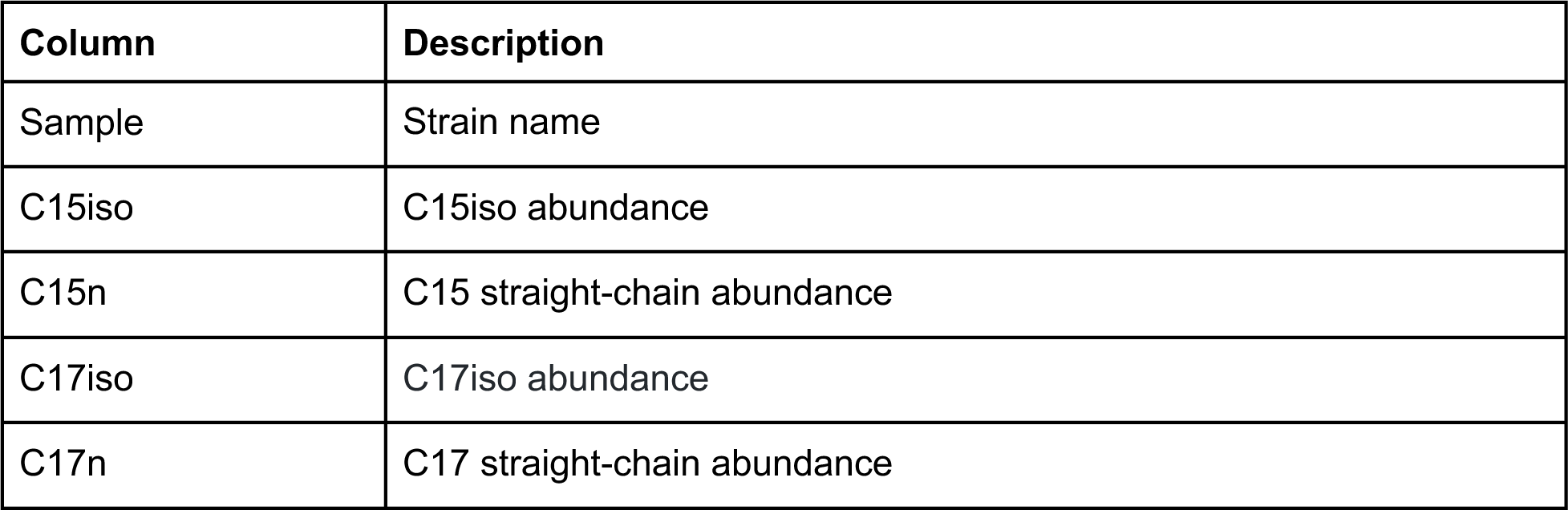
Metabolite measurements of L1 animals exposed to 100 µM arsenic trioxide or control conditions

**Figure 4-source data 2:**
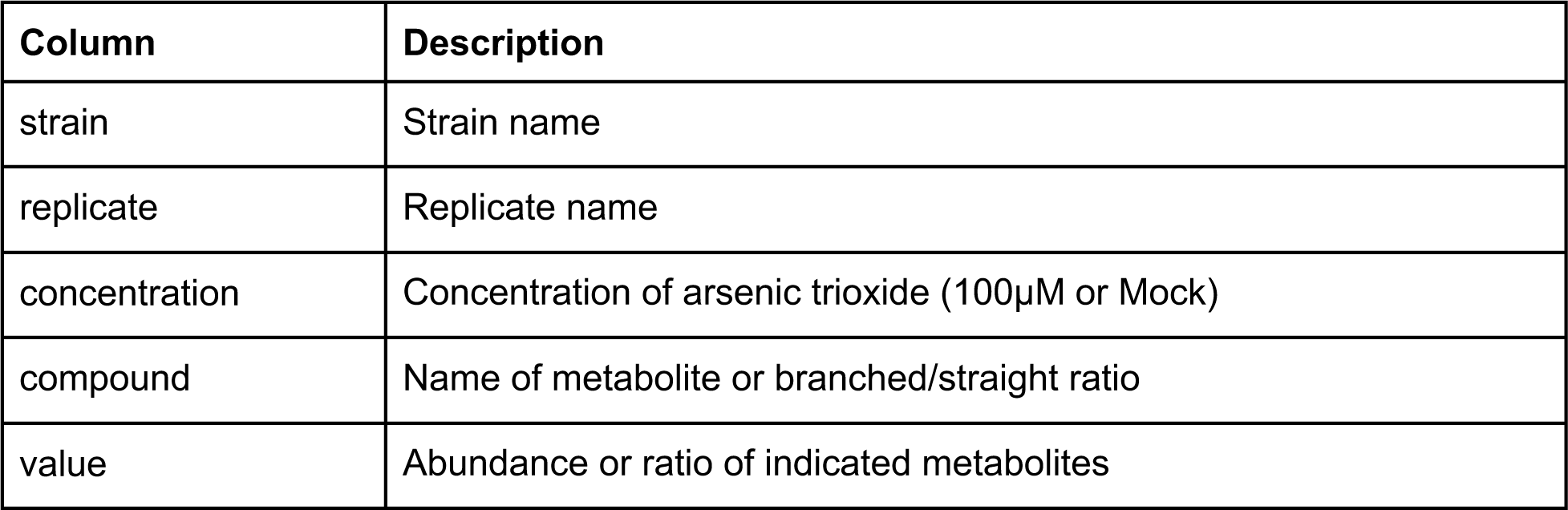
Processed metabolite measurements of L1 animals exposed to 100 µM arsenic trioxide or control conditions. Only data for CB4856 and ECA590 were used from this data set.

**Figure 4-source data 3:**
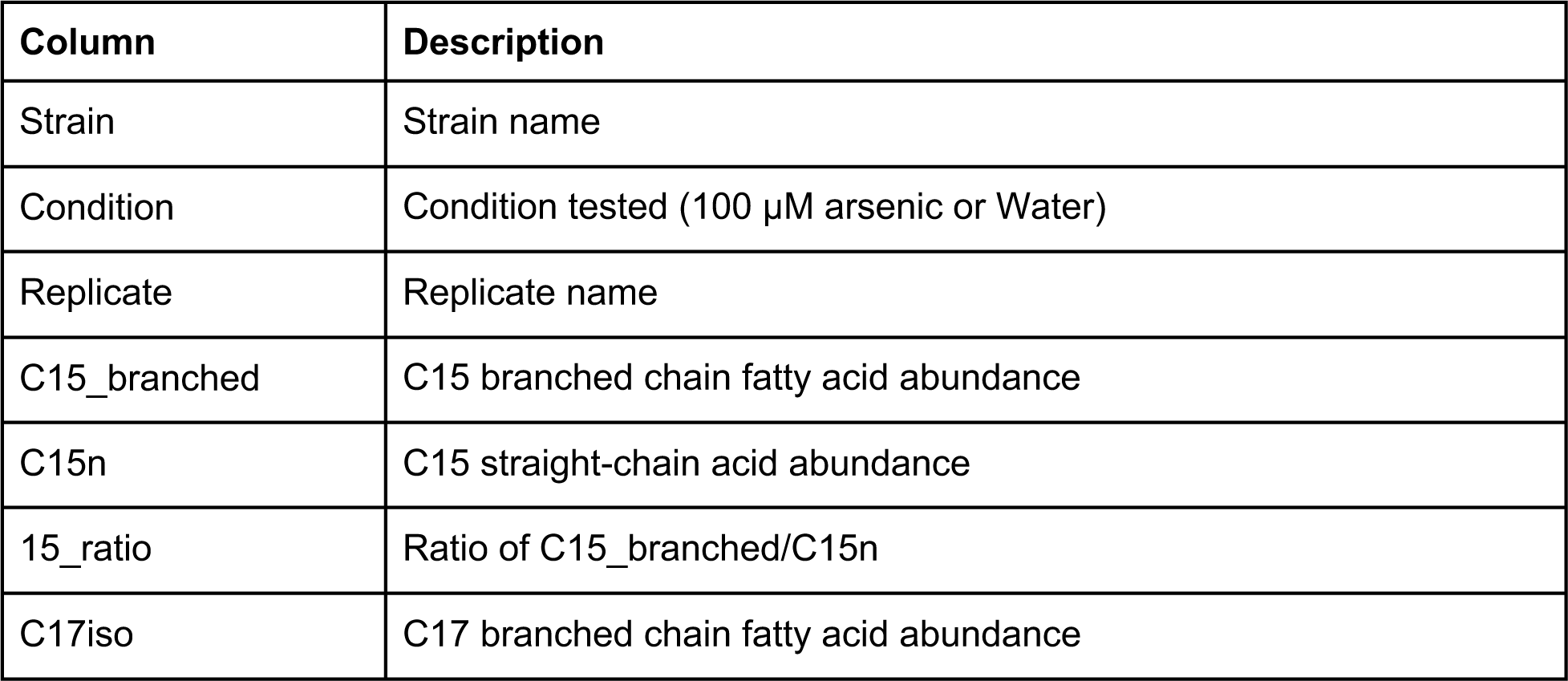
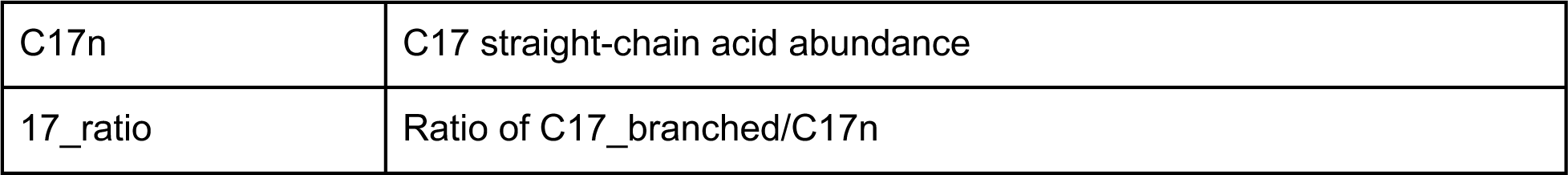
Processed metabolite measurements of N2 and ECA581 L1 animals exposed to 100 µM arsenic trioxide or control conditions. Strain, Condition, Replicate, C15_branched, C15n, 15_ratio, C17iso, C17n, 17_ratio

**Figure 4-source data 4:**
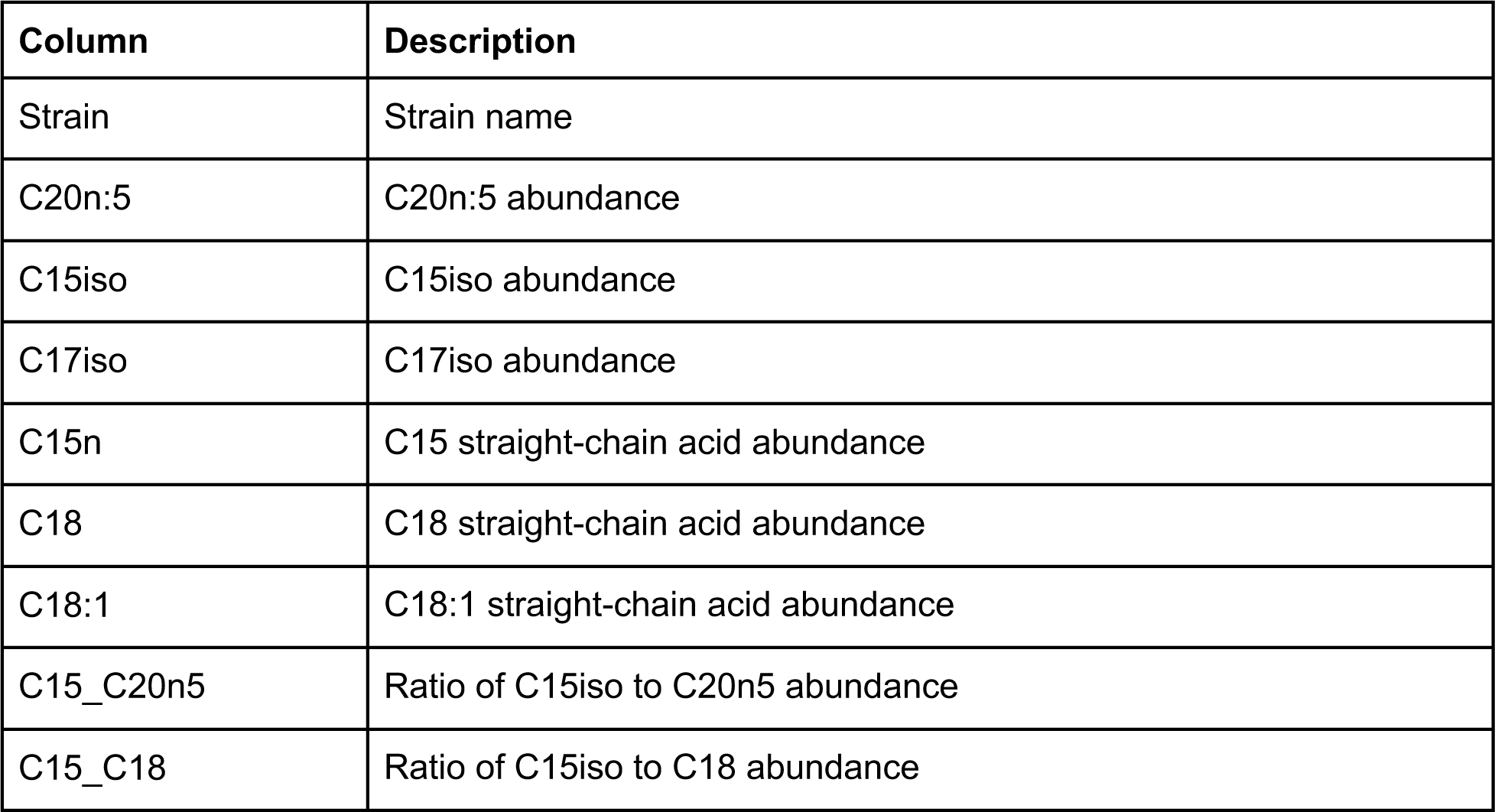
Processed metabolite measurements of CB4856, N2, ECA590, and ECA581 young adult animals in control conditions. Strain, C20n:5, C15iso, C17iso, C18, C18:1, C14, C15_C20n5, C15_C18

**Figure 4-source data 5:**
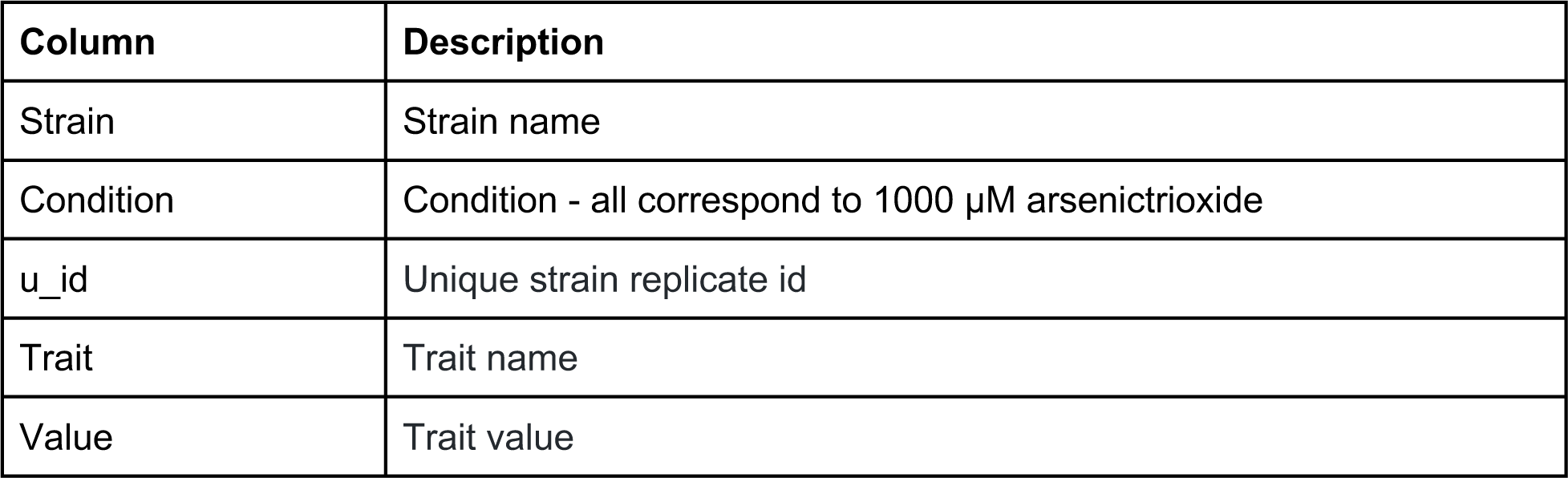
C15ISO rescue experiment phenotypes of parental and CRISPR allele swap strains in the presence of 1000 µM arsenic trioxide after correcting for strain differences in control conditions.

**Figure 4-source data 6:**
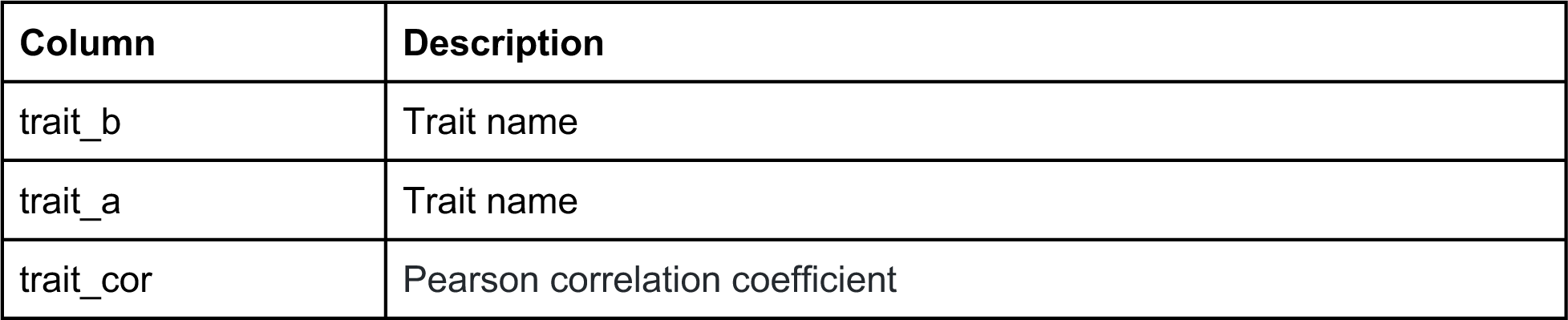
C15ISO rescue experiment trait correlations, where each row corresponds to the Pearson correlation coefficient for two traits. All traits correspond to those in 1000 µM arsenic after correcting for strain differences in control conditions

**Figure 4-source data 7:**
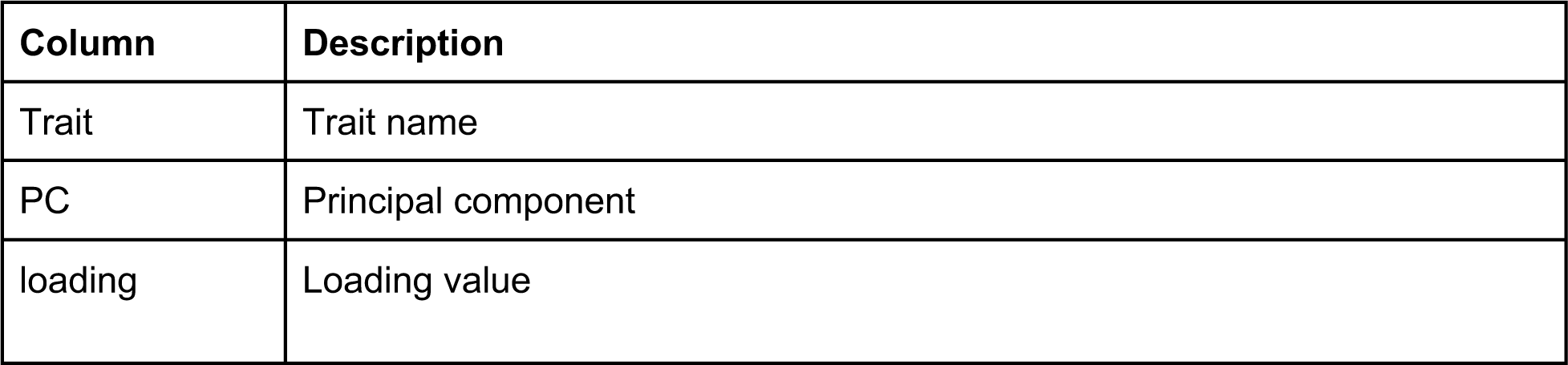
C15ISO rescue experiment trait loadings of principal components (PCs) for the PCs that explain up to 90% of the total variance in the trait data. All traits correspond to those in 1000 µM arsenic after correcting for strain differences in control conditions

**Figure 5-source data 1:**
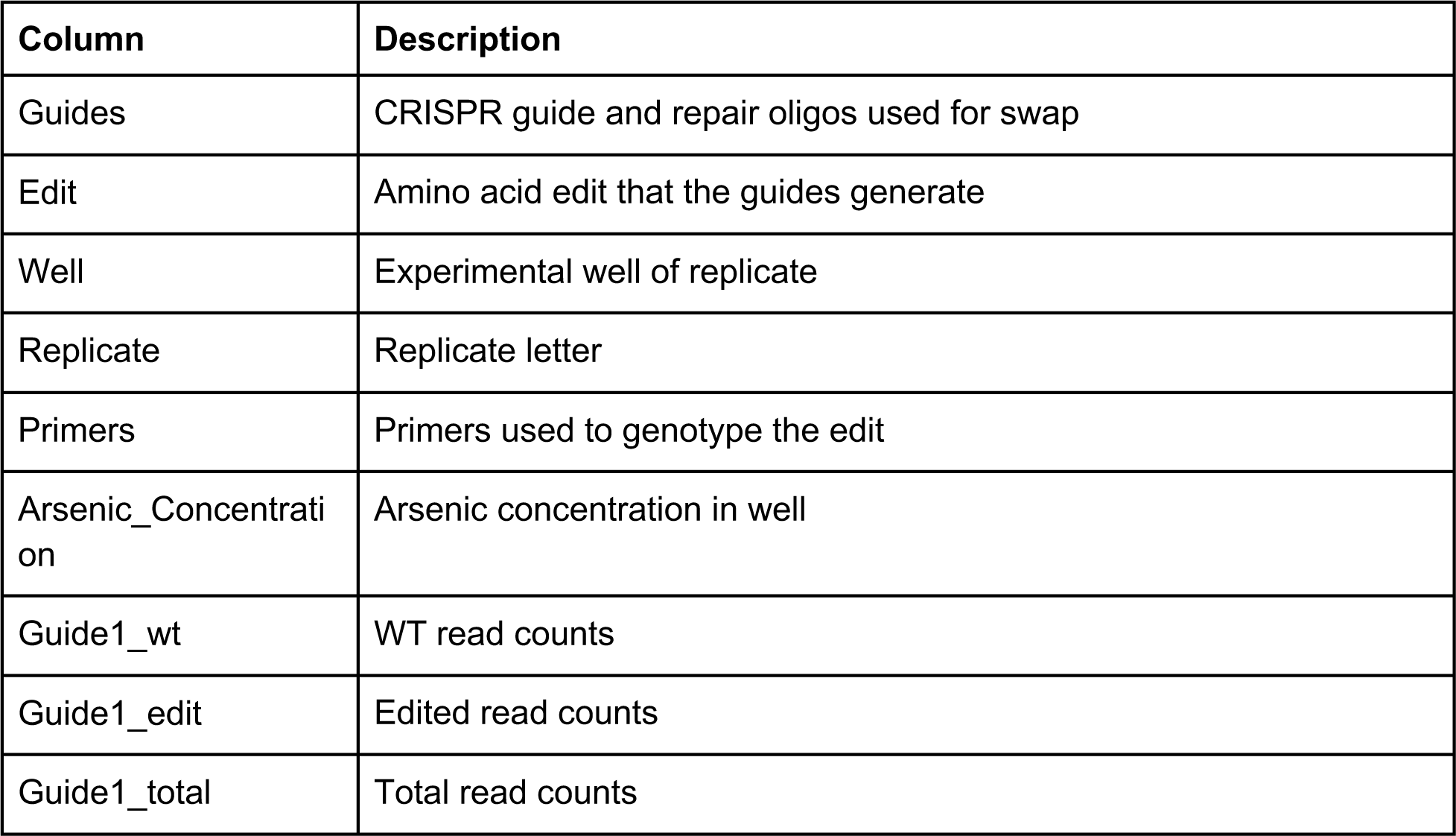
Results from human cell editing experiment.

**Figure 5-source data 2:**
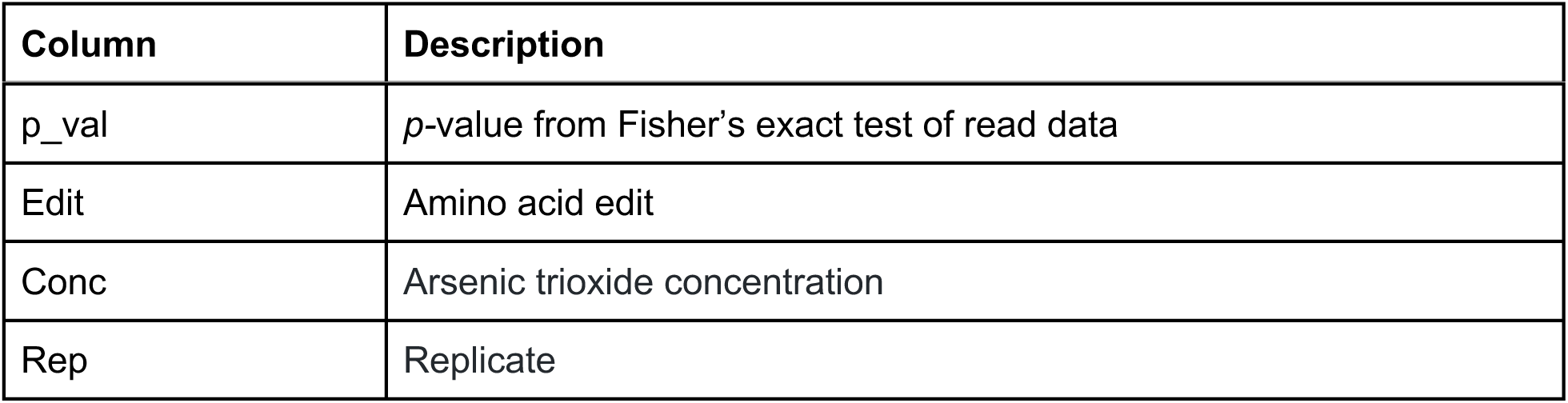
Fisher’s exact test *p*-values for human cell editing experiment

**Figure 5-source data 3:**
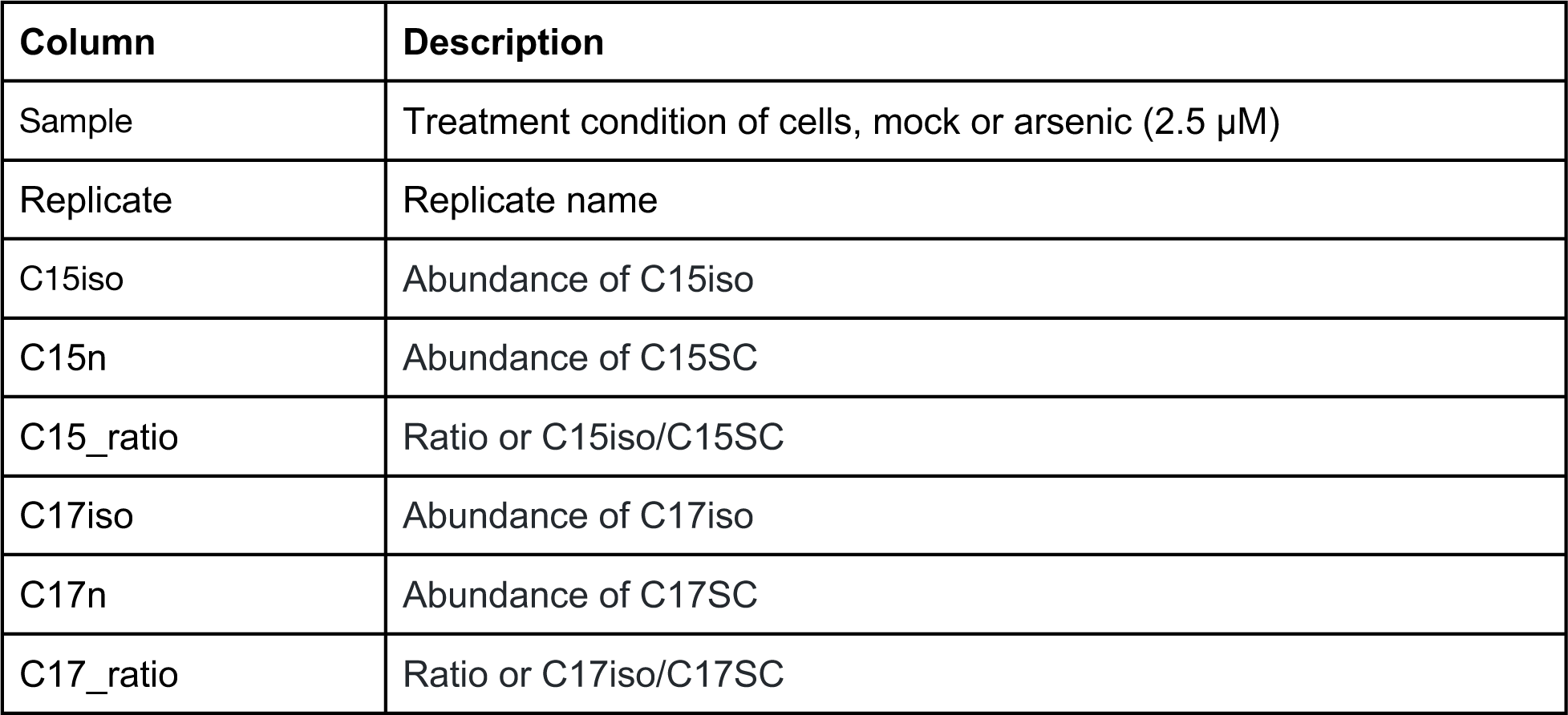
Metabolite measurements for human cell line experiments.

**Figure 2-source data 9:**
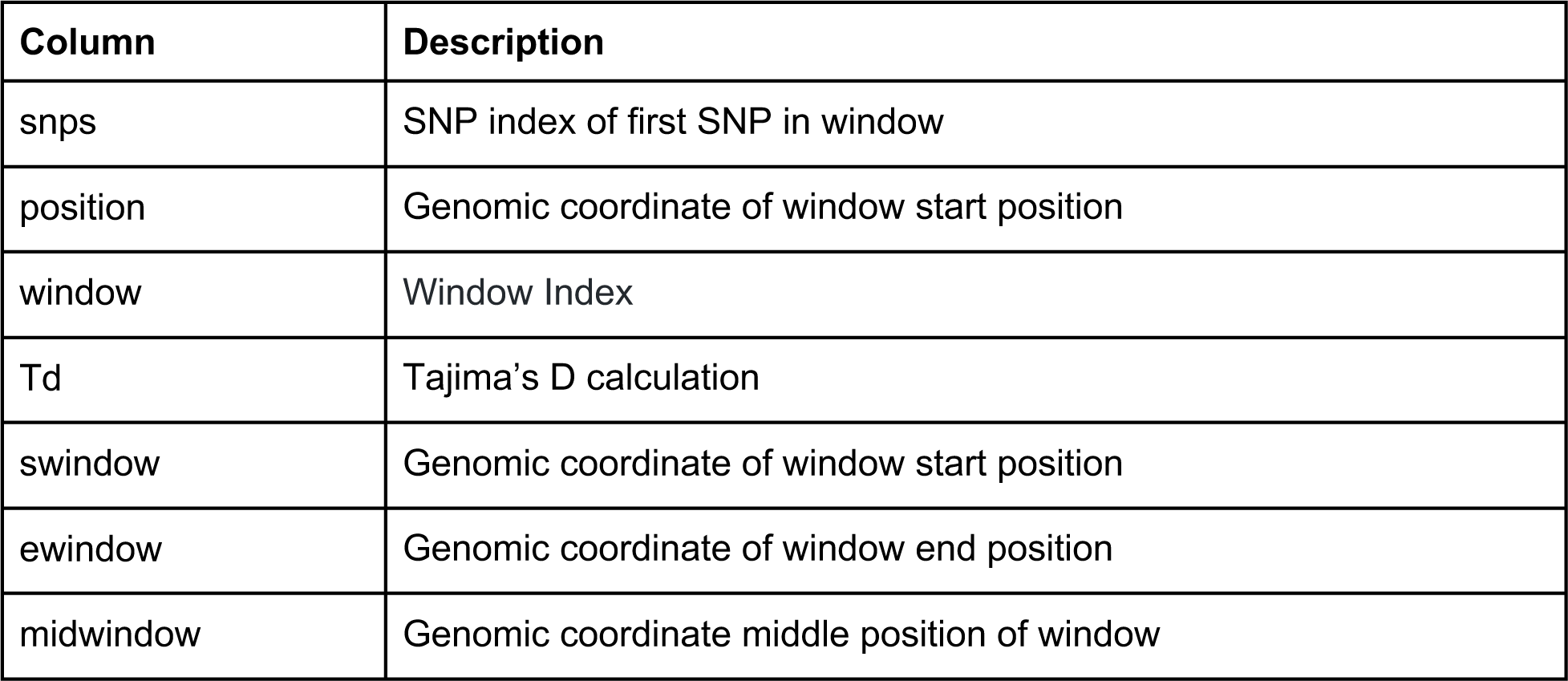
Tajima’s D calculation of GWA QTL

**Figure 2-source data 10:**
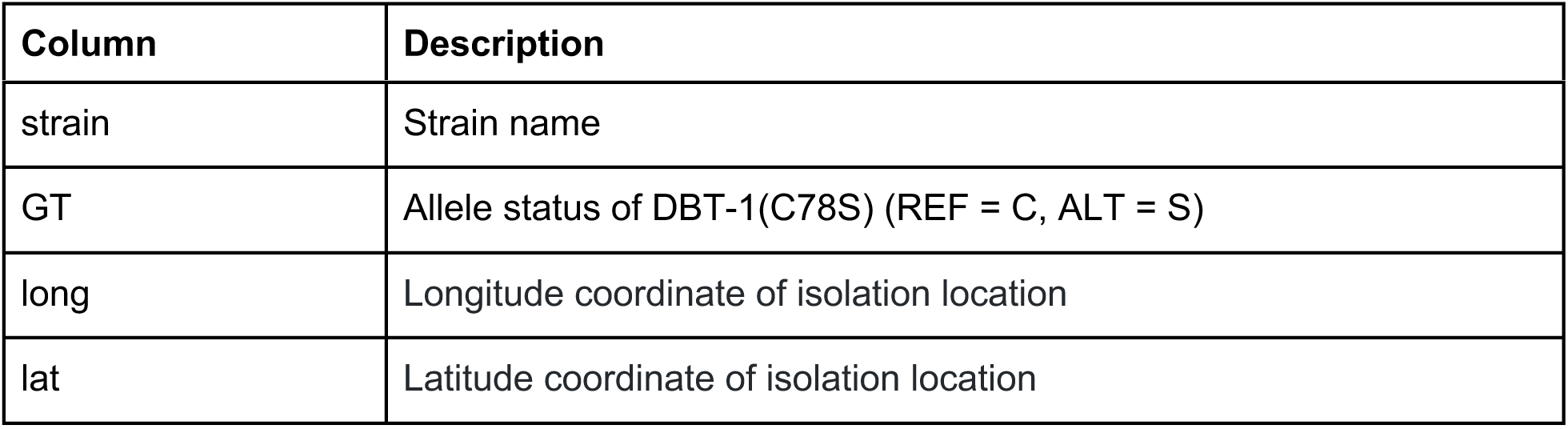
Strain isolation locations

